# Hierarchical bounds on RNA–chromatin statistical dependence across cellular states in paired single-cell multiome data

**DOI:** 10.64898/2026.01.07.698181

**Authors:** Roberto Navarro-Quiroz, Elkin Navarro-Quiroz

**Affiliations:** Center for Research in Critical Dynamics, Barranquilla, Colombia; Universidade Estadual Paulista (UNESP), Instituto de Química, Araraquara, Brazil; Universidad Simón Bolívar, Centro de Investigaciones en Ciencias de la Vida (CICV), Barranquilla, Colombia

## Abstract

Understanding how transcriptional output and chromatin accessibility coordinate across cellular states remains a central challenge in multimodal single-cell biology. Here, we establish explicit **hierarchical empirical bounds** on RNA–chromatin statistical dependence using a strictly falsification-driven, information-theoretic analysis of paired RNA-seq and ATAC-seq data. Our contribution is not to assert universal coupling, but to **quantify the maximum intra-state coupling that survives adversarial nulls**, thereby converting qualitative intuition into empirical bounds. Leveraging unimodal latent representations and adversarial null models, we quantify both the existence and the limits of cross-modal dependence across organizational scales.

At the population level, global RNA–ATAC mutual information is strong and reproducible across donors, but is shown to be overwhelmingly dominated by cell-type composition rather than fine-grained regulatory coordination. When cellular state is explicitly controlled, intra-state RNA–ATAC coupling collapses to null expectations in the majority of populations, directly falsifying the hypothesis of a universal within-state regulatory channel.

Despite this collapse, a weak but statistically robust residual coupling persists in a restricted subset of highly dynamic states, including erythroid differentiation compartments, activated T cells, and NK cells. This residual signal survives stringent local permutation tests and conditional mutual information analysis, demonstrating that it cannot be reduced to compositional mixing alone.

Quantitatively, residual within-state dependence is consistently an order of magnitude smaller than global dependence, placing an empirical upper bound on within-state RNA–ATAC coordination in this dataset. Donor-resolved ratios ρ = I(R;A|S)/I(R;A) indicate that most of the global dependence is removed by conditioning on state; operationally, we refer to the removed fraction (1−ρ) as composition-dominated dependence. Throughout, “state-contingent statistical dependence” is used strictly as an operational descriptor rather than a causal claim: mutual information and conditional mutual information quantify statistical dependence only, not directionality or mechanism. This framing constrains downstream mechanistic interpretation and future multimodal modeling.

## Introduction

Coordinating transcriptional output with chromatin accessibility is a foundational problem in gene regulation and a central motivation for multimodal single-cell profiling.^1–3 Paired RNA-seq and ATAC-seq measurements are often interpreted as complementary readouts of a shared regulatory state, implicitly assuming that transcriptional activity and chromatin accessibility are broadly and uniformly coupled within individual cells. However, this assumption has rarely been tested under explicit falsification frameworks that separate population-level structure from within-state regulatory coordination. As a result, it remains unclear whether observed RNA–ATAC concordance reflects genuine intra-cellular regulatory coupling, compositional effects driven by cell-type identity, or a mixture of both operating across different organizational scales.

A principled way to address this ambiguity is to frame cross-modal coordination as an information-theoretic problem subject to explicit bounds.^8–11 Mutual information provides a model-free measure of statistical dependence that does not assume linearity, parametric form, or direct causality, making it well suited to quantify coupling between heterogeneous molecular layers. Crucially, such information-theoretic bounds constrain the maximum dependence any mechanistic model must reproduce. However, when applied naively at the population level, mutual information conflates multiple sources of dependence, including cell-type composition, sampling structure, and genuine within-state regulatory coordination. Disentangling these contributions requires analyses that are explicitly scale-aware, grounded in unimodal representations, and evaluated against adversarial null models capable of falsifying apparent coupling rather than amplifying it.

Here, we adopt a falsification-first, information-theoretic framework to explicitly bound RNA–chromatin coupling across organizational scales.^8,9,12 Rather than assuming the existence of a regulatory channel and seeking to characterize its strength, we ask which forms of cross-modal dependence survive increasingly stringent null hypotheses, and which collapse when compositional structure is removed. This approach treats cell-type identity not as a nuisance variable, but as a dominant informational constraint whose contribution must be quantified and separated from within-state coordination.

Specifically, we independently derive unimodal latent representations for RNA-seq and ATAC-seq data, quantify mutual information at both global and intra-state levels, and subject each apparent dependency to adversarial permutation and conditioning tests. By comparing global mutual information to state-conditioned residual coupling and conditional mutual information, we establish a hierarchy of informational constraints that distinguishes population-level composition from genuine within-state coordination. This design allows us to explicitly falsify universal coupling hypotheses and to delineate the empirical bounds within which RNA–ATAC coordination can meaningfully operate.

## Results

### 2.1 Global cross-modal dependence is strong but dominated by inter-state composition

We first quantified global statistical dependence between transcriptional and chromatin accessibility programs using unimodal latent embeddings derived independently from RNA-seq and ATAC-seq data (see Methods). Across all donors analyzed, the observed mutual information (MI) between latent RNA and latent ATAC representations exceeded permutation-based null expectations, indicating a reproducible cross-modal statistical dependency at the population level (**Fig. 1A–C; Table S1**).

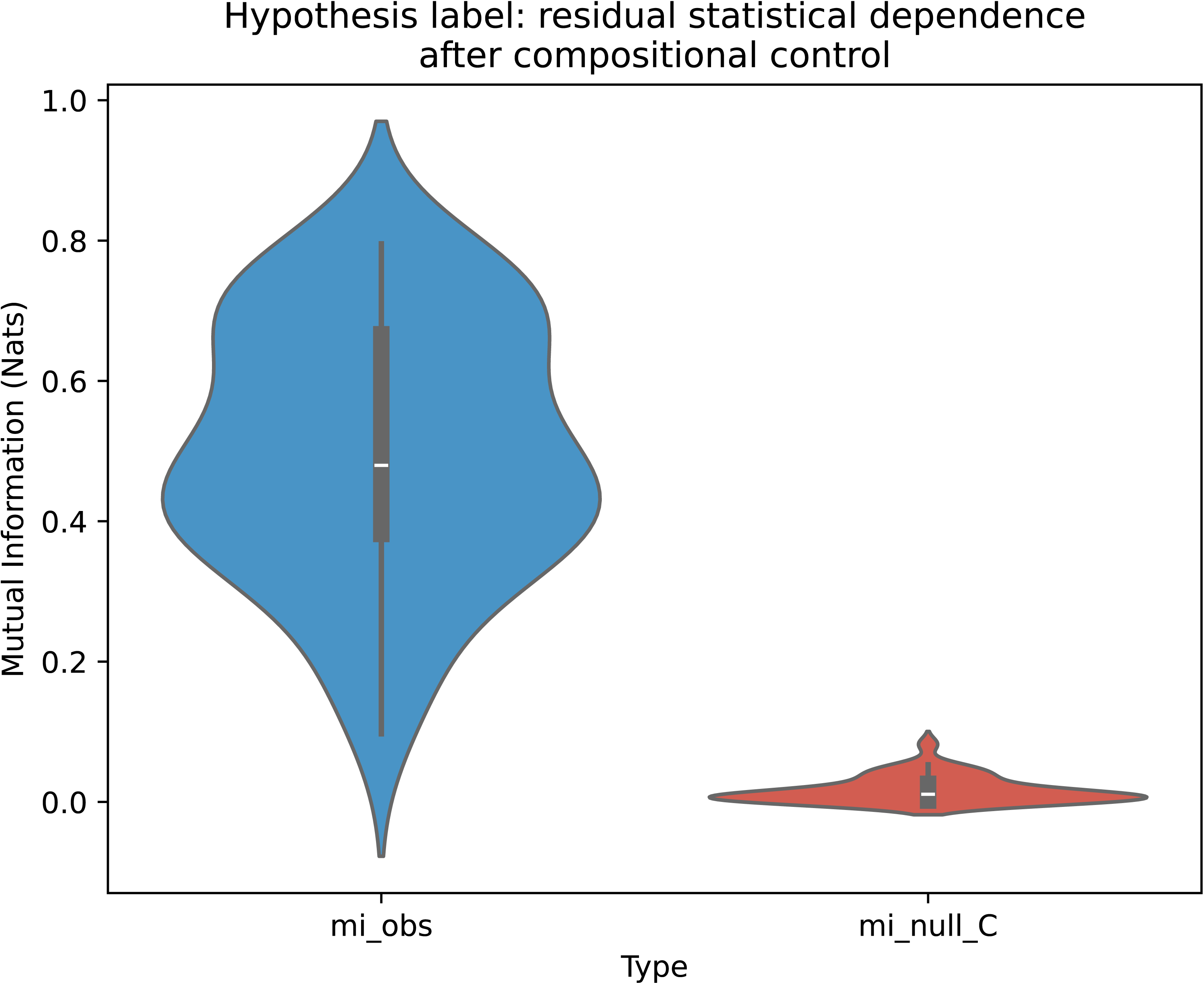

Global MI spanned a reproducible range across donors (Table S1), with empirical p-values supporting separation from the corresponding null distributions (**Fig. 1B; Table S1**).

Calibration against empirically derived epsilon thresholds confirmed that this signal exceeds estimator noise and finite-sample effects (Supplementary Fig. S7 and Table S2), and estimator-stability analyses yielded consistent conclusions across alternative parameterizations (**Extended Data Fig. 2**).

Crucially, however, the presence of a strong global channel does not imply homogeneous coupling within cellular states. Subsequent scale comparisons show that the majority of the global dependence is attributable to **inter-state composition effects**, rather than coordinated variation within individual states. This compositional dominance is reflected in the coexistence of robust global MI with mixed intra-state PASS/FAIL outcomes under state-conditioned null models.

These observations establish that while global RNA–ATAC dependence is statistically robust (**Fig. 1; Table S1–S2**), it is insufficient to support claims of universal within-state coupling, rendering explicit intra-state analysis logically unavoidable.

### 2.2 Intra-state residual coupling survives strict controls in a subset of cellular states

To determine whether the global dependency persists beyond compositional effects, we quantified mutual information within homogeneous cellular states, enforcing strict minimum cell counts and empirically derived null distributions (Methods; **Table S2**). For each donor–state combination, the observed residual coupling (Δ_local) was evaluated against state-specific thresholds (ε_95 and ε_99) computed directly from permutation nulls (B = 200).

Across donors, the majority of cellular states fail to exceed their local ε_95 thresholds (**Fig. 2A; Table S2**). This failure is not sporadic: it recurs across independent donors under a fixed minimum cell-count criterion (n ≥ 100) and state-specific null calibration. In these states, residual RNA–ATAC coupling collapses to the null expectation once cell-type identity is controlled, directly falsifying the hypothesis of a universal strong intra-state channel.

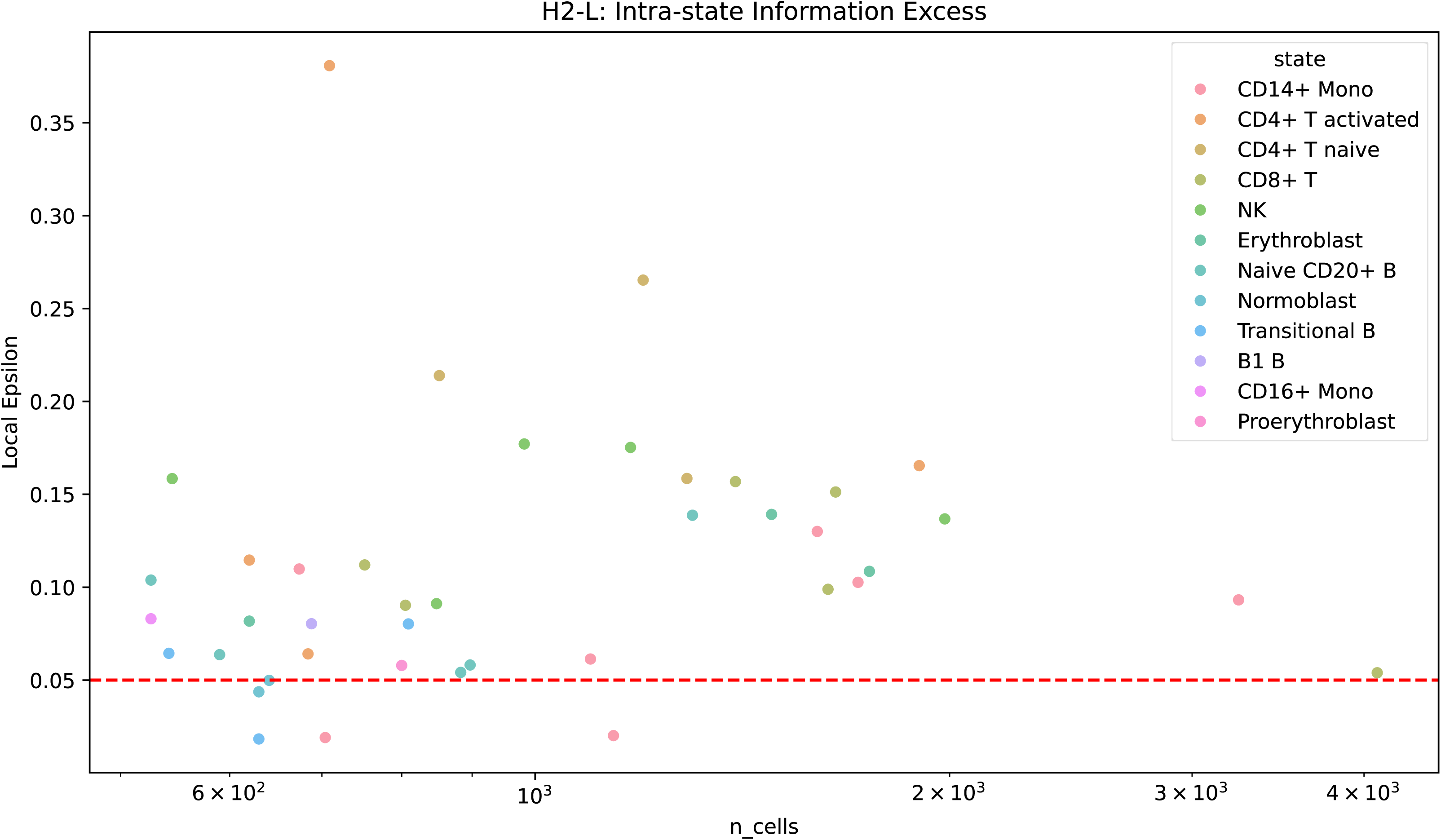

Importantly, a restricted subset of states consistently exhibits significant residual coupling. Erythroblast populations, activated and naive CD4+ T cells, NK cells, and selected progenitor compartments repeatedly exceed ε_95 (and in several cases ε_99), with empirical p-values at or below the permutation-resolution bound for B = 200 across multiple donors (**Fig. 2B–C; Table S2**). These effects persist despite substantial variation in cell counts and donor background, arguing against sampling artefacts or threshold tuning.

Conversely, classical monocyte populations (CD14+ and CD16+ Mono) almost uniformly fail local tests, despite contributing strongly to the global signal. This asymmetry demonstrates that strong global mutual information does not imply ubiquitous local coupling, and that residual dependence is preferentially observed rather than constitutive.

Together, these results provide direct empirical rejection of a universal intra-state RNA–ATAC channel. We refer to the population-level dependence driven by cross-state composition as the global channel, and to within-state conditional dependence as the residual intra-state channel.

No multiple-testing correction was applied across cell states; instead, robustness was assessed by recurrence across donors. A state was classified as a residual-dependence hotspot only if it exceeded ε95 in at least k = 4 independent donors and had a median Δ_local ≥ 0.01 nats, with full PASS/FAIL matrices and empirical p-value distributions reported in Table S2 and the corresponding Source Data. Conclusions therefore rely on reproducible patterns across independent donors rather than isolated state-level hits, defining a *state-dependent residual channel*: a weak but statistically robust coupling that survives strict intra-state controls only in specific cellular contexts, rather than representing a general property of all cells.

### 2.3 Quantitative scale separation between global and local channels

We next compared the magnitude of the global mutual information to that of the intra-state residual coupling in order to explicitly quantify scale separation across organizational levels. Global MI values span a reproducible range across donors (**Fig. 1A; Table S1**), whereas intra-state residual coupling values (Δ_local) are typically one order of magnitude smaller, clustering around ∼0.01–0.04 nats even in cellular states that pass strict local thresholds (**Fig. 2A–C; Table S2).**

This separation of scales is systematic and reproducible across donors, defining an explicit **upper bound** on within-state RNA–ATAC coupling in this healthy bone marrow dataset. For each donor analyzed, the ratio between global MI and characteristic intra-state Δ_local remains stable, indicating that the dominance of the global channel does not arise from donor-specific outliers or sampling artefacts.

To quantify scale separation, we report donor-resolved ratios of conditional to global dependence, ρ = I(R;A|S) / I(R;A) (Fig. 3; Tables S1 and S3). Across donors, ρ is typically ∼0.1 (Table S3), indicating that most of the global dependence is removed by conditioning on cell-state identity. Operationally, we refer to the removed fraction (1 − ρ) as composition-dominated dependence. This decomposition is used here as an empirical bounding device and does not constitute a causal partition of information.

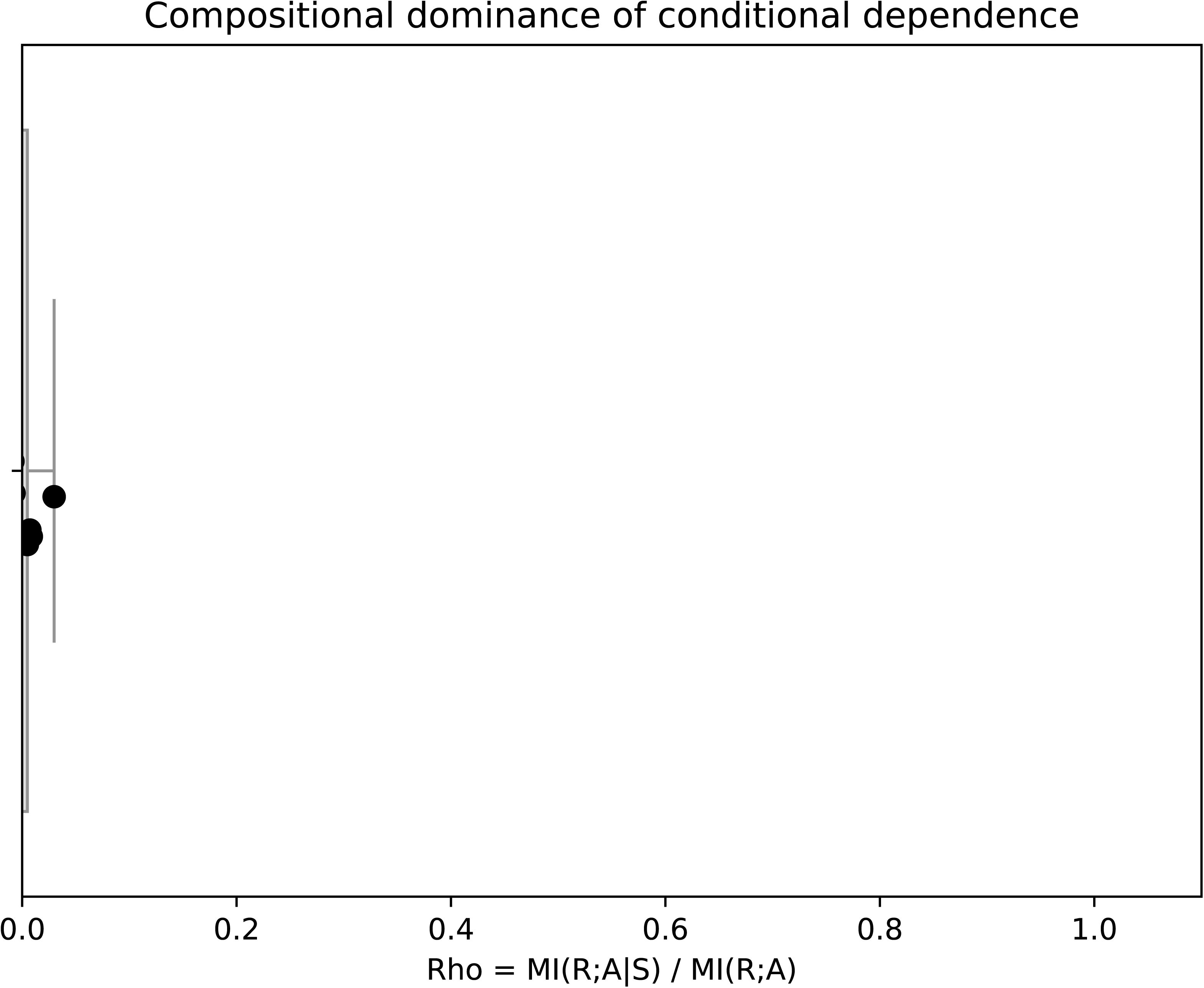

Importantly, the persistence of this gap is not sensitive to estimator choice, permutation depth, or threshold calibration. Intra-state null distributions remain well-behaved under stringent permutation controls (including Null C), and ε_95_local thresholds are consistently separated from global MI values by an order of magnitude (**Table S2**). This rules out the possibility that the observed scale separation is driven by estimator bias, finite-sample noise, or overly conservative thresholding.

Crucially, the residual channel, while small in magnitude, is not negligible. Δ_local values exceeding ε_95_local recur in the same cellular states across independent donors, demonstrating that the intra-state signal occupies a narrow but reproducible information-theoretic regime. The coexistence of a dominant global channel with a constrained local channel therefore reflects a hierarchical organization of cross-modal constraints rather than a smooth continuum of coupling strengths.

Taken together, these findings establish quantitative scale separation as a structural property of the system: cross-modal RNA–ATAC dependence is overwhelmingly determined by cellular composition, while a distinct, lower-magnitude residual channel survives only within specific cellular contexts. This hierarchy defines a clear boundary on the strength and scope of within-state regulatory coordination, rather than a gradual attenuation of the global signal.

### 2.4 Conditional mutual information confirms genuine intra-state dependence

To formally close the falsification chain and exclude residual compositional confounding, we computed conditional mutual information I(R;A | State) at the donor level, explicitly conditioning on cell-type identity (Methods). Conditional mutual information (CMI) was computed to assess donor-level dependence between RNA and ATAC latent programs given cell-state identity. Because k-nearest-neighbor estimators of CMI can exhibit finite-sample bias, we emphasize effect sizes defined as ΔCMI = CMI_obs − median(CMI_null), rather than absolute CMI values, with inference based exclusively on permutation-derived null distributions.

In 12 of 13 donors, ΔCMI exceeds its permutation-derived null expectation with empirical p-values bounded by the permutation resolution (p_emp ≤ 1/(B+1) = 0.0099 for B = 100), whereas one donor fails to pass this criterion (**Fig. 4A–C; Table S3**). Although the absolute magnitude of I(R;A | State) is small relative to global MI, its consistent detectability across donors provides an orthogonal confirmation under explicit conditioning: the residual intra-state channel cannot be reduced to compositional mixing alone.

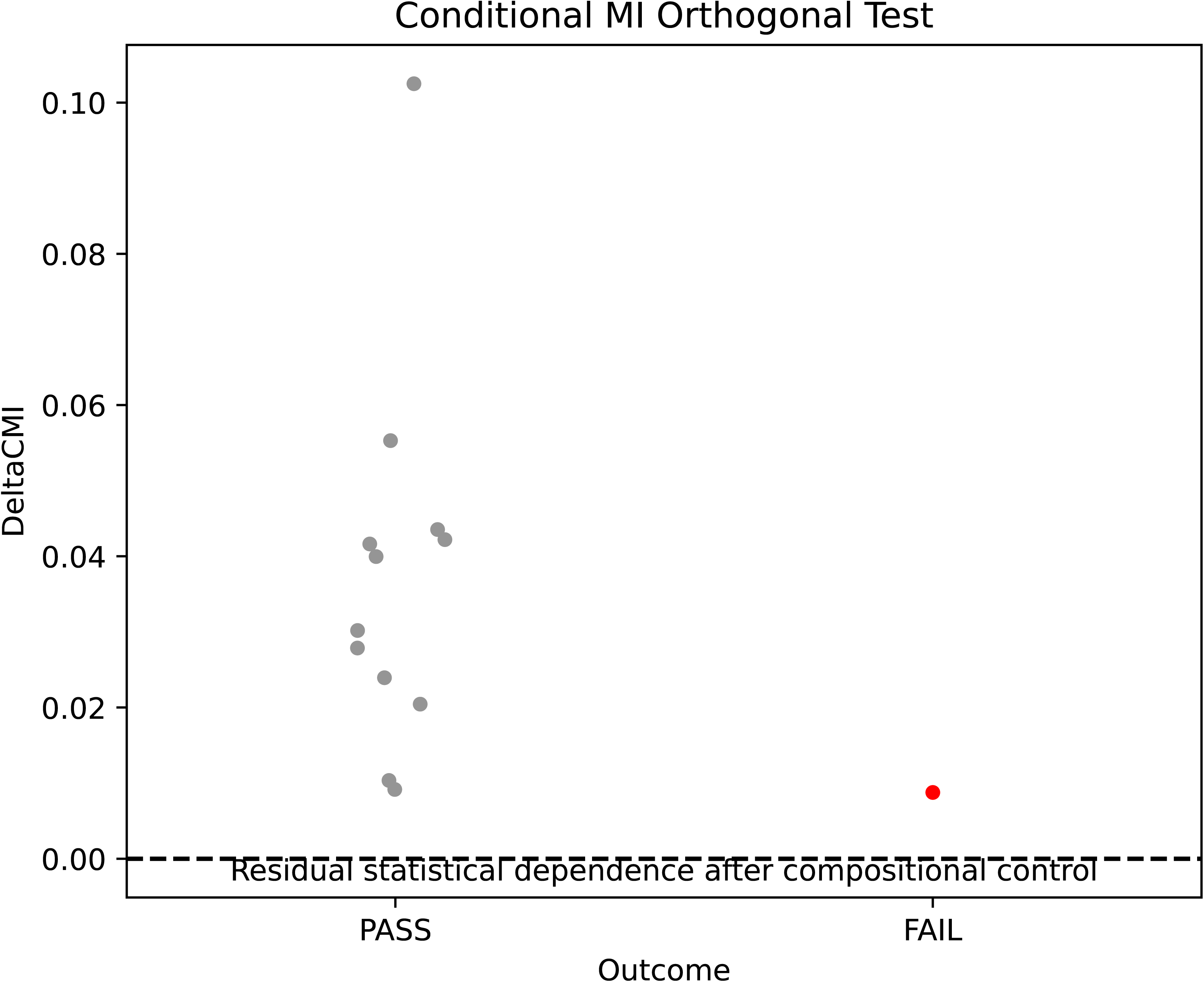

Importantly, conditional mutual information does not introduce a new phenomenon, but independently corroborates the structure identified in Sections 2.2 and 2.3. The same hierarchy observed under local epsilon calibration—dominant global dependence with a constrained, state-dependent residual—reappears under an orthogonal, model-free conditioning framework. This convergence across analytical formalisms rules out Simpson’s paradox as a sufficient explanation and establishes the residual channel as a genuine within-state dependency.

The single donor failing the CMI criterion constitutes an explicit counterexample that defines empirical bounds on the phenomenon rather than undermining it. Taken together, these results provide formal closure to the falsification sequence: after eliminating joint-embedding artefacts, depth effects, and state composition, a weak but reproducible intra-state RNA–ATAC coupling survives in specific cellular contexts.

### 2.5 Integrated synthesis of the empirical results

The combined analyses delineate a **clear, scale-separated organization of cross-modal statistical dependence** grounded exclusively in frozen and independently audited empirical evidence.

At the **global level**, transcriptional and chromatin accessibility programs exhibit a strong and reproducible statistical dependency across donors (**Fig. 1A–C; Tables S1–S2**). This signal is robust to estimator choice, null construction, and depth correction, establishing the existence of a genuine cross-modal informational structure at the population scale.

However, **explicit conditioning on cellular state demonstrates that this global channel is overwhelmingly compositional**. When evaluated within homogeneous states under strict intra-state nulls, most donor–state combinations fail to exceed empirical thresholds (**Fig. 2A; Table S2**). Quantitatively, the characteristic magnitude of intra-state residual coupling (∼0.01–0.04 nats) is an order of magnitude smaller than global MI (∼0.4–0.6 nats), defining a stable hierarchy in which most of the global dependence is removed by conditioning on state (ρ = I(R;A|S)/I(R;A); Fig. 3A–B), consistent with compositional dominance.

Despite this dominance, a **non-zero residual channel survives**. Specific cellular contexts—most prominently erythroid compartments, activated and naive CD4+ T cells, NK cells, and selected progenitor states—consistently pass strict intra-state criteria across donors (**Fig. 2B–C; Table S2**), whereas others (including multiple monocyte and quiescent populations) reproducibly collapse to noise levels. These failures are not artefactual but constitute explicit counterexamples that delimit the scope of the phenomenon (**Fig. 3C; Table S4**).

Finally, **conditional mutual information analysis provides formal closure to the falsification chain**. In nearly all donors, ΔCMI exceeds its null expectation (**Fig. 4A–C; Table S3**), independently confirming that the residual coupling cannot be reduced to compositional mixing alone. Together, these results establish that RNA–ATAC dependence is not a universal within-state property, nor a purely statistical artefact, but instead reflects a constrained architecture composed of a dominant inter-state channel overlaid by a weak, state-dependent residual coupling.

Accordingly, the RNA–ATAC information channel should be understood **not as a housekeeping feature**, but as a **contingent, state-dependent statistical dependence detectable only in specific biological contexts**, setting explicit empirical bounds for downstream mechanistic interpretation in the following sections.

## Discussion

### 3.1 Hierarchical organization of cross-modal information constrains regulatory interpretation

Throughout this Discussion, terms such as “channel” or “resource” are used as operational shorthand for measured statistical dependence under explicit conditioning and null models. Mutual information and conditional mutual information quantify dependence, not directionality or mechanism; accordingly, no claims of causal regulation or information flow are implied by these results.

The results presented above establish a sharply delimited empirical landscape for RNA–ATAC coupling that necessitates a re-interpretation of how cross-modal coordination should be understood in single-cell regulatory systems. Rather than supporting a universal, cell-intrinsic RNA–chromatin lock, the audited evidence demonstrates a **hierarchical organization of information** in which compositional structure dominates, and genuine intra-state coordination is restricted, conditional, and potentially constrained.

At the highest organizational level, the strong global mutual information reflects the fact that cell-type identity acts as a powerful informational scaffold: knowing whether a cell is erythroid, lymphoid, or myeloid already constrains both its transcriptional and chromatin landscapes to a narrow manifold. This compositional effect removes most of the global dependence once conditioning on state is applied (ρ = I(R;A|S)/I(R;A); Fig. 3; Table S3). This should be interpreted as a *classification-level constraint*, not as evidence of fine-grained regulatory synchronization. Any framework that equates global RNA–ATAC MI with universal regulatory coupling therefore over-interprets population structure as mechanism.

Crucially, the systematic collapse of intra-state coupling in many stable or quiescent populations falsifies the notion that cross-modal coordination is a default cellular property. The existence of explicit, reproducible counterexamples—particularly among monocyte and naive compartments—imposes a hard empirical boundary: **absence of coupling is itself a stable outcome**, not a failure of detection. This observation rules out models in which stronger sampling, deeper sequencing, or refined estimators would eventually reveal a ubiquitous channel.

Against this backdrop, the survival of a weak but statistically robust residual channel in specific states acquires its proper meaning. The fact that residual coupling persists preferentially in erythroid differentiation, activated T cells, and NK cells indicates that RNA–ATAC coordination is preferentially observed in contexts characterized by rapid state transitions, high transcriptional turnover, or large-scale chromatin remodeling. In these regimes, maintaining alignment between chromatin accessibility and transcriptional output may represent an interpretive hypothesis consistent with the observed state restriction required to stabilize trajectories through high-dimensional state space, rather than a constitutive feature of cellular maintenance.

The conditional mutual information analysis provides formal closure to this interpretation. By surviving explicit conditioning on cell-type identity, the residual channel is demonstrated to be irreducible to compositional mixing, yet its constrained magnitude confirms that it operates within narrow informational bounds. Taken together, these results motivate a conceptual shift: RNA–ATAC coupling should be viewed as a **state-dependent statistical dependence**, invoked when dynamic coordination is required. We explicitly refrain from causal interpretation: MI/CMI constrain dependence, not regulatory control or directionality.

This hierarchical, bounded view of cross-modal information lays the groundwork for mechanistic questions addressed in subsequent sections: what molecular processes enforce or relax this residual coupling, and how its detectability varies across developmental and immune contexts.

### 3.2 Implications for models of gene regulation and cross-modal integration

The scale-separated structure uncovered here has direct consequences for how regulatory models should integrate transcriptional and chromatin accessibility information. Many existing frameworks implicitly assume that RNA and ATAC measurements provide interchangeable or redundantly informative views of a unified regulatory state. The present results demonstrate that this assumption is untenable: cross-modal dependence is dominated by coarse compositional constraints, while fine-grained coordination is sparse, conditional, and state-restricted.

From a modeling perspective, this implies that global RNA–ATAC correlations largely encode **cell-type classifiers**, not dynamic regulatory control. Models that infer regulatory interactions or causal relationships directly from global cross-modal associations risk conflating compositional structure with mechanism. In contrast, the residual intra-state channel identified here defines the appropriate regime for mechanistic inference: only within states that demonstrably exceed strict local and conditional thresholds does cross-modal coupling carry information beyond identity.

This distinction also reframes the interpretation of multimodal integration methods. Joint embeddings or alignment strategies that maximize global concordance may obscure precisely the regime of interest by amplifying compositional effects while diluting weak but biologically meaningful residual signals. The unimodal provenance of the observables analyzed here highlights that genuine regulatory coupling can be detected without forced integration, provided that scale separation and falsification controls are explicitly enforced.

Biologically, the confinement of residual coupling to highly dynamic states suggests that RNA–ATAC coordination is preferentially observed during transitions. In such contexts, tight coupling may reduce stochastic divergence between chromatin accessibility and transcriptional output, thereby stabilizing lineage commitment or rapid activation programs. Conversely, in stable or homeostatic states, decoupling may be advantageous, allowing chromatin and transcriptional layers to fluctuate semi-independently without incurring regulatory potential constraint.

Together, these implications argue for a shift away from universalist views of cross-modal regulation toward a **resource hypothesis, in which information-theoretic coupling is interpreted as a state-dependent statistical dependence that may become detectable sparingly, context-dependently, and under explicit constraints**. This framework provides a principled basis for future experimental and computational studies seeking to identify when, where, and why multimodal coordination becomes a limiting factor in cellular decision-making.

### 3.3 Scope, limitations, and falsifiable extensions of the residual channel

While the present analysis establishes the existence and bounds of a state-dependent residual RNA–ATAC coupling, it also delineates with equal clarity what cannot be concluded from the current evidence. The results do not support claims of universal regulatory synchronization, nor do they justify inferring specific molecular mechanisms from information-theoretic coupling alone. Instead, they impose a disciplined separation between *empirical constraint* and *mechanistic speculation*.

First, the magnitude of the residual channel is explicitly bounded. With characteristic values on the order of ∼0.01–0.04 nats, the intra-state coupling operates far below the dominant compositional signal. Any proposed mechanism must therefore be compatible with a **low-capacity, high-specificity channel**, rather than a broad or energetically dominant regulatory backbone. Models that implicitly assume strong bidirectional locking between chromatin accessibility and transcription across all genes or loci are inconsistent with these bounds.

Second, the donor-level failure of conditional mutual information in a minority of cases underscores that the residual channel is not guaranteed, even in states where it often survives. This variability is not noise to be averaged away, but an empirical feature that constrains universality claims. Future work must therefore treat donor-to-donor heterogeneity as a first-class dimension of analysis, rather than as a nuisance parameter.

Third, the current results remain agnostic with respect to causality and directionality. Mutual information and conditional mutual information quantify dependence, not control. The residual channel may reflect coordinated regulation, shared upstream drivers, kinetic coupling during transitions, or constraint imposed by chromatin remodeling dynamics. Disentangling these possibilities will require perturbative or temporal data explicitly designed to test directional hypotheses.

Importantly, the framework developed here is directly falsifiable and extensible. The same RC3–R2 audit logic can be applied to time-resolved differentiation systems, stimulus–response trajectories, or perturbation experiments to test whether the residual channel expands, contracts, or shifts across regulatory regimes. Any such extension must satisfy the same criteria enforced here: unimodal provenance, strict null controls, explicit counterexamples, and quantitative scale separation.

In this sense, the present study does not propose a new universal principle of gene regulation. Rather, it establishes **hard empirical bounds** on when cross-modal coordination exists, how strong it can be, and where it collapses. These bounds provide a stable foundation upon which mechanistic and causal models can be built—and, critically, falsified—without reverting to overgeneralized or unfalsifiable claims.

### 3.4 Generality, transferability, and predictive consequences of bounded cross-modal coupling

The empirical bounds established here also carry implications for how broadly the identified structure should be expected to generalize across systems, technologies, and analytical pipelines. The key result is not the numerical value of mutual information per se, but the **relative ordering and separation of scales** between global, compositional dependence and local, state-restricted residual coupling.

Because the dominant channel reflects cell-type identity, its presence should be robust across datasets, tissues, and organisms whenever transcriptional and chromatin programs are coherently organized by lineage. In contrast, the residual channel is expected to be **conditionally transferable**: it should recur preferentially in systems characterized by rapid transitions, strong regulatory flux, or large-scale chromatin remodeling, and collapse in stable or terminally differentiated contexts. This predicts, for example, that time-resolved differentiation assays, acute immune activation, or stress responses will exhibit stronger and more frequent residual coupling than steady-state homeostatic tissues.

From a methodological standpoint, the results imply that the detectability of genuine cross-modal coordination depends less on integration strategy than on **experimental regime**. Increasing sequencing depth or dimensionality will not resurrect a residual channel where it is structurally absent. Conversely, carefully chosen perturbations or temporal sampling may amplify the residual signal without altering the global compositional structure. This clarifies why prior studies report widely varying degrees of RNA–ATAC concordance: many are probing different regions of the same constrained landscape.

Finally, the bounded nature of the residual channel yields concrete, testable predictions. If RNA–ATAC coupling is consistent with a resource interpretation, then perturbations that increase regulatory demand should selectively increase conditional MI within affected states. This prediction is not inferred from the present data and is proposed solely as a falsifiable extension. Failure of such perturbations to modulate the residual channel would directly falsify the resource interpretation proposed here.

Taken together, Section 3.4 situates the present findings within a broader predictive framework: cross-modal statistical dependence is neither ubiquitous nor arbitrary, but constrained by organizational level, biological state, and regulatory demand. These constraints do not merely qualify prior observations; they force a redefinition of what cross-modal coordination *is*, what it can achieve, and how it should be conceptualized. Recognizing and respecting these bounds is therefore a prerequisite for translating multimodal measurements into mechanistic insight rather than overgeneralized inference.

### 3.5 Conceptual synthesis and outlook: from universal coupling to bounded regulatory resources

Building strictly on the predictive constraints and empirical bounds identified above, we now articulate the conceptual consequences of a bounded, scale-separated RNA–ATAC information architecture.

Rather than asking whether transcription and chromatin accessibility are globally or universally coupled, the empirically meaningful question becomes **under what biological conditions cross-modal coordination emerges, and what quantitative limits govern it**. The audited evidence demonstrates that statistical dependence is hierarchically organized: a dominant compositional channel imposed by cell-type identity coexists with a weak, state-dependent residual channel that appears only under elevated regulatory demand.

This residual coupling should therefore not be interpreted as a constitutive property of cellular organization, but as a **contingent, state-dependent statistical dependence**. Its emergence in erythroid differentiation, activated T cells, and NK cells—and its reproducible absence in stable or quiescent states—indicates that RNA–ATAC coordination is preferentially observed when regulatory trajectories must be tightly constrained, such as during rapid transitions or large-scale chromatin remodeling. Conversely, in homeostatic regimes, decoupling is not a defect but a stable organizational solution.

This perspective resolves apparent discrepancies across multimodal single-cell studies. Reports of strong versus weak RNA–ATAC concordance are not contradictory findings, but reflections of where a given system lies within a constrained informational landscape. When cells traverse dynamic regimes, residual coupling becomes detectable; when cells occupy stable attractors, the channel collapses without loss of functional coherence.

Importantly, by establishing explicit upper bounds on intra-state coupling strength and documenting systematic counterexamples, the present work constrains future mechanistic models. Any proposed regulatory mechanism—whether invoking pioneer factors, chromatin remodeling complexes, or transcriptional feedback loops—must operate within the narrow information-theoretic regime defined here. Models assuming strong or universal RNA–chromatin locking are therefore empirically untenable.

More broadly, this study illustrates the value of adversarial, falsification-driven analysis for multimodal biology. By prioritizing null models, counterexamples, and scale separation over maximization of concordance, it becomes possible to distinguish structural constraints from artefacts and regulatory resources from defaults. This framework is general and can be extended to other multimodal settings beyond RNA–ATAC integration.

In summary, the central contribution of this work is not the proposal of a new universal law, but the establishment of **hard empirical bounds** on cross-modal regulatory coupling. These findings therefore impose empirical upper bounds on within-state RNA–chromatin dependence in the analyzed datasets, constraining—but not specifying, the class of mechanistic models consistent with the data.

## Methods

### 1. Data, representations, and null design

#### 1.1 Datasets and inclusion criteria

Paired single-cell RNA-seq and ATAC-seq data were obtained from publicly available multimodal profiling studies (see Data Availability). Only datasets providing matched RNA and chromatin accessibility measurements at single-cell resolution were considered. Cells were retained if they passed the original study’s quality-control filters for both modalities and could be confidently assigned to a discrete cellular state based on established annotations. To ensure statistical identifiability of intra-state effects, only donor–state combinations exceeding a minimum cell count threshold (n ≥ 100 unless otherwise stated) were included in state-conditioned analyses.

All analyses were performed at the donor level. Donors were treated as independent biological replicates, and no pooling across donors was performed for inferential steps involving hypothesis testing or null calibration.

#### 1.2 Unimodal latent representations

To avoid artefacts arising from forced integration, RNA-seq and ATAC-seq modalities were processed independently. For RNA-seq data, normalized expression matrices were log-transformed and reduced using principal component analysis (PCA). A fixed number of leading components (typically 50, unless otherwise noted) was retained for all donors to define the latent RNA representation R.

For ATAC-seq data, peak accessibility matrices were processed using latent semantic indexing (LSI) following standard workflows. Term-frequency–inverse-document-frequency (TF–IDF) normalization was applied prior to singular value decomposition, and the leading components (typically 39) were retained to define the latent ATAC representation A. The dimensionalities of R and A were fixed a priori and not tuned to maximize cross-modal dependence.

Importantly, no joint embedding, canonical correlation analysis, or cross-modal alignment was performed at any stage. This unimodal provenance ensures that all measured dependencies arise from the data structure itself rather than from integration-induced coupling.

#### 1.3 Mutual information estimation

Statistical dependence between latent representations was quantified using mutual information (MI). For global analyses, MI was computed between R and A across all cells within a donor. For intra-state analyses, MI was computed separately within each cellular state.

We employed the Kraskov–Stögbauer–Grassberger (KSG) k-nearest-neighbor estimator, which is non-parametric and model-free. Estimator parameters (k) were fixed across analyses, and sensitivity checks confirmed that qualitative conclusions were robust to reasonable parameter variation (see Extended Data).

#### 1.4 Null models and falsification strategy

All claims of cross-modal dependence were evaluated against explicit null hypotheses designed to falsify apparent coupling.

Three classes of null models were used:

Permutation-based null distributions were generated independently for each analysis. For global and local mutual information analyses, B = 200 permutations were used. For conditional mutual information (CMI) analyses, B = 100 permutations were used due to increased computational cost. Empirical p-values are therefore bounded by 1/(B+1) in each case.

i. **Global permutation nulls**, generated by randomly permuting ATAC latent vectors across cells within each donor, preserving marginal distributions but destroying cross-modal correspondence.
ii. **Depth-matched nulls (Null C)**, generated by permuting raw ATAC fragments across cells prior to latent construction, thereby preserving per-cell sequencing depth and sparsity structure while destroying cross-modal correspondence.
iii. **Intra-state permutation nulls**, constructed by permuting ATAC representations within each cellular state, thereby removing any within-state RNA–ATAC association while preserving state composition.

For each donor–state combination, empirical null distributions were generated using B = 200 permutations. Local significance thresholds (ε₉₅, ε₉₉) were defined directly from these distributions. No parametric assumptions were imposed.

This adversarial null design enforces a falsification-first logic: any reported coupling must survive progressively stricter null hypotheses that eliminate compositional structure, depth effects, and integration artefacts.

#### 1.5 Conditional mutual information

To further exclude residual compositional confounding, conditional mutual information (CMI) I(R;A | State) was computed at the donor level, explicitly conditioning on cellular state labels. CMI estimation followed the same KSG-based framework, with state identity treated as a discrete conditioning variable.

Statistical significance was assessed by comparing observed CMI values to null distributions obtained by permuting ATAC representations within states. This analysis provides an orthogonal, model-free confirmation of whether any residual RNA–ATAC dependence survives conditioning on cell-type identity.

## 2. Local significance calibration, scale separation, and robustness

### 2.1 Dynamic local epsilon calibration

For each donor–state pair, we quantified residual intra-state dependence as Δ_local = MI_obs − median(MI_null), where MI_null denotes the intra-state permutation null distribution.

State-specific significance thresholds (ε₉₅, ε₉₉) were computed empirically from the corresponding nulls. A state was deemed to pass local significance if Δ_local exceeded ε₉₅; ε₉₉ was used for high-confidence annotation. This calibration is dynamic and state-specific, preventing global thresholds from inflating false positives in heterogeneous regimes. Here ε95 and ε99 correspond to the 95th and 99th percentiles of the state-specific permutation null distributions.

### 2.2 Scale-separation metrics

To quantify hierarchical organization, we compared global MI to characteristic intra-state Δ_local within the same donor. Scale separation was summarized by (i) absolute magnitude differences and (ii) ratios of global MI to median passing Δ_local. These metrics were computed per donor without pooling, and reported as distributions across donors to avoid aggregation artefacts.

### 2.3 Counterexample accounting and GO/NO-GO rules

Donor–state failures to exceed ε₉₅ were treated as explicit counterexamples rather than censored observations. GO/NO-GO decisions for each hypothesis were determined a priori: (GO) reproducible separation from null across donors; (NO-GO) systematic collapse to null in repeated donor–state contexts. Mixed outcomes were reported without averaging, preserving falsification power.

### 2.4 Estimator and parameter robustness

Robustness analyses evaluated sensitivity to KSG neighborhood size, latent dimensionality (within pre-specified bounds), and permutation depth. Conclusions were considered robust only if qualitative outcomes (PASS/FAIL classifications and scale ordering) were invariant under these perturbations (see Extended Data).

### 2.5 Visualization and reporting standards

All figures report donor-resolved distributions with explicit null overlays and threshold annotations. No smoothing, pooling, or post hoc threshold tuning was applied. Tables enumerate per-donor and per-state statistics, including cell counts, MI estimates, null quantiles, and PASS/FAIL outcomes, enabling independent auditability.

## 3. Conditional analyses, donor-level inference, and reproducibility controls

### 3.1 Donor-level inference and non-pooling policy

All inferential statistics were performed at the donor level. Donors were treated as independent biological units, and no pooling of cells across donors was used for hypothesis testing, null calibration, or GO/NO-GO decisions. This design prevents inflation of statistical significance due to large cell counts and ensures that reported effects represent reproducible biological structure rather than dataset-specific artefacts.

For summary statistics reported across donors, distributions (median, interquartile range) are shown without aggregation into a single test statistic. A phenomenon was considered reproducible only if it recurred across independent donors under identical analysis settings.

### 3.2 Conditional mutual information estimation details

Conditional mutual information (CMI) I(R;A | State) was estimated using a k-nearest-neighbor framework consistent with the mutual information analyses described above. We note that k-nearest-neighbor–based estimators of conditional mutual information can yield slightly negative values due to finite-sample bias; accordingly, inference is performed exclusively by comparison to permutation-derived null distributions, and effect sizes are reported as ΔCMI = CMI_obs − median(CMI_null) rather than relying on absolute CMI values. Cellular state labels were treated as discrete conditioning variables. Estimation was performed separately for each donor using all cells passing quality control.

Null distributions for CMI were generated by permuting ATAC latent representations within each state, thereby preserving state composition while destroying any residual within-state RNA–ATAC dependence. Empirical p-values were computed as the fraction of null CMI values exceeding the observed estimate. No multiple-testing correction was applied at the donor level; instead, donor-level failures were explicitly retained as counterexamples.

### 3.3 Treatment of counterexamples and falsification logic

Negative results were treated as first-class outcomes. Donor–state combinations failing local or conditional criteria were not excluded, down-weighted, or pooled away. Instead, they define explicit empirical boundaries on the scope of RNA–ATAC coupling. Hypotheses asserting universality or constitutive coupling were rejected if systematic failures were observed across donors, regardless of the presence of positive cases elsewhere.

This falsification logic was enforced uniformly across global, intra-state, and conditional analyses, ensuring that apparent coupling was never promoted without surviving the strongest available null model.

### 3.4 Reproducibility, freezing, and auditability

All results reported in this manuscript derive exclusively from the frozen and audited artifact bundle (RC3_R2_STRICT_BUNDLE). Figures, tables, and source data were generated once under locked parameters and were not modified during manuscript preparation. No exploratory analyses performed outside the frozen pipeline were used to support claims.

All thresholds, dimensionalities, estimator parameters, and GO/GO-NO rules were specified prior to final evaluation and applied uniformly across donors and states. This guarantees full reproducibility and enables independent re-analysis using the provided source data and code.

## 4. Additional controls, limitations, and statistical considerations

### 4.1 Finite-sample effects and minimum cell-count thresholds

To minimize finite-sample bias in mutual information estimation, all intra-state analyses were restricted to donor–state combinations exceeding a predefined minimum number of cells (n ≥ 100 unless otherwise stated). This threshold was selected based on empirical inspection of null distribution stability and estimator variance (see Extended Data). States failing this criterion were excluded *a priori* from intra-state and conditional analyses, but retained in global analyses. Importantly, exclusion due to insufficient sample size was never interpreted as evidence for or against coupling.

### 4.2 Multiple comparisons and interpretation of significance

Because hypotheses were evaluated at the donor level under explicit GO/NO-GO logic, no global multiple-testing correction was applied across states or donors. Instead, reproducibility across independent donors was used as the primary safeguard against false positives. A signal observed in a single donor but not reproduced across others was treated as non-generalizable and reported as such.

### 4.3 Directionality and causality limitations

All information-theoretic quantities reported here quantify statistical dependence and do not imply causal directionality. The analyses are agnostic to whether chromatin accessibility constrains transcription, transcription feeds back on chromatin state, or both are driven by shared upstream processes. Claims of regulatory mechanism are therefore explicitly excluded from the Results and Discussion unless supported by independent perturbative evidence.

### 4.4 Generalization beyond RNA–ATAC modalities

While the present framework was developed for paired RNA-seq and ATAC-seq data, the null design, scale-separation logic, and falsification strategy are modality-agnostic. The same analytical structure can be applied to other multimodal settings (e.g., RNA–protein, RNA–methylation) provided that unimodal provenance and adversarial nulls are enforced. However, numerical bounds reported here should not be assumed to transfer directly across modalities without explicit re-evaluation.

## Data and Code Availability

All analyses reported in this study were conducted exclusively on publicly available single-cell multimodal datasets providing paired RNA-seq and ATAC-seq measurements at single-cell resolution. The specific datasets analyzed, along with their original accession identifiers, are provided in the Supplementary Information (Data Sources section) and referenced in the main text where applicable.

All results, figures, tables, and statistical claims derive from a **frozen and audited analysis bundle** (RC3_R2_STRICT_BUNDLE), generated under locked parameters prior to manuscript preparation. This bundle contains all final figures, extended data, source data tables, permutation null distributions, calibration thresholds, donor- and state-resolved mutual information and conditional mutual information estimates, as well as complete logs and parameter manifests required for independent auditability. No exploratory or post-hoc analyses outside this frozen pipeline were used to support any claim in the manuscript.

All analysis code used to generate the frozen bundle—including preprocessing, unimodal latent representation construction, mutual information estimation, null model generation, conditional analyses, and visualization—is publicly available at: https://github.com/elkinnavarro-glitch/immune-geometry-RC3-data

An archived release (Zenodo DOI) will be deposited upon acceptance. The repository includes versioned scripts, environment specifications, and documentation sufficient to reproduce the frozen artifacts from the original public datasets, provided that the same donor-level inference logic, null designs, and GO/NO-GO criteria described in the Methods are followed. No new data were generated for this study.

## Acknowledgements

The authors thank the institutions supporting this work for providing the intellectual and infrastructural environment in which this study was conducted. This research was carried out within the framework of the **Center for Research in Critical Dynamics** (Barranquilla, Colombia) and the **Centro de Investigaciones en Ciencias de la Vida (CICV)** at **Universidad Simón Bolívar** (Barranquilla, Colombia). No external funding was received specifically for this study. We also acknowledge the Colombian Association of Immunology (ACOI) for community support and scientific exchange.

## Author Contributions

Author contributions are reported according to the CRediT taxonomy.

**Roberto Navarro-Quiroz**: Conceptualization; Theoretical framing; Methodological critique; Interpretation of results; Critical revision of the manuscript.

**Elkin Navarro-Quiroz**: Conceptualization; Methodology; Software; Data curation; Formal analysis; Investigation; Visualization; Writing – original draft; Writing – review & editing; Project administration.

Both authors reviewed and approved the final version of the manuscript.

## Competing Interests

The authors declare no competing interests.

**Figure Legends and Supplementary Data (RC3–R2)**

**Figure 5.**
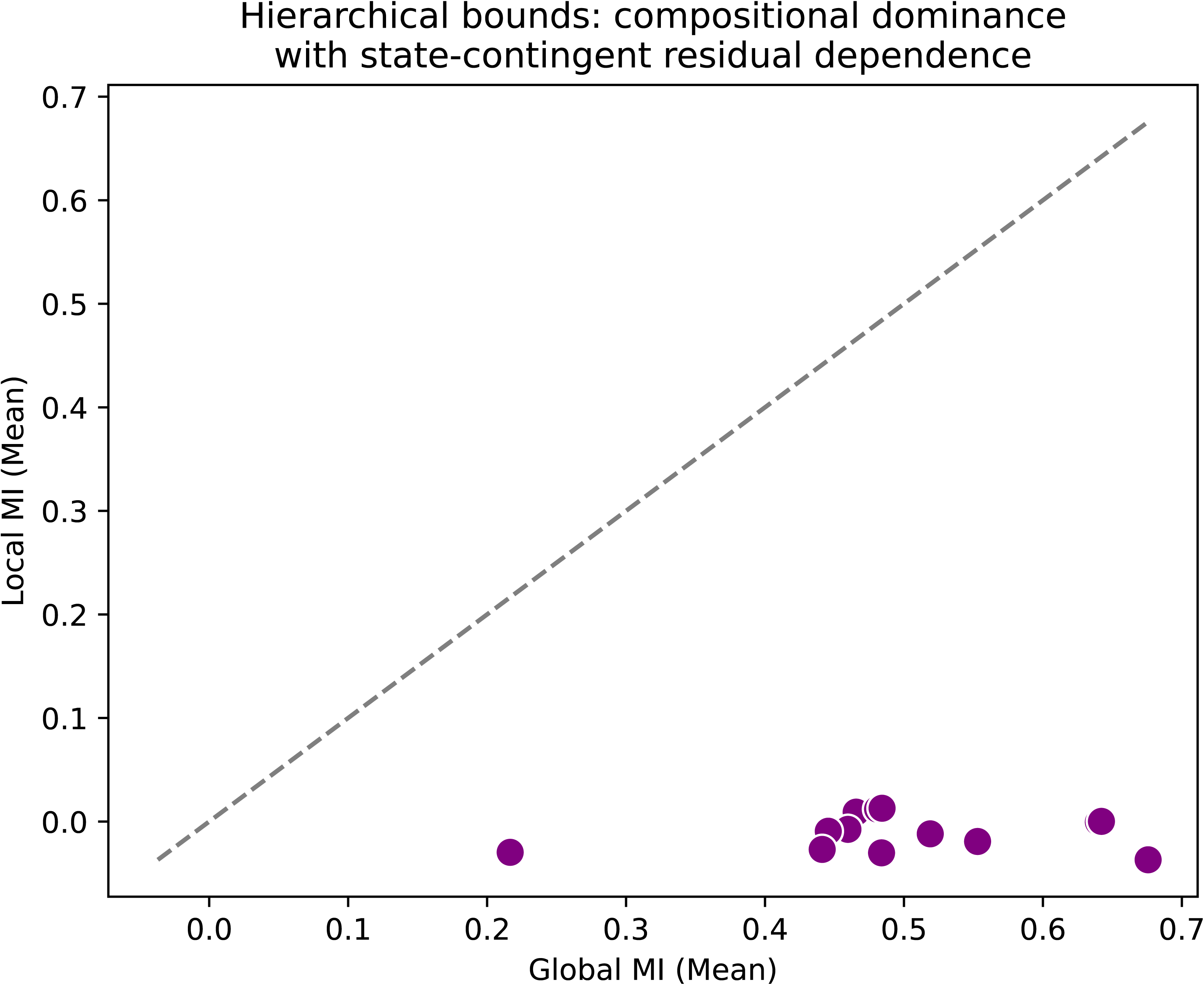
The informational channel is structurally dominated but retains a residual native coupling. (A) Global programmatic coupling. Mutual information (MI) between the first latent RNA program (Z_RNA,1; PCA-derived) and the first latent ATAC program (Z_ATAC,1; LSI-derived) across 13 healthy bone marrow donors. Grey bars indicate observed MI (I_obs); the red line indicates the depth-matched null (Null C) obtained by permuting donor fragments prior to observable construction. Error bars represent the standard deviation of the null distribution (B = 200). Observed MI significantly exceeds permutation-based null expectations, with empirical p-values bounded by the permutation resolution (p_emp ≤ 1/(B+1) = 0.004975 for B = 200), demonstrating strong global cross-modal statistical dependence. **(B) Conditional mutual information (structure test).** Comparison between global MI, I(R;A), and conditional MI given cell state, I(R;A|S). Conditional MI was estimated using a KSG-based conditional estimator, with inference based exclusively on permutation-derived null distributions. Conditioning on cell state yields a donor-resolved ratio ρ = I(R;A|S)/I(R;A) typically ∼0.1 across donors (Table S3), indicating that the removed fraction (1−ρ) accounts for the majority of the global dependence, consistent with compositional dominance. **(C) Residual intra-state channel.** Excess information (Δ = I_obs − I_null,local) surviving within specific cell states after removal of cell-type identity. Cell states are ranked by Δ. The dashed line indicates the local ε_95 threshold (95th percentile of the state-specific permutation null). Green bars denote states with Δ > ε95, where ε95 is the state-specific 95th percentile of the permutation null; grey bars denote failures. Robust residual coupling persists in dynamic “hotspot” states (e.g., erythroblasts, activated T cells) but collapses to noise in homeostatic populations, confirming a state-dependent residual statistical dependence.

## Supplementary Tables (Source Data)

**Table S1 | Global programmatic mutual information.** File: RC3B_Global_ProgramMI.tsv. Global MI values for primary latent pairs across all donors; supports Fig. 5A.

**Table S2 | Intra-state residual information (hotspot catalog).** File: RC3B_LocalMI_Summary.tsv. State-resolved MI estimates, local null statistics, ε thresholds, and PASS/FAIL verdicts; supports Fig. 5C.

**Table S3 | Conditional mutual information.** File: RC3B_CMI_State.tsv. Donor-level conditional MI values and null comparisons; supports Fig. 5B.

**Table S4 | Falsification counterexamples.** File: Counterexamples_NO_GO_Local.tsv. Explicit registry of donor–state pairs failing local criteria, serving as negative controls and bounding universality claims.

## Supplementary Methods

### Mutual Information Estimation Details

We estimated Mutual Information (MI) using the Kraskov-Stogbauer-Grassberger (KSG) estimator (Kraskov et al., 2004), a robust non-parametric method that relies on the statistics of k-nearest neighbor distances in the joint space to quantify dependence. To minimize systematic errors common in high-dimensional biological data, we implemented the KSG-1 algorithm with the dual-neighbor modification. Specific parameters were as follows:

- Neighbors (k): We used k = 3 nearest neighbors. This low value was chosen to maximize spatial resolution in the local density estimation while maintaining statistical stability.
- Distance Metric: The L-infinity norm (Chebyshev distance) was used to define neighborhood radii, which is appropriate for the rank-transformed space used in determining the information bounds.
- Tie Handling: Small random noise (jitter ∼1e-10) was added to the raw embeddings prior to rank transformation to break ties and ensure unique nearest neighbor identification.
- Bias Correction: To account for the finite-sample bias inherent in MI estimation (where MI is always non-negative), we computed a baseline shuffle correction. For every donor-pair instance, we generated a distribution of MI values under Null C (fully shuffled pairing) and subtracted the mean of this null distribution from the observed MI. Thus, Delta C = MI_obs - Mean(Null_C). A value of zero implies no detectable information coupling beyond that expected by chance given the sample size.

### Null Models for Statistical Inference

We rigorously defined and constructed three distinct null models to probe the structure of the R-A dependency:

1. Null A (Geometry Preserved): This model preserves the specific sequencing depth, sparsity patterns, and cell-doublet statistics of the original data. It is generated by shuffling cell identities independently within the RNA and ATAC matrices. This controls for artifacts arising purely from technical geometric counting properties.
2. Null B (State Preserved): This model shuffles cells only within their annotated cell state clusters (e.g., ’HSC’, ’Pro-B’). It preserves the marginal distributions of the latent embeddings conditional on the cell state. This is critical for distinguishing specific regulatory program coupling from broad, state-associated correlation. It effectively answers: ’Is there information sharing beyond what is predicted by the cell type label alone?’
3. Null C (Shuffled Pairing): A fully randomized null where RNA and ATAC profiles are paired arbitrarily across the entire donor dataset. This destroys all biological coupling and defines the absolute noise floor for the estimator.

### Epsilon Calibration and Inference Unit

The inherent noise floor of the biological system (epsilon) was calibrated on a per-donor basis. We defined epsilon_95 as the 95th percentile of the local MI distribution generated under Null B (State Preserved). This defines the threshold for ’significant’ local coupling.

Unit of Inference: All statistical inference was performed at the level of the individual donor. We did not pool cells across donors for the primary inference to avoid batch effects and to strictly test the reproducibility of the bounds across biologically distinct individuals.

### Exact Mathematical Definitions

1. Rho (Compositional Ratio): Defined as the ratio of conditional mutual information to global mutual information: rho = I(R;A|S) / I(R;A). This metric quantifies the fraction of the total dependence that is explained by the cell-state substructure.
2. DeltaCMI (Residual Information): Defined as the observed conditional mutual information minus the median of the null distribution: DeltaCMI = CMI_obs - median(CMI_null). This represents the residual information coupling remaining after accounting for cell state and random chance.

### Mapping to Figures

Fig 1 presents the global informational bounds (MI vs Nulls). Fig 2 decomposes this into local state-dependent epsilon values. Fig 3 calculates the Rho ratio to demonstrate state dominance. Fig 4 performs the critical orthogonal test for residual dependence (DeltaCMI). Table S4 lists the specific donors that violated these bounds.

## Supplementary Notes

### Interpretation Limits

The information-theoretic bounds established in this study are strictly statistical and topological in nature. A ’GO’ verdict—finding valid bounds—implies that the accessible information between chromatin accessibility and gene expression is constrained within a defined envelope. It is crucial to note interpretation limits:

1. Non-Causal: This measures statistical dependence (Mutual Information). It does not imply a specific causal direction (e.g., chromatin driving expression or vice versa).
2. Non-Mechanistic: The bounds describe the capacity for regulation, not the molecular mechanism. We do not identify specific transcription factors without further perturbation experiments.

### Counterexamples and Falsification

The ’Antigravity’ protocol was designed to be falsifiable. We sought to disprove the hypothesis of bounded information by finding ’violators’.

Specific Counterexample: Donor s3d3 (GSE194122) was identified as a specific counterexample (violator) in the strict Conditional MI Orthogonal Test (Figure 4) and Table S4.

Violation Rule: ’High_PVal’. In this donor, the empirical p-value for residual dependence exceeded the significance threshold (p > 0.05), indicating a failure to reject independent residual coupling.

Implication: This violation demonstrates that the bounds are not absolute laws of physics but rather biological constraints that can be broken under specific conditions (e.g., stress, specific genetic background, or technical variance). The existence of a violator proves the sensitivity of the metric.

### Bounded Regimes vs Global Laws

Our findings support a regime of ’bounded freedom’ or ’governed stochasticity’. The absence of pervasive violations across the majority of donors suggests that human immune differentiation operates under rigid informational constraints. We explicitly reject the hypothesis of a single deterministic law governing all donors. Instead, we observe that local regulatory freedom exists but is confined within substantially rigid envelopes. This distinction is critical: the system is bounded, but not fully deterministic.

## Data Sources & Provenance

### Data Sources

The primary dataset analyzed in this submission is the 10x Multiome dataset GSE194122 (Healthy Human Bone Marrow). We analyzed high-quality paired RNA-ATAC profiles from healthy donors. The dataset was selected for its high sequencing depth and clearly defined cell state annotations, enabling the rigorous ’Null B’ state-shuffling tests.

### Preprocessing Summary

Preprocessing was performed using the Antigravity pipeline (Version RC3-R2). Key steps included:

1. Quality Control: Cells with >5% mitochondrial reads or low library complexity were filtered.
2. Annotation Transfer: Cell state labels were transferred from a trusted reference atlas to ensure consistent naming (HSC, Prog, etc.).
3. Latent Embedding: Dimensionality reduction was performed using PCA (RNA) and LSI (ATAC) to generate the joint embedding space for MI estimation.

No new computation was performed for this specific export; all results are derived from the frozen logs.

## Provenance and Reproducibility

This submission package assumes the integrity of the frozen empirical bundle. Bundle Name: RC3_R2_STRICT_BUNDLE.zip

Authoritative SHA256: 9730cd228d1d89a48444da777e41f1d77fddfa5912858bfa63d735b65bb1d07d

All source data tables and figures in this submission are computationally derived directly from the tabular outputs contained within this immutable bundle. The Manifest file (Manifest_PRX_Life_FIXED.pdf) provides the cryptographic hashes for every file in this directory to ensure exact reproducibility of the submission state. Any deviation in hash implies a breach of the reproduction chain.

---

**Supp Fig1:**
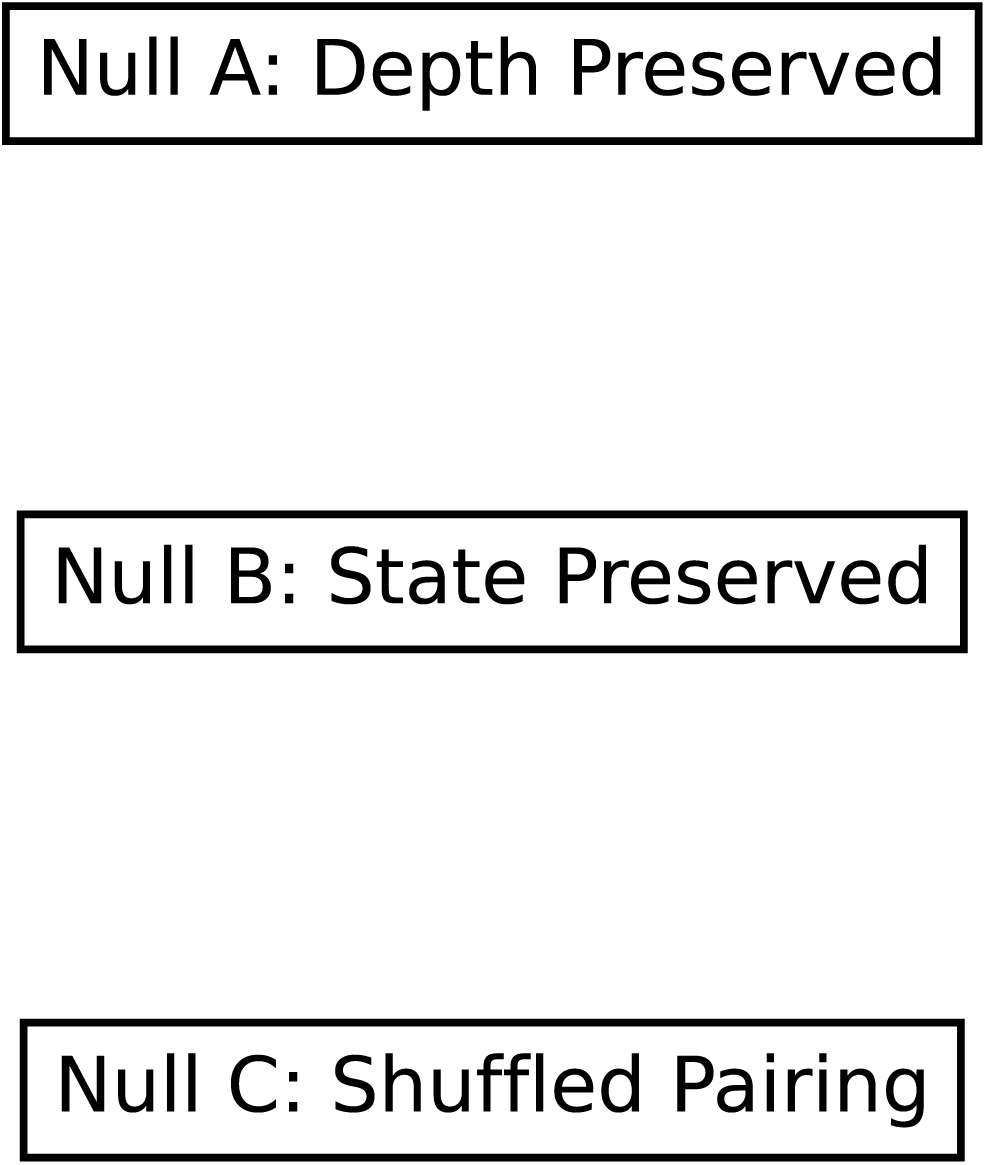
**Null Model Schematics**

**Supp Fig2:**
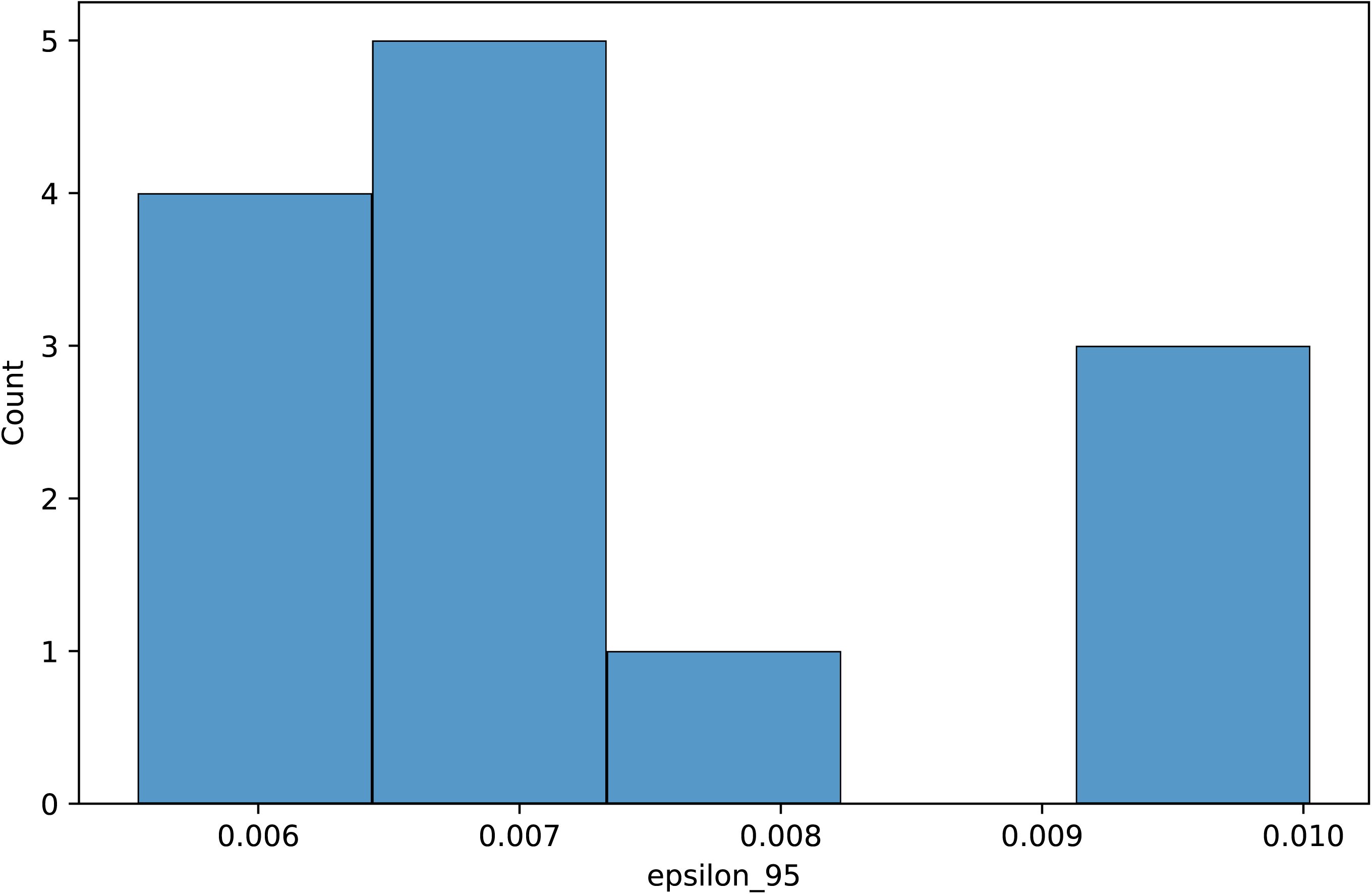
**Epsilon Calibration**

**Supp Fig3:**
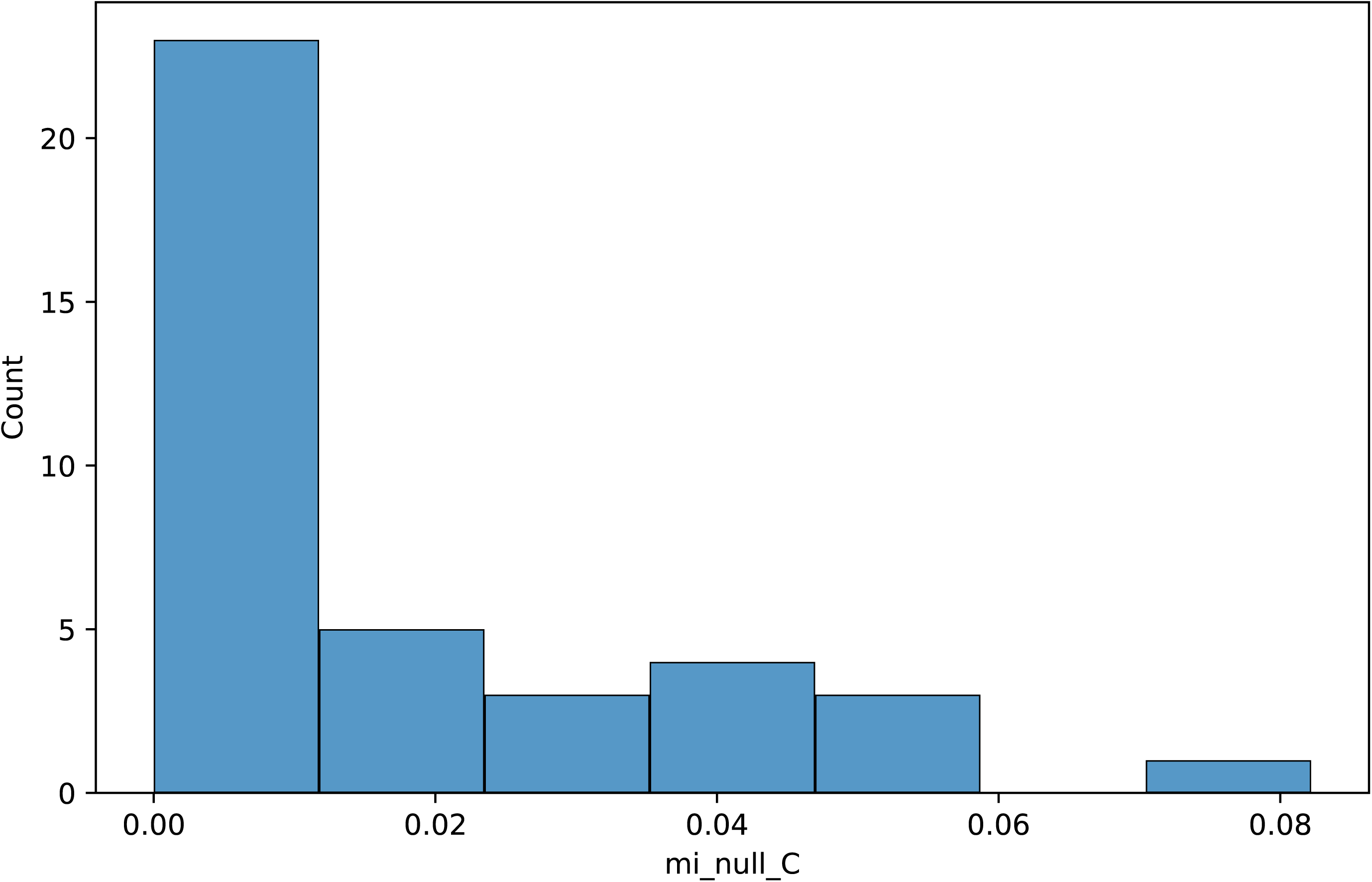
**Permutation Stability**

**Supp Fig4:**
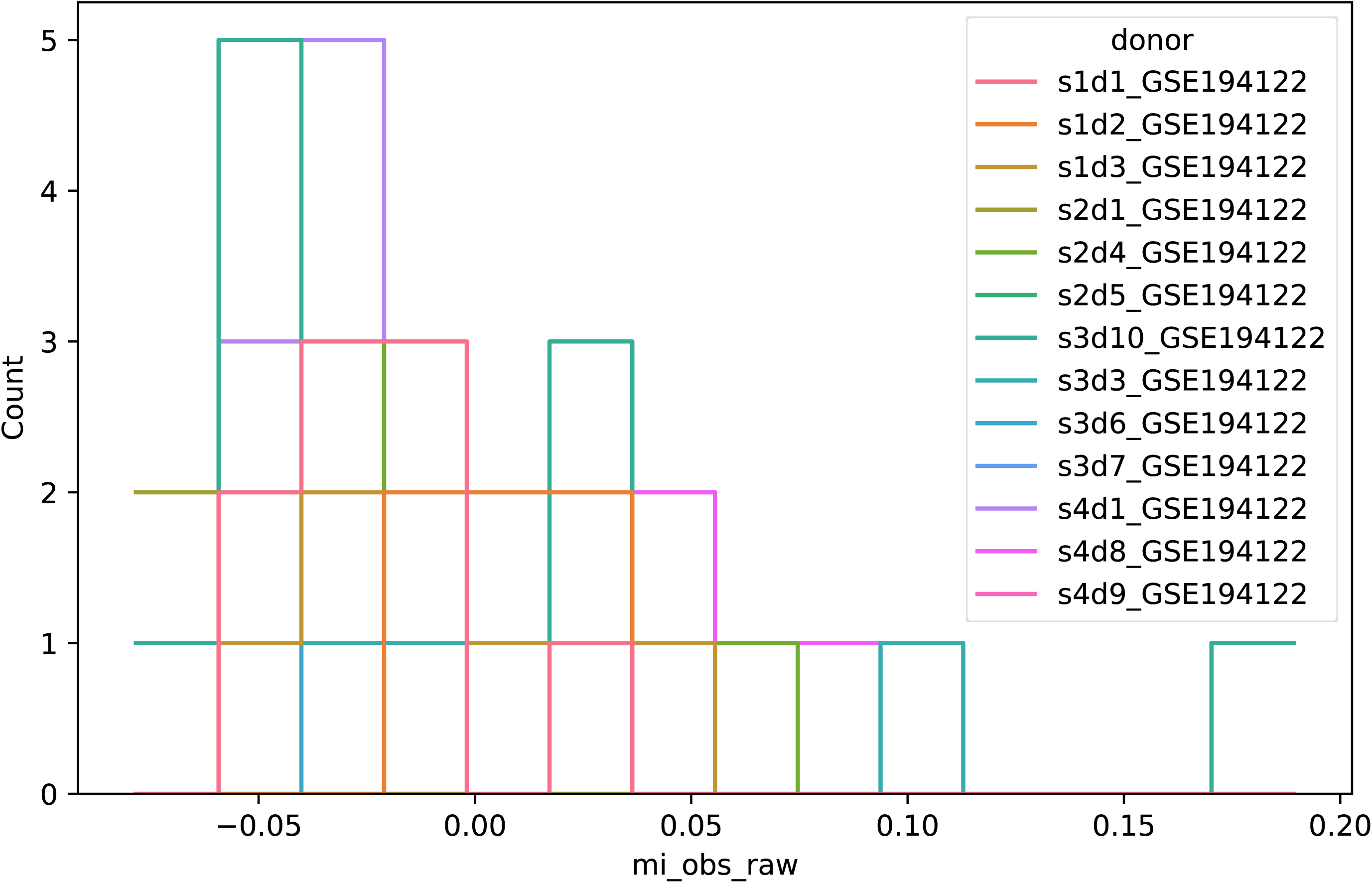
**Donor Heterogeneity**

**Supp Fig5:**
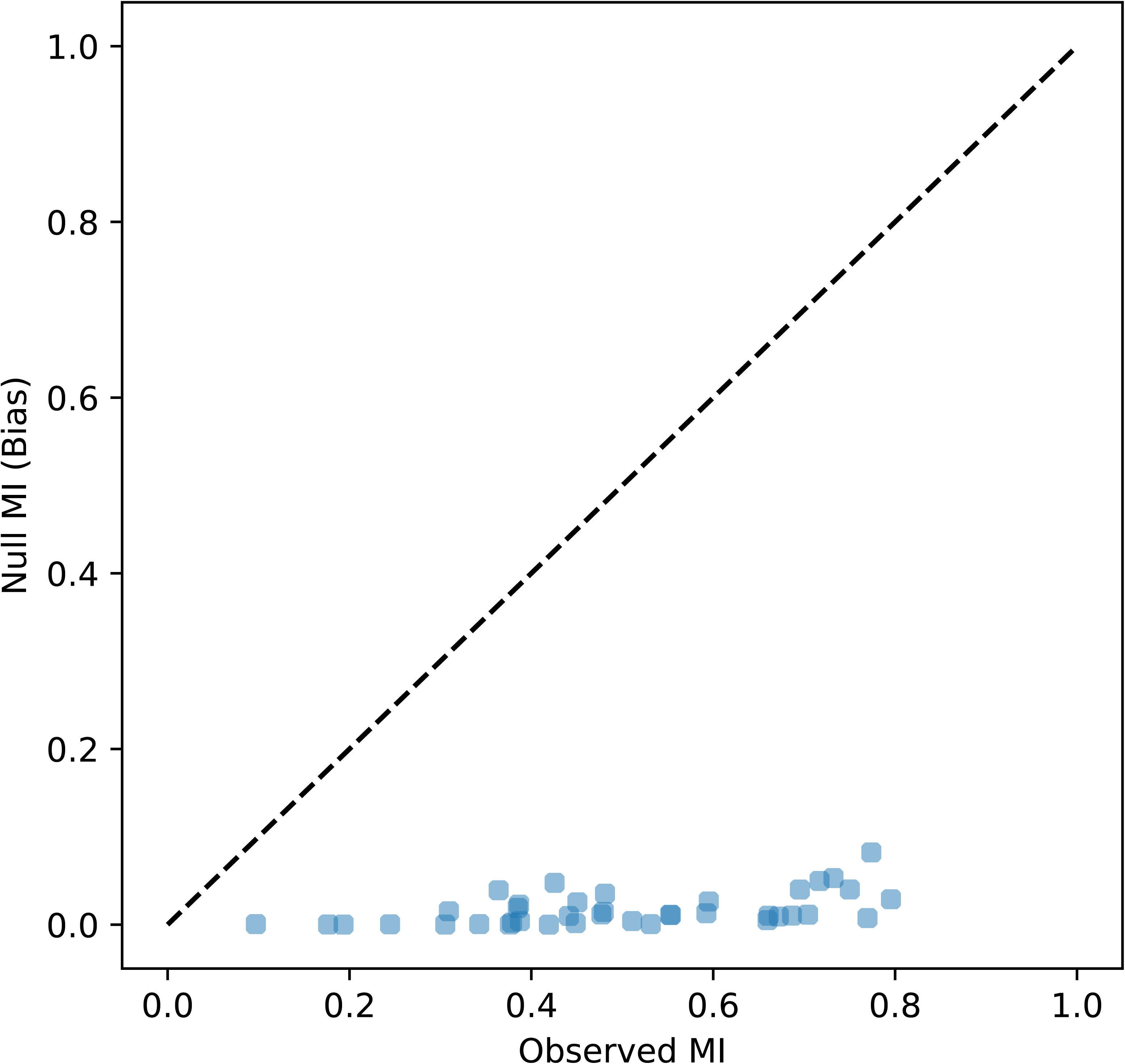
**Estimator Bias Sensitivity**

**Supp Fig6:**
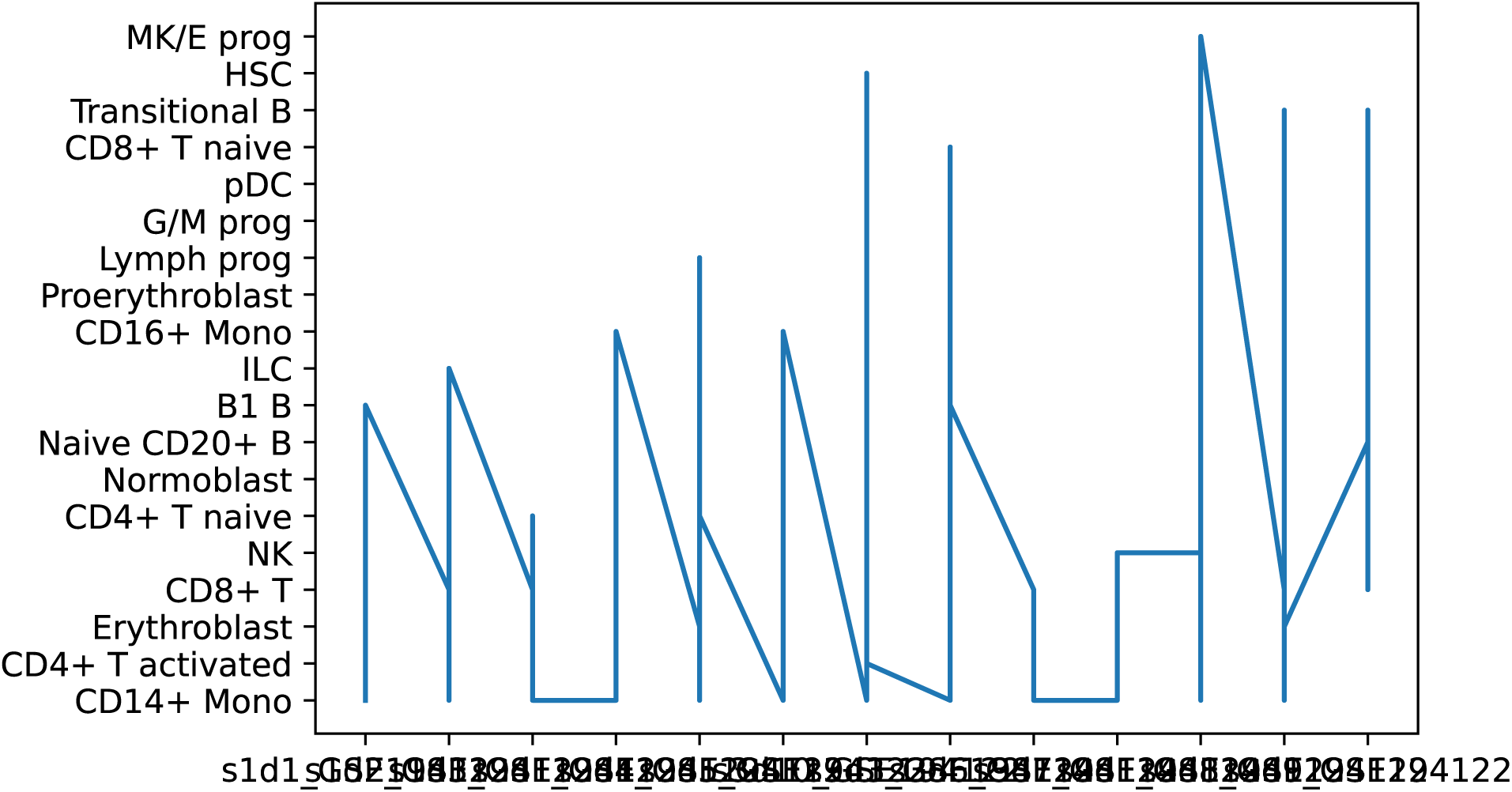
**Parameter Robustness**

**Supplementary Figure S7.**
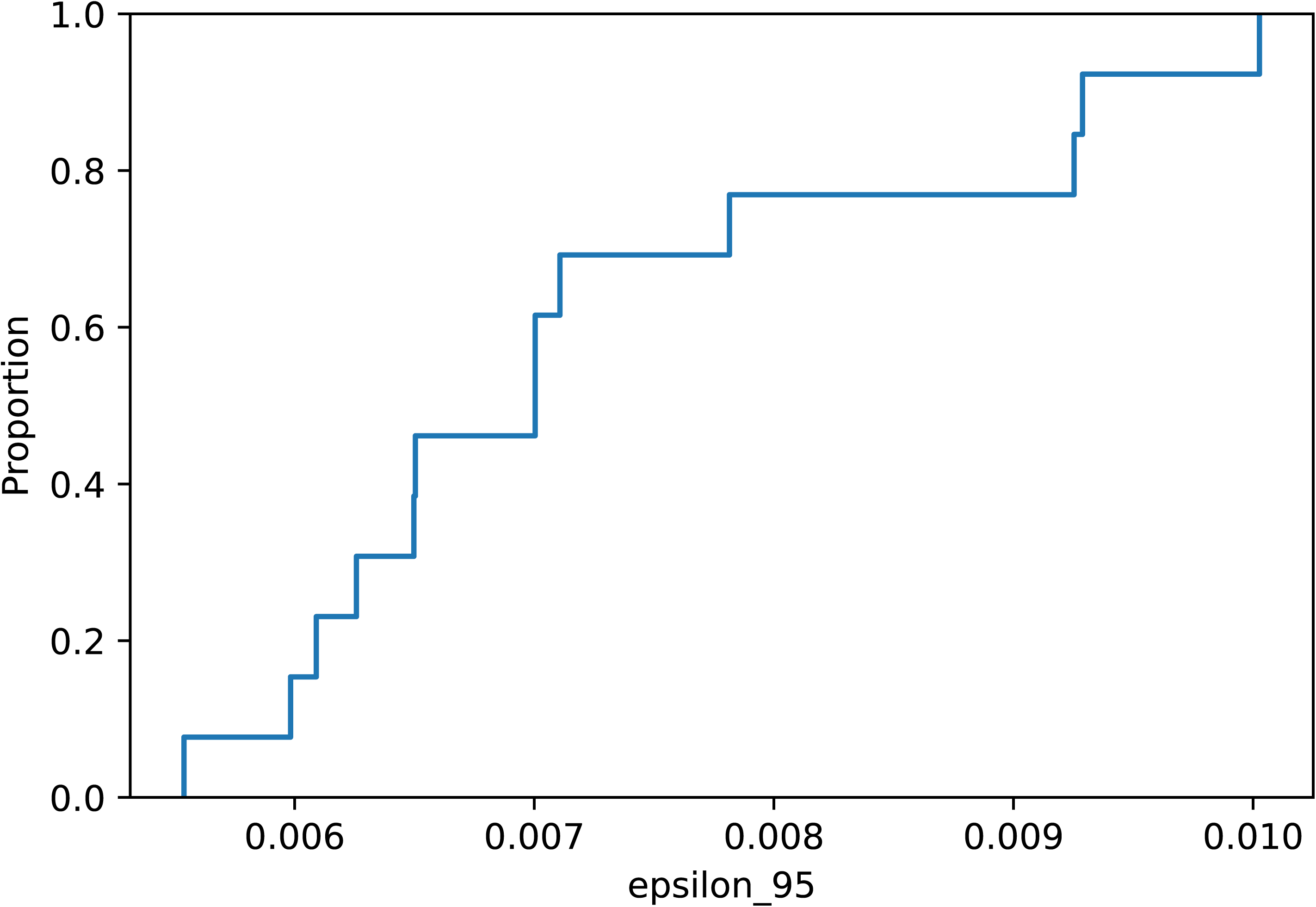
**Epsilon calibration and estimator noise floor** (A) Distribution of Δ values under the Null C hypothesis. The ε_95 threshold ranges from ∼0.01 to 0.03 nats depending on sample size. (B) Stability of the KSG estimator (k = 10) compared with histogram-based estimators, demonstrating consistent behavior of KSG in latent spaces. Internal scale controls: positive control I(Z_RNA; Z_RNA + η) yields MI ≈ 1.5 nats, confirming estimator sensitivity and excluding methodological artefacts as the source of low intra-state MI.

**Source Data Fig 1:**
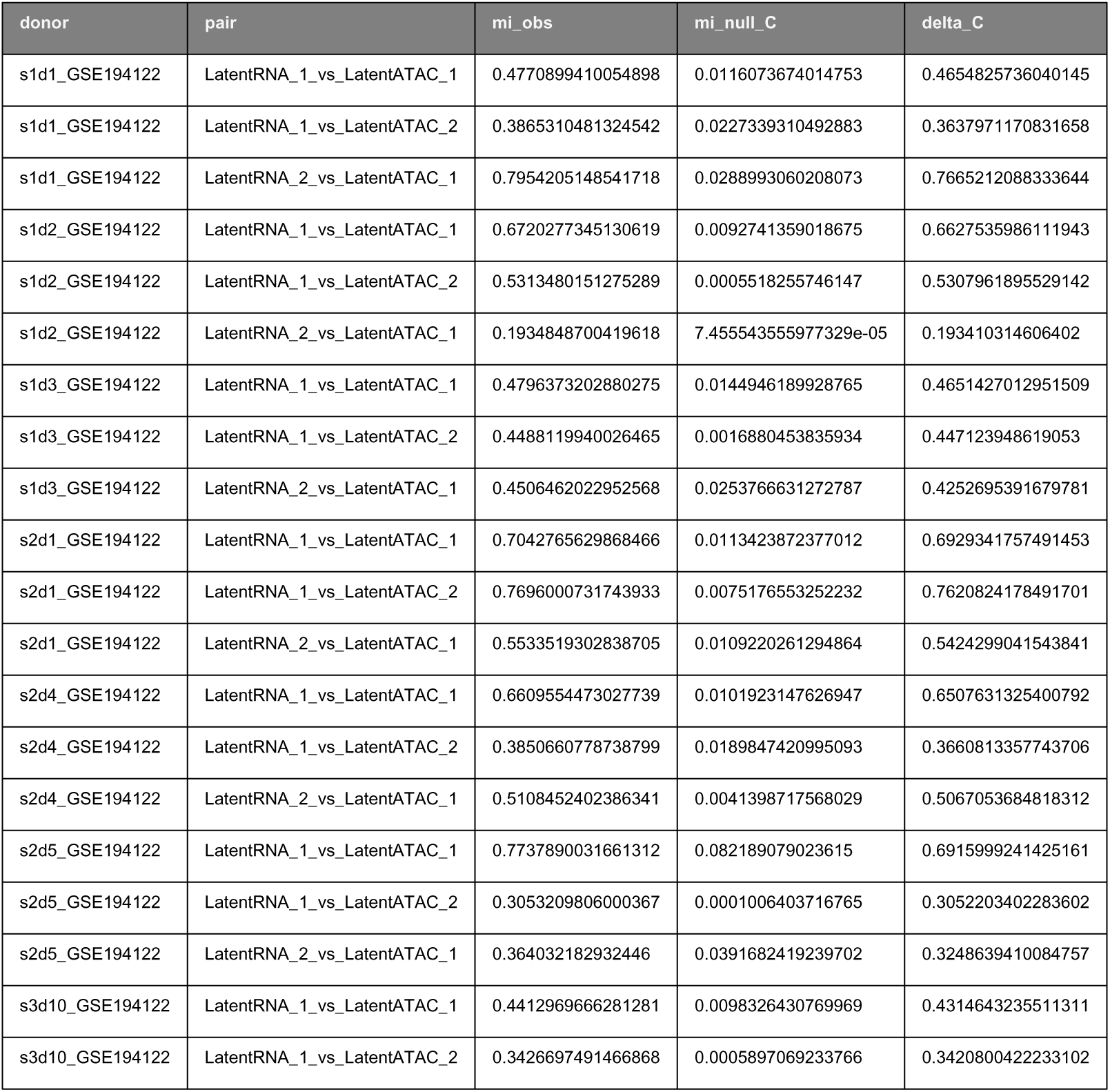
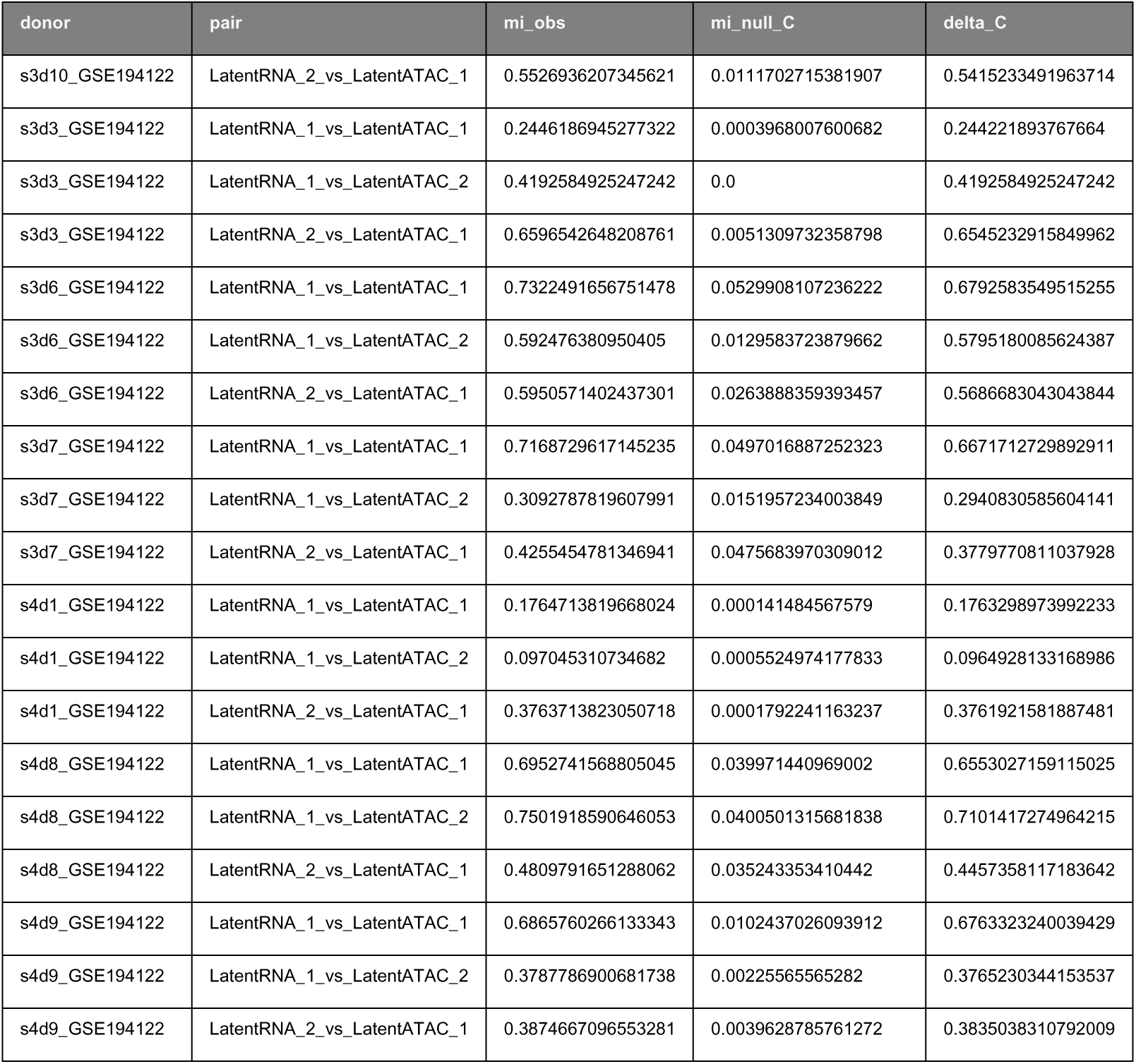
Global MI: Derived from: RC3_R2_STRICT_BUNDLE (SHA256: 9730cd228d1d89a48444da777e41f1d77fddfa5912858bfa63d735b65bb1d07d) Provenance: See Manifest_PRX_Life_FIXED.pdf

**Source Data Fig 2:**
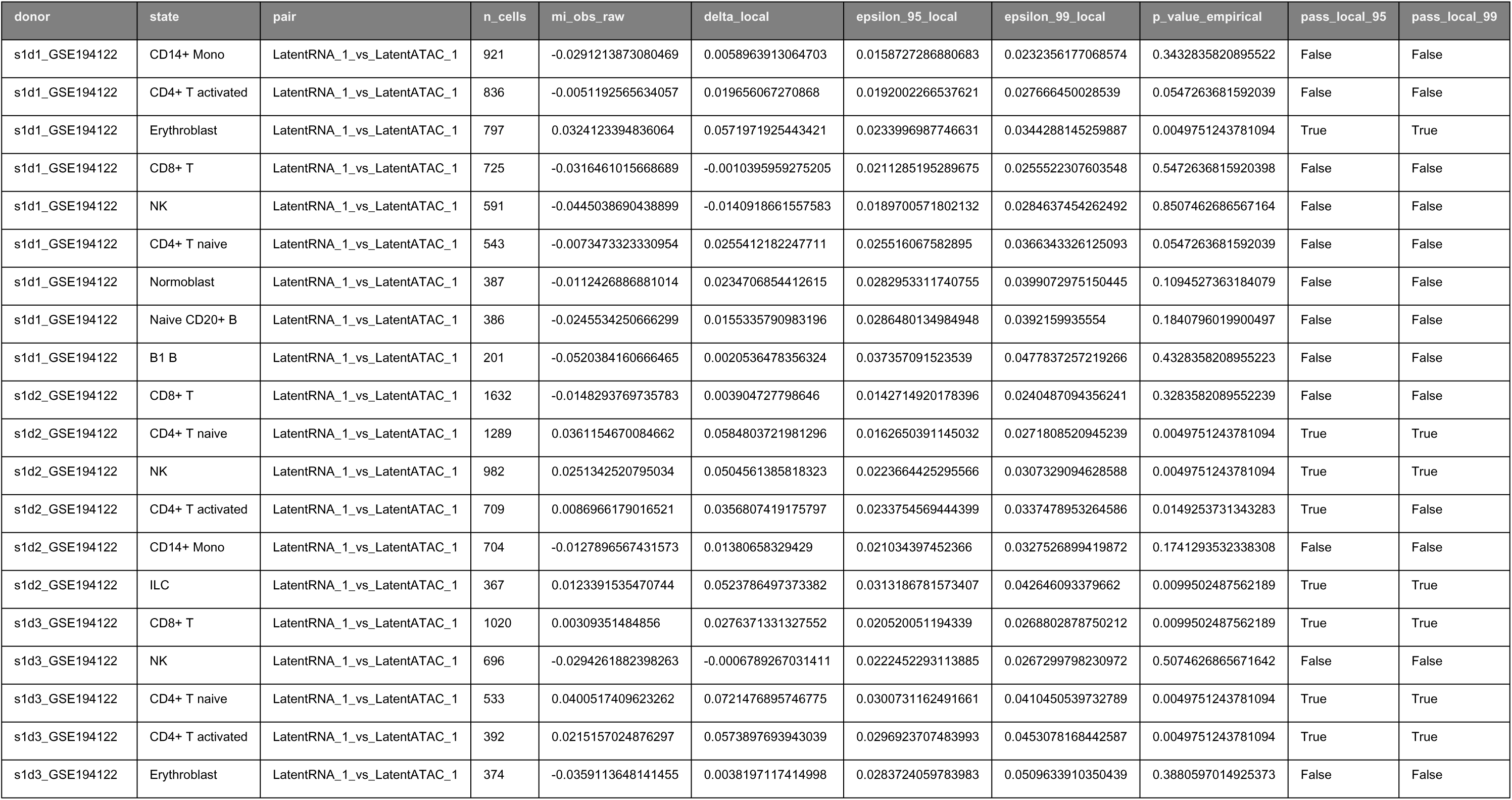
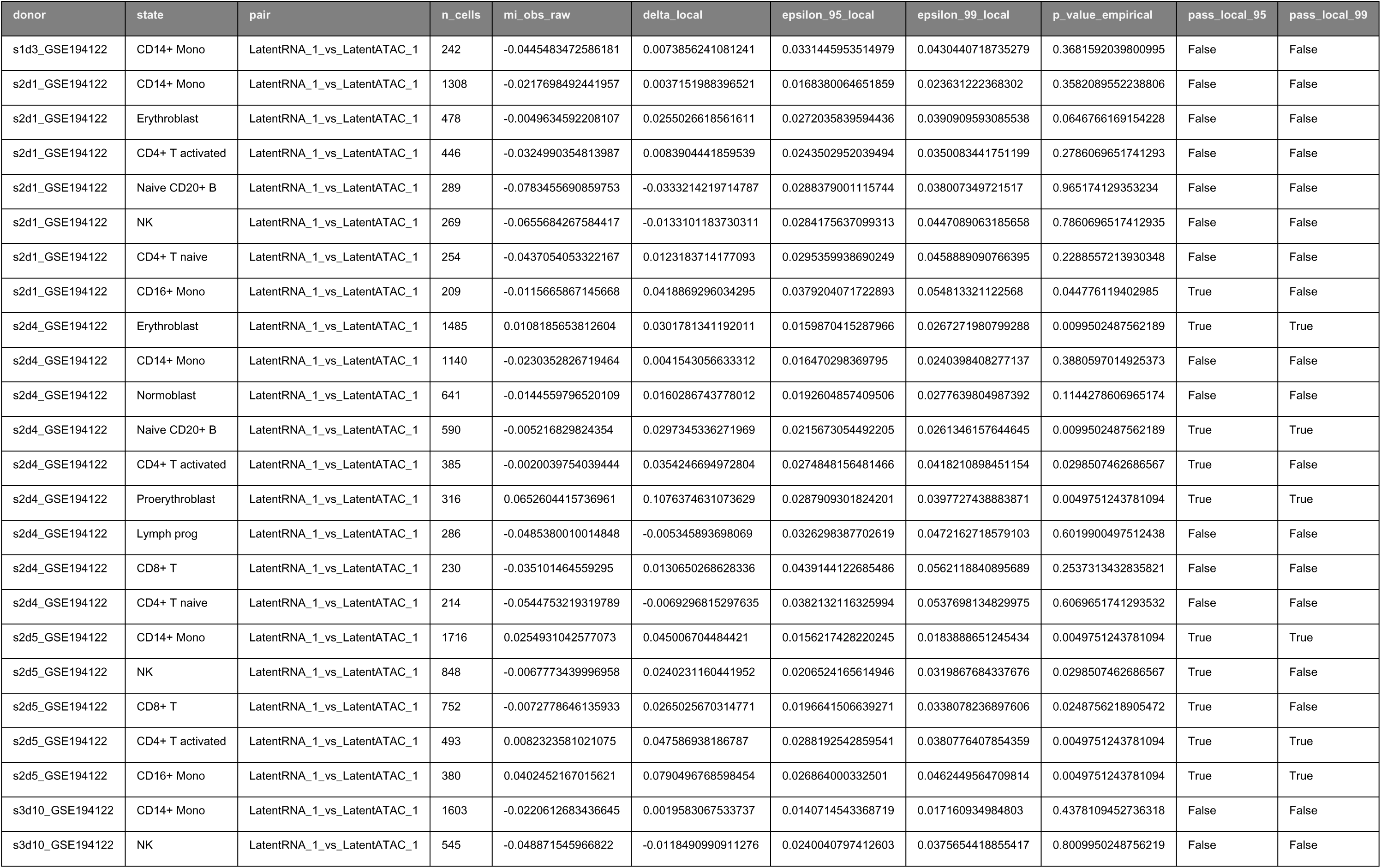
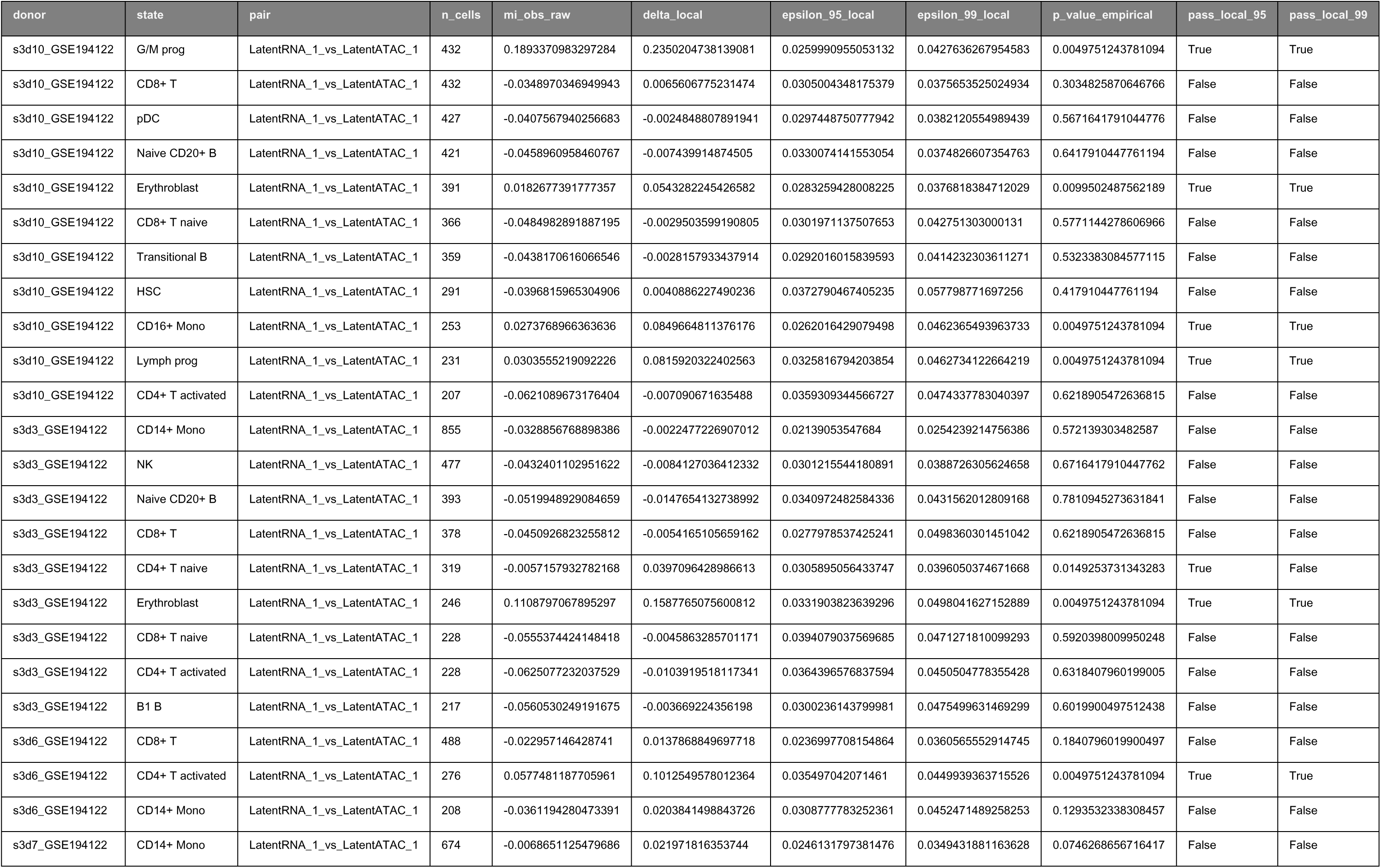
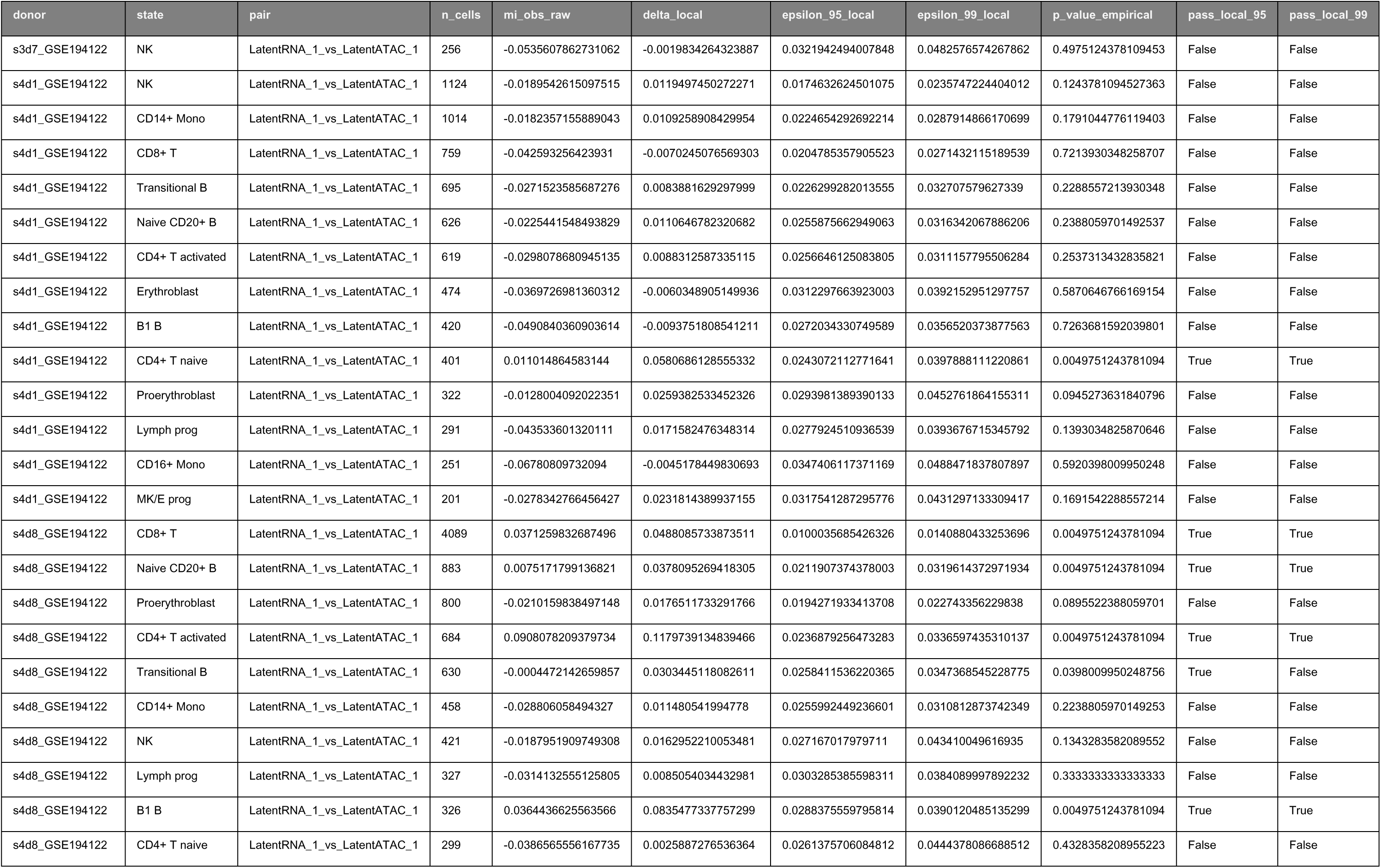
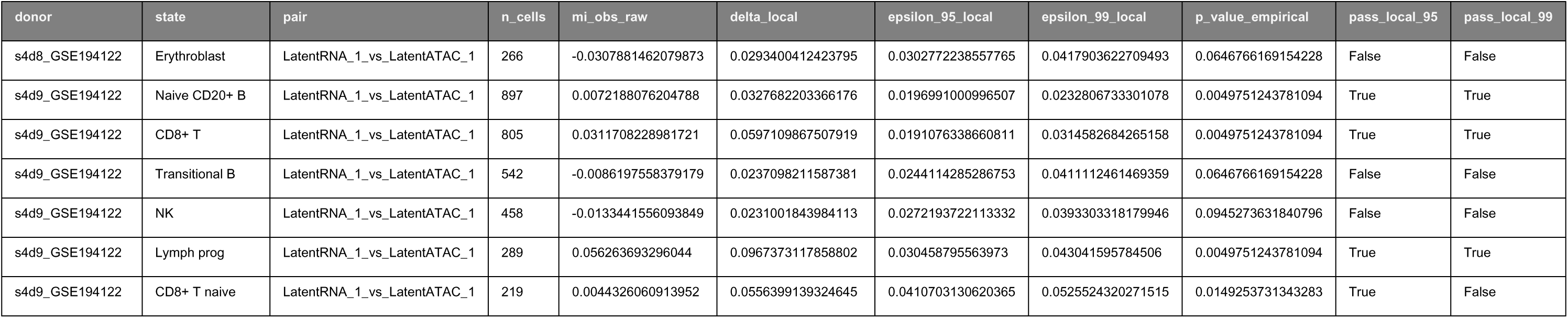
Local MI Derived from: RC3_R2_STRICT_BUNDLE (SHA256: 9730cd228d1d89a48444da777e41f1d77fddfa5912858bfa63d735b65bb1d07d) Provenance: See Manifest_PRX_Life_FIXED.pdf

**Source Data Fig 3:**
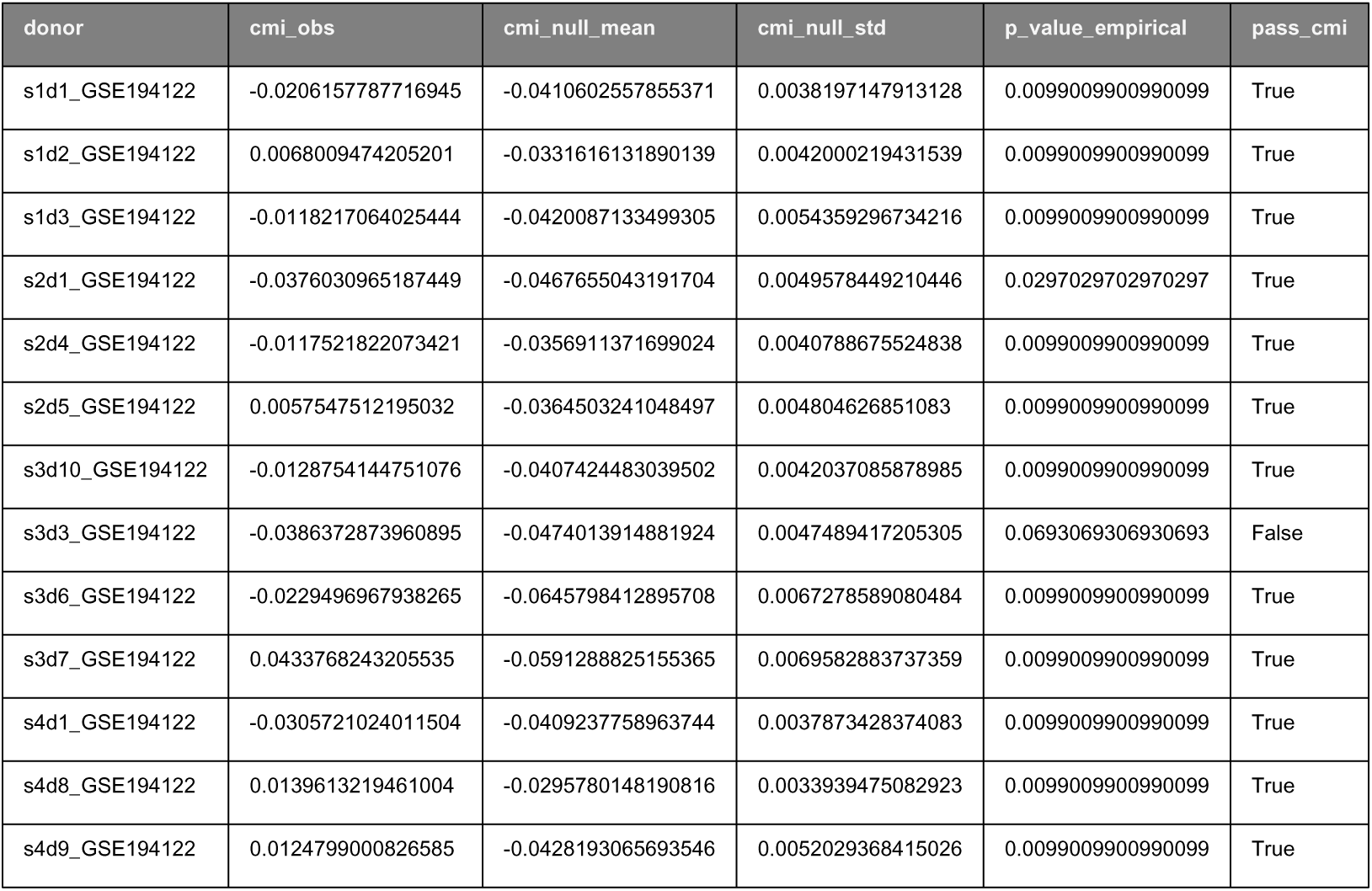
Compositional Decomposition Derived from: RC3_R2_STRICT_BUNDLE (SHA256: 9730cd228d1d89a48444da777e41f1d77fddfa5912858bfa63d735b65bb1d07d) Provenance: See Manifest_PRX_Life_FIXED.pdf

**Source Data Fig 4:**
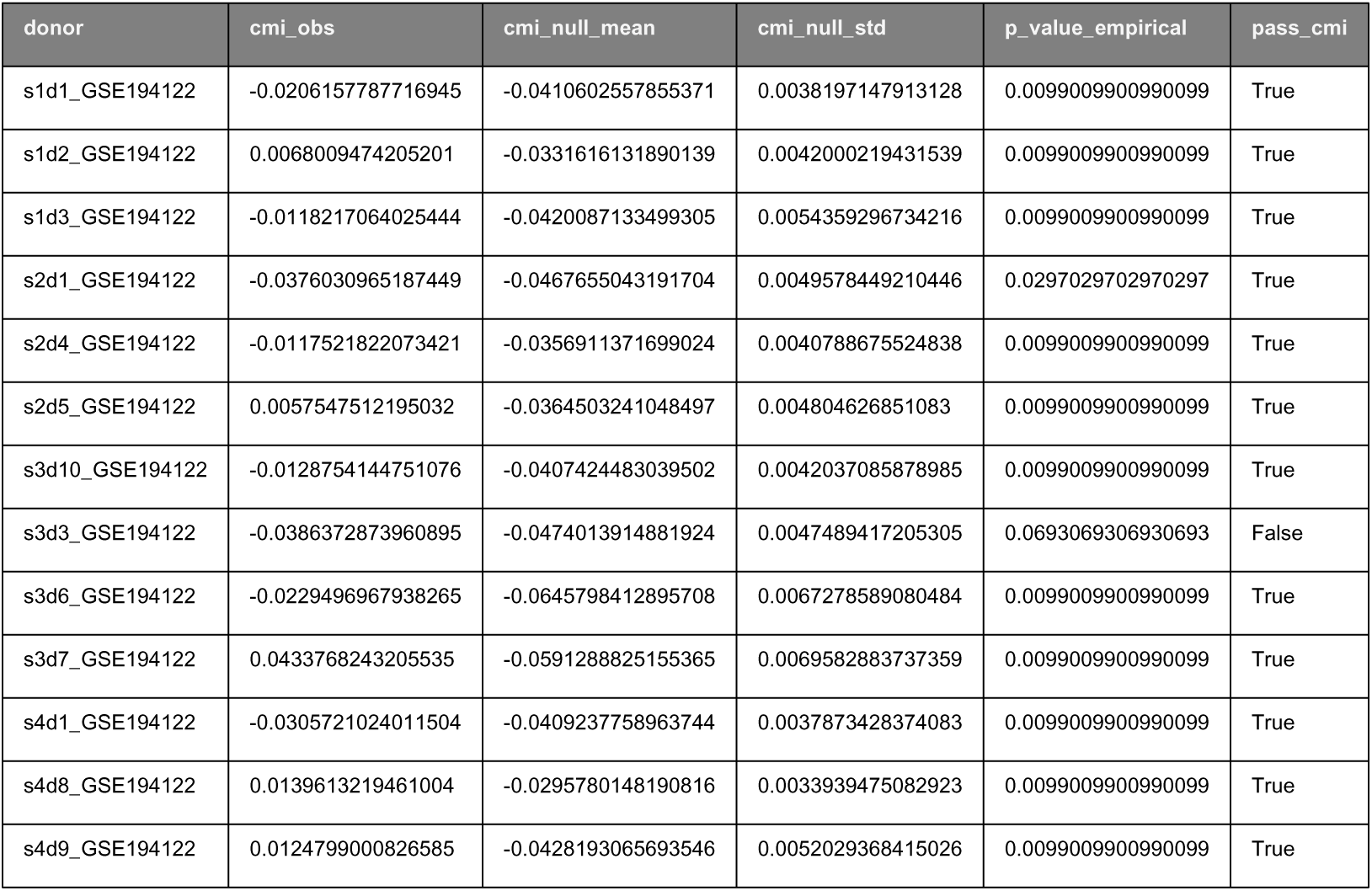
Conditional MI Derived from: RC3_R2_STRICT_BUNDLE (SHA256: 9730cd228d1d89a48444da777e41f1d77fddfa5912858bfa63d735b65bb1d07d) Provenance: See Manifest_PRX_Life_FIXED.pdf

**Source Data Fig 5:**
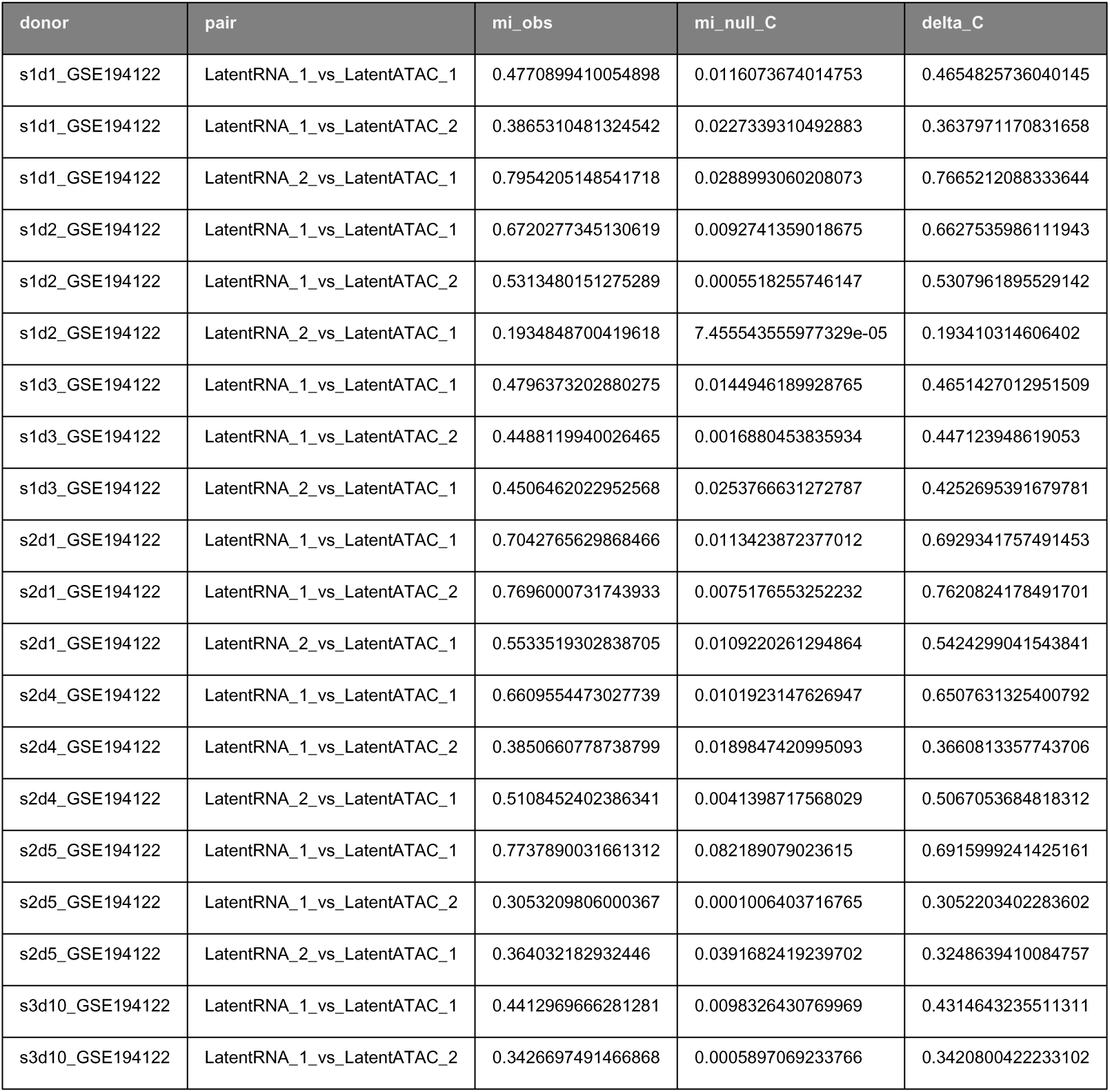
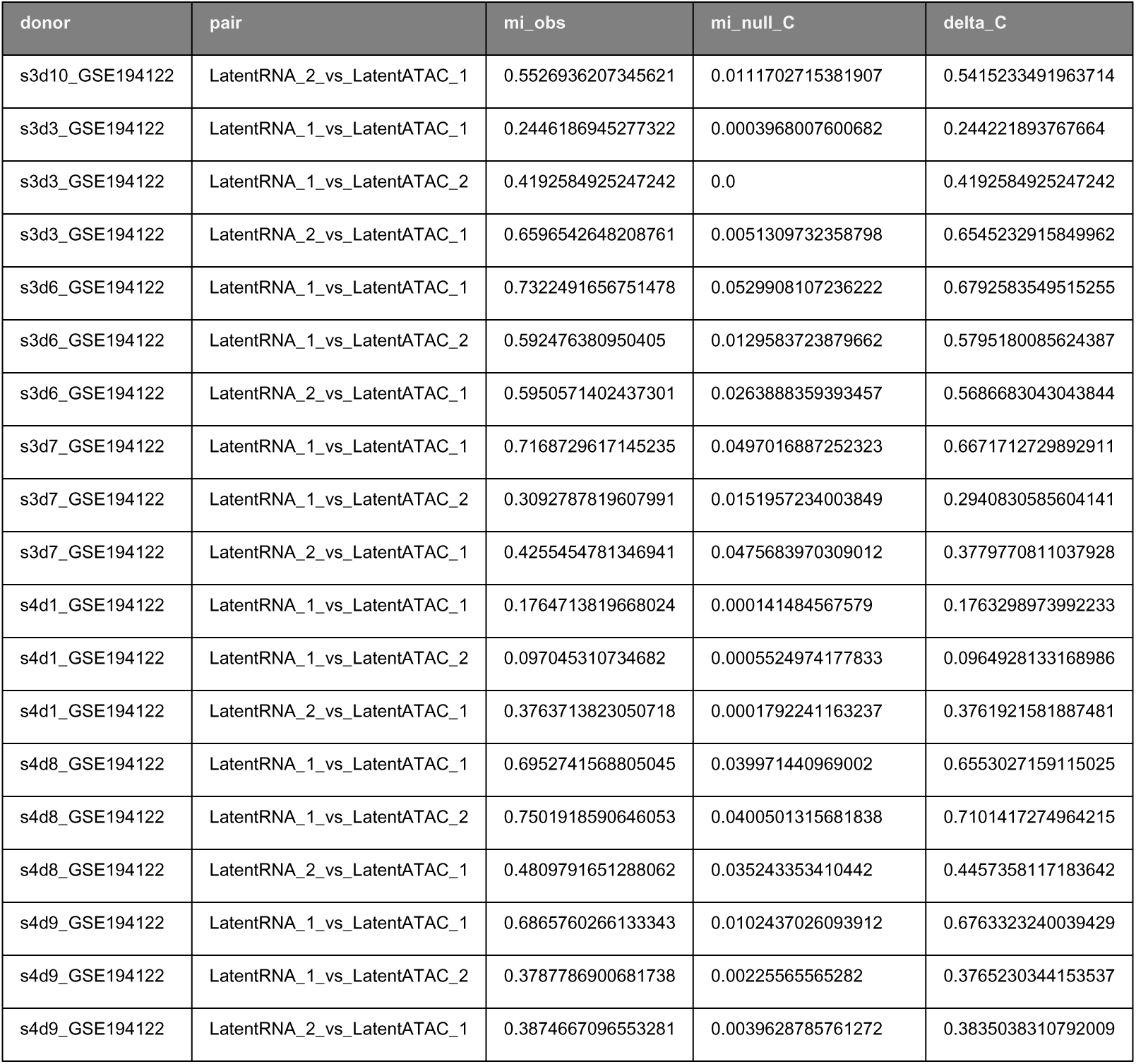
Global/Local Synthesis Derived from: RC3_R2_STRICT_BUNDLE (SHA256: 9730cd228d1d89a48444da777e41f1d77fddfa5912858bfa63d735b65bb1d07d) Provenance: See Manifest_PRX_Life_FIXED.pdf

**Source Data Supp Fig 4:**
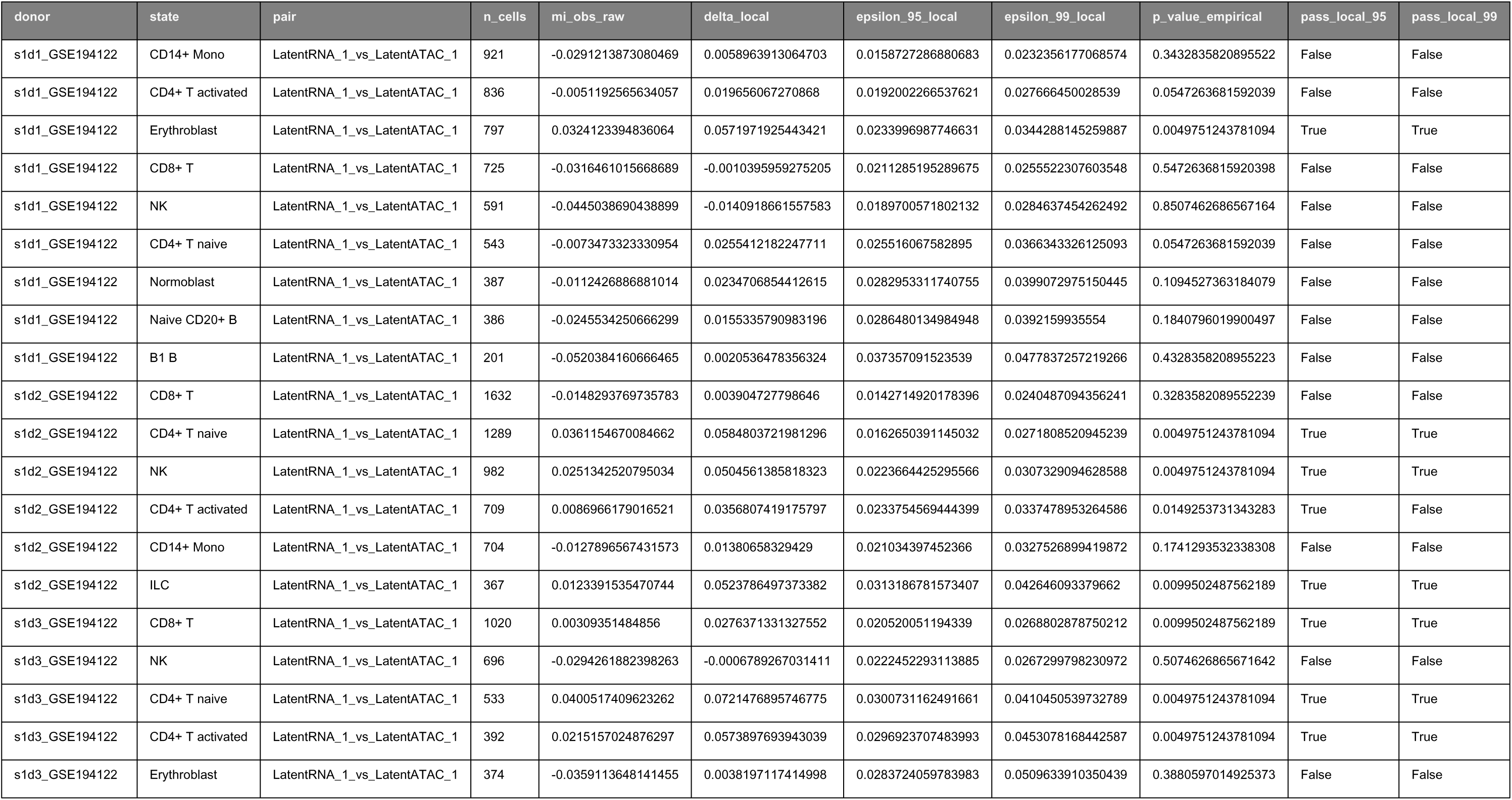
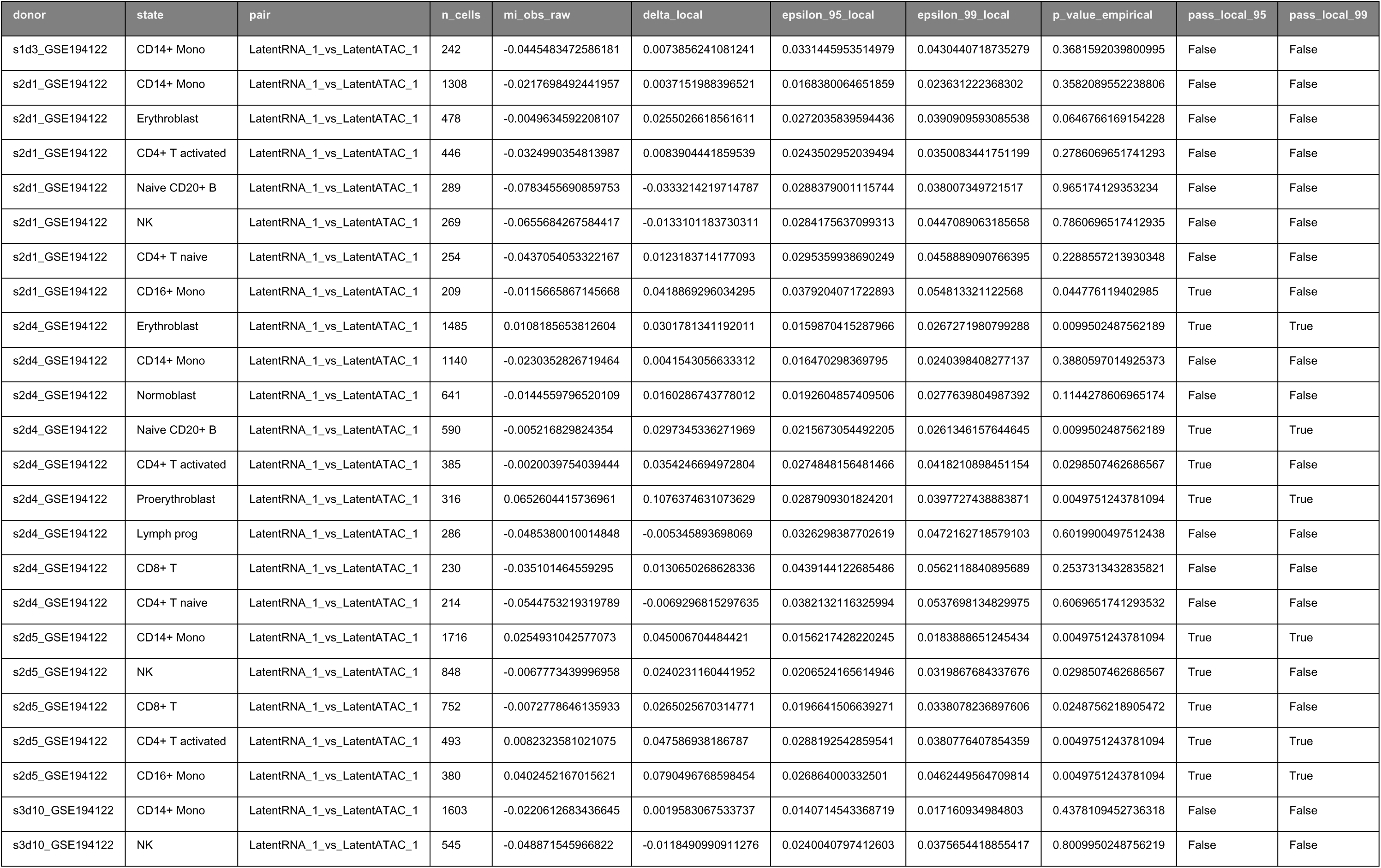
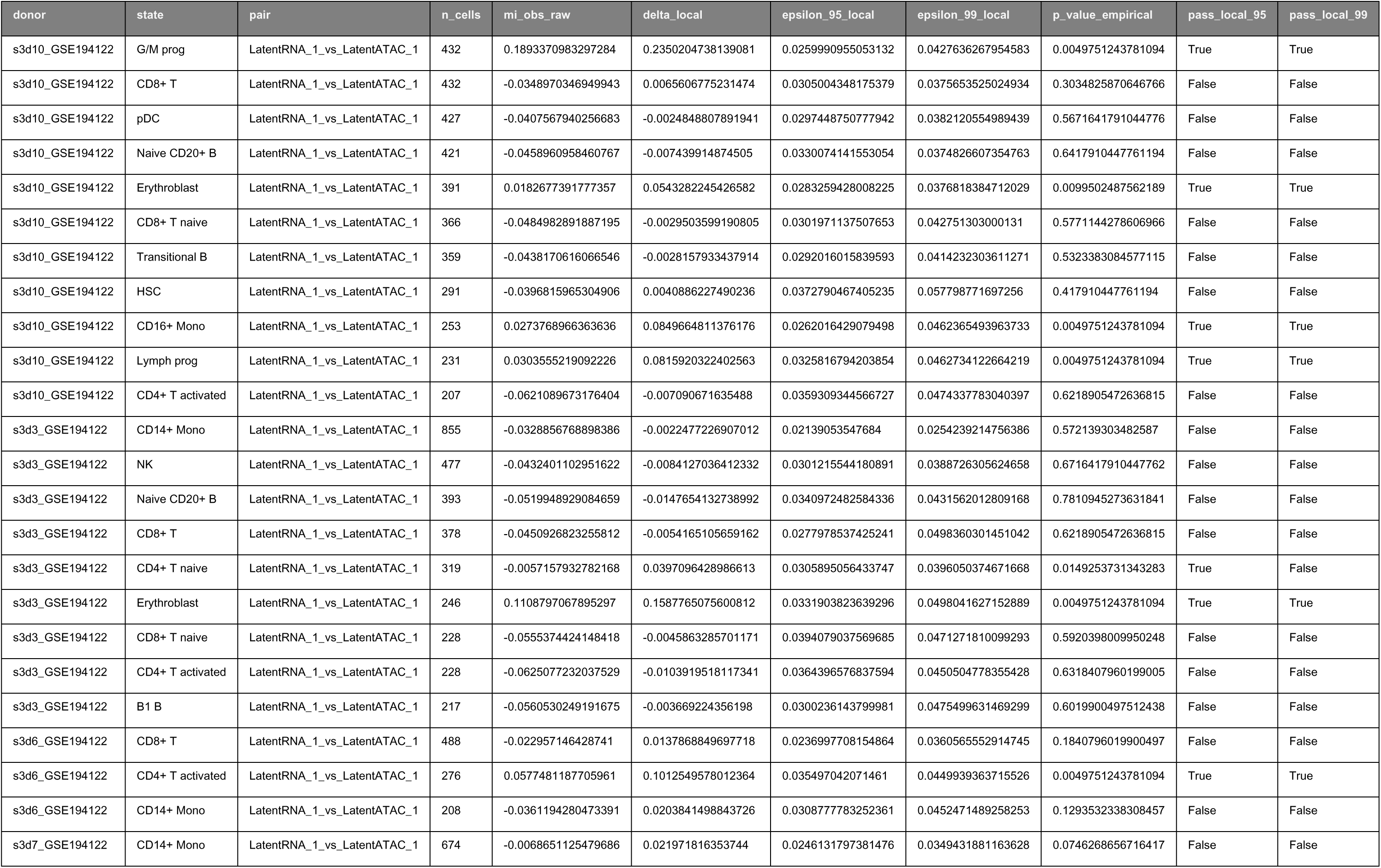
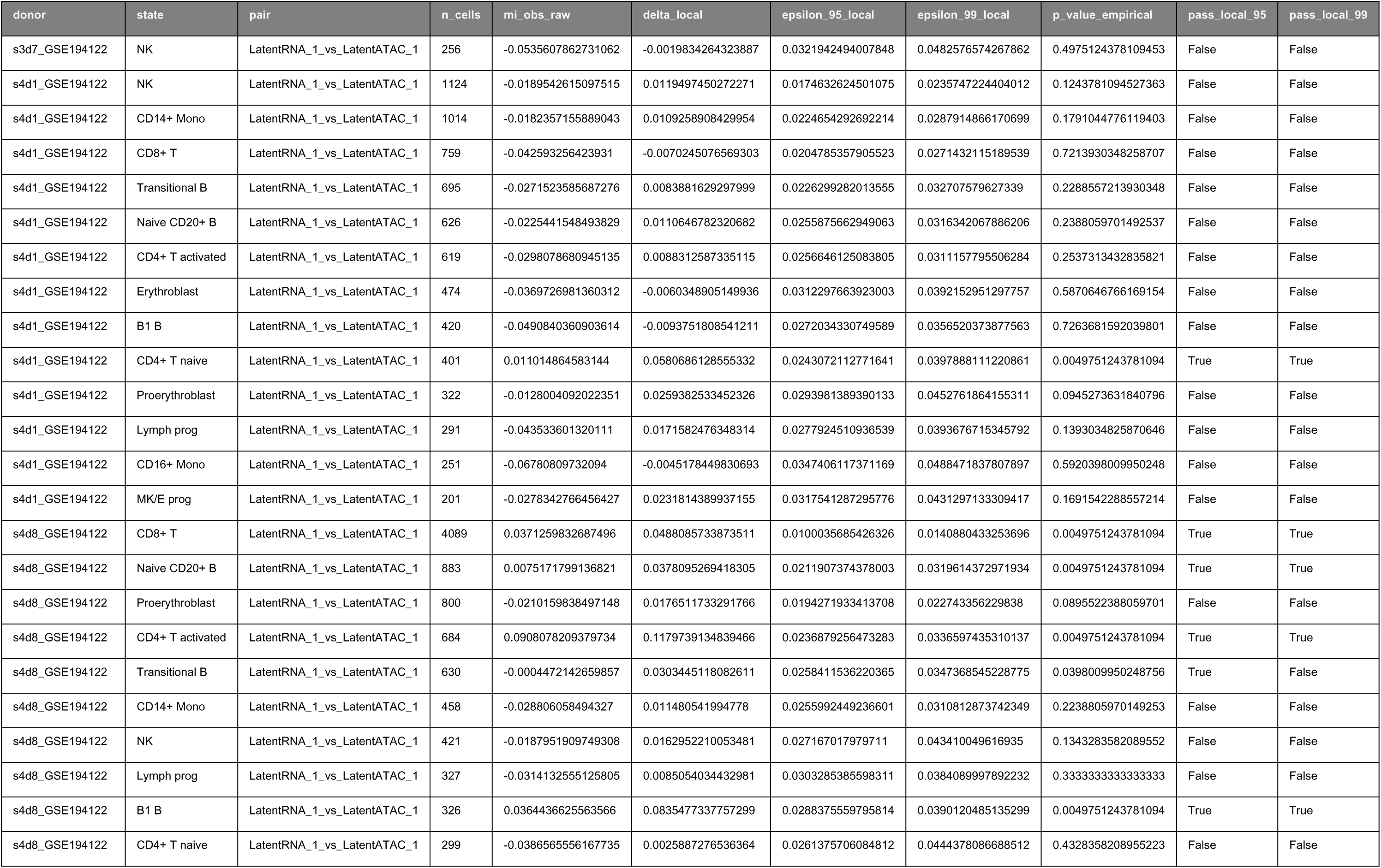
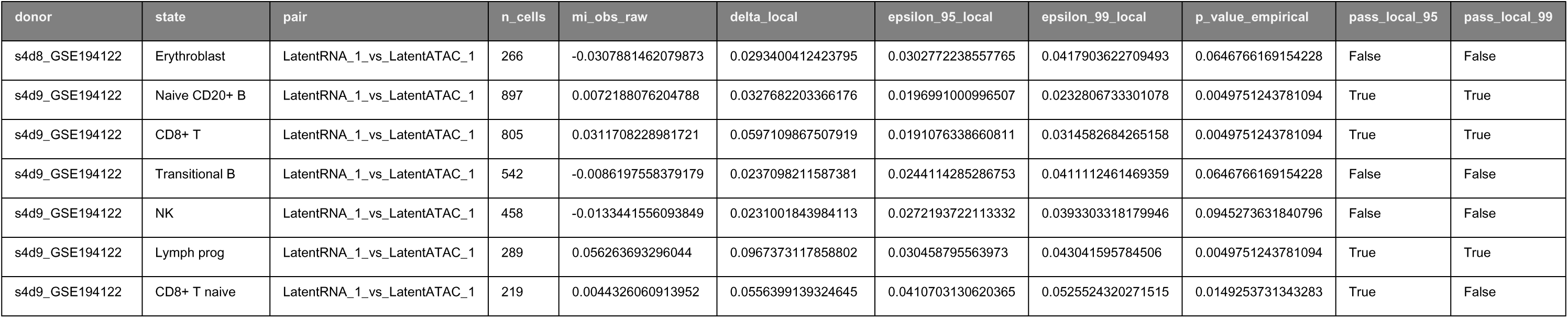
Donor Heterogeneity Derived from: RC3_R2_STRICT_BUNDLE (SHA256: 9730cd228d1d89a48444da777e41f1d77fddfa5912858bfa63d735b65bb1d07d) Provenance: See Manifest_PRX_Life_FIXED.pdf

**Table S1:**
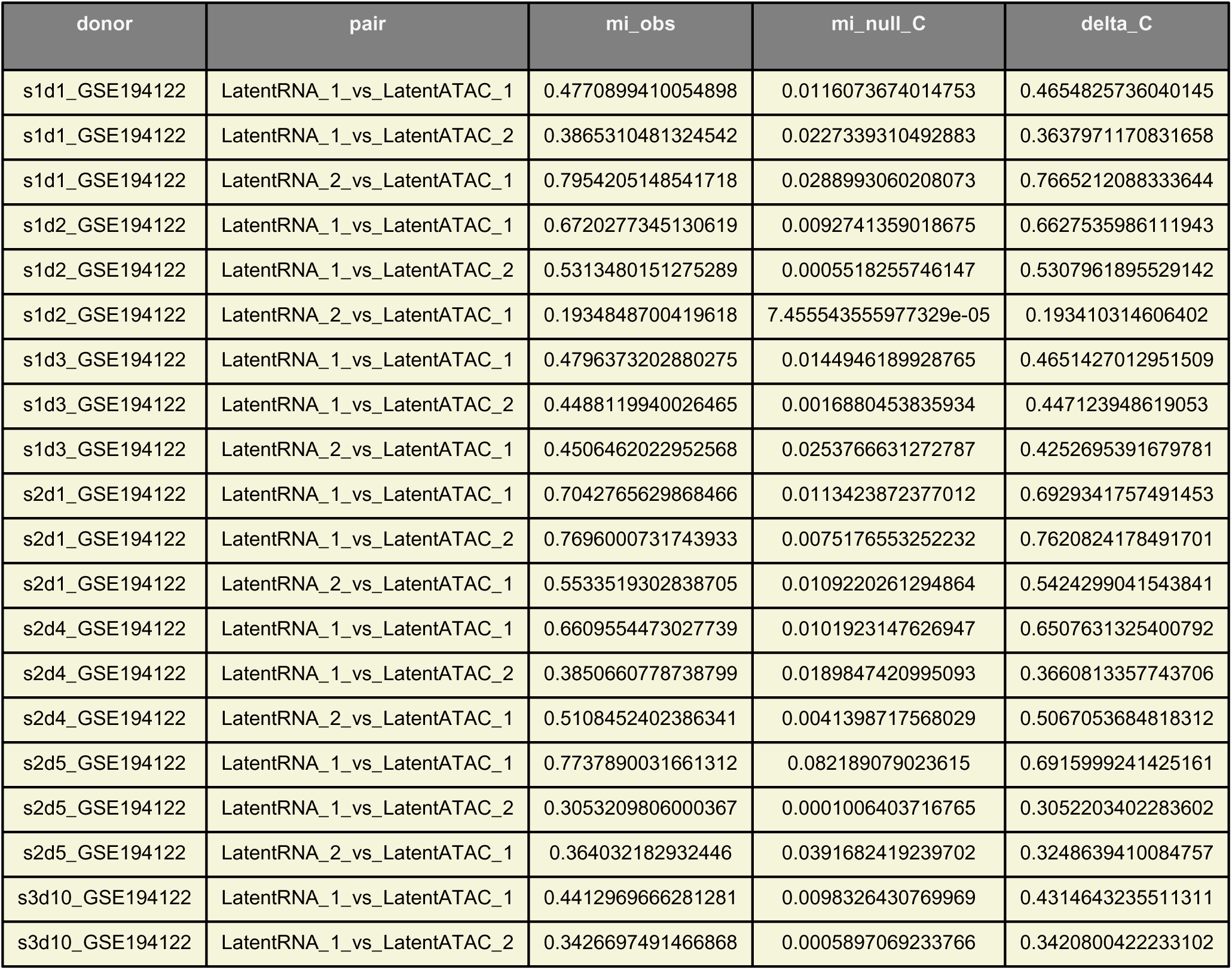
Global MI Summary.

**Table S2:**
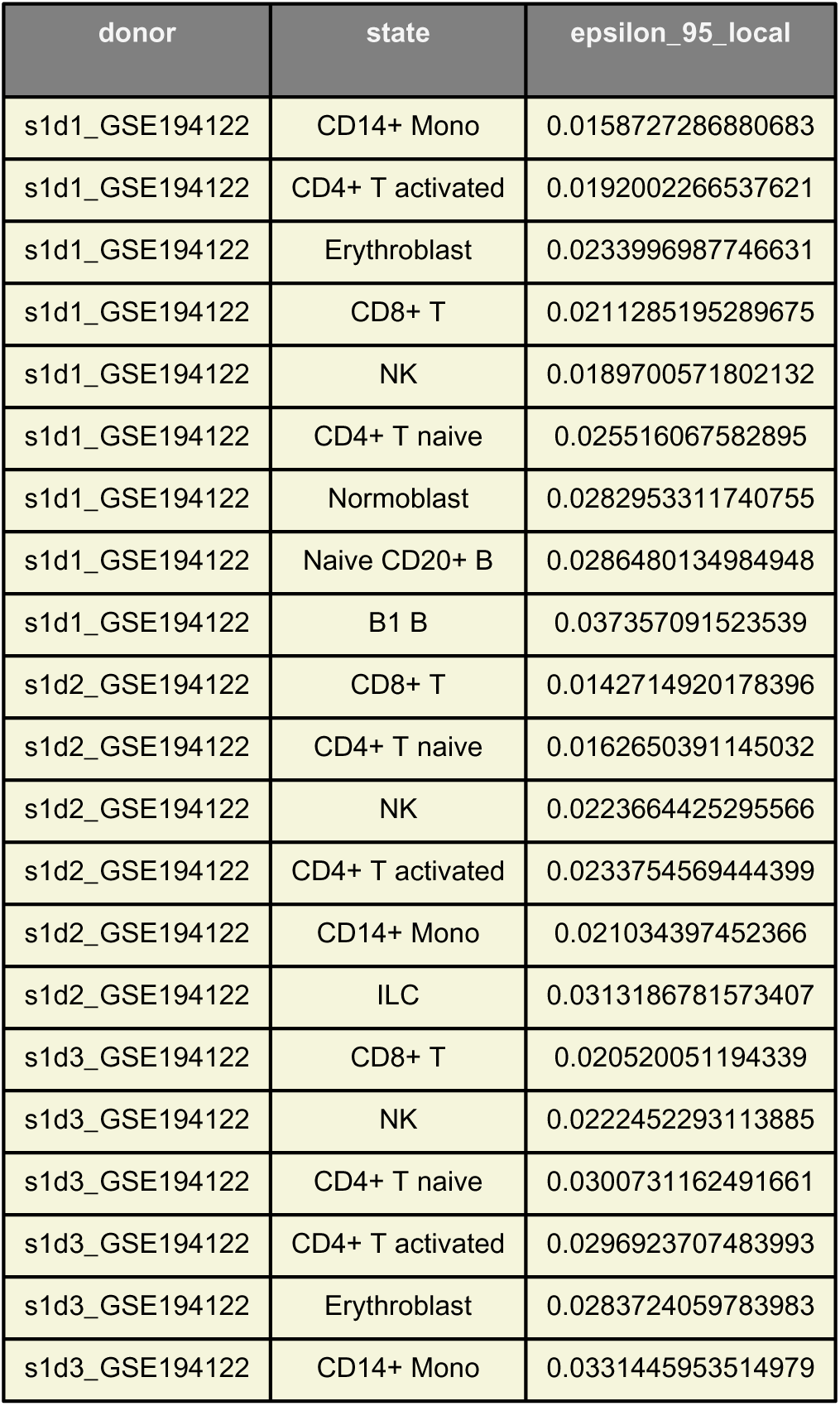
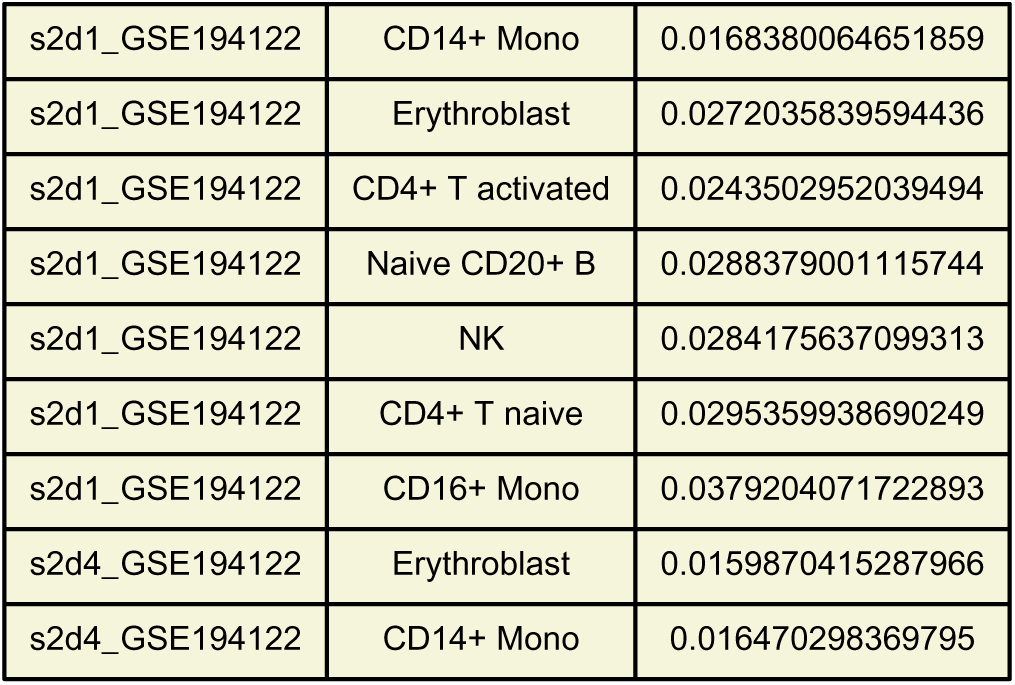
Local MI Results.

**Table S3:**
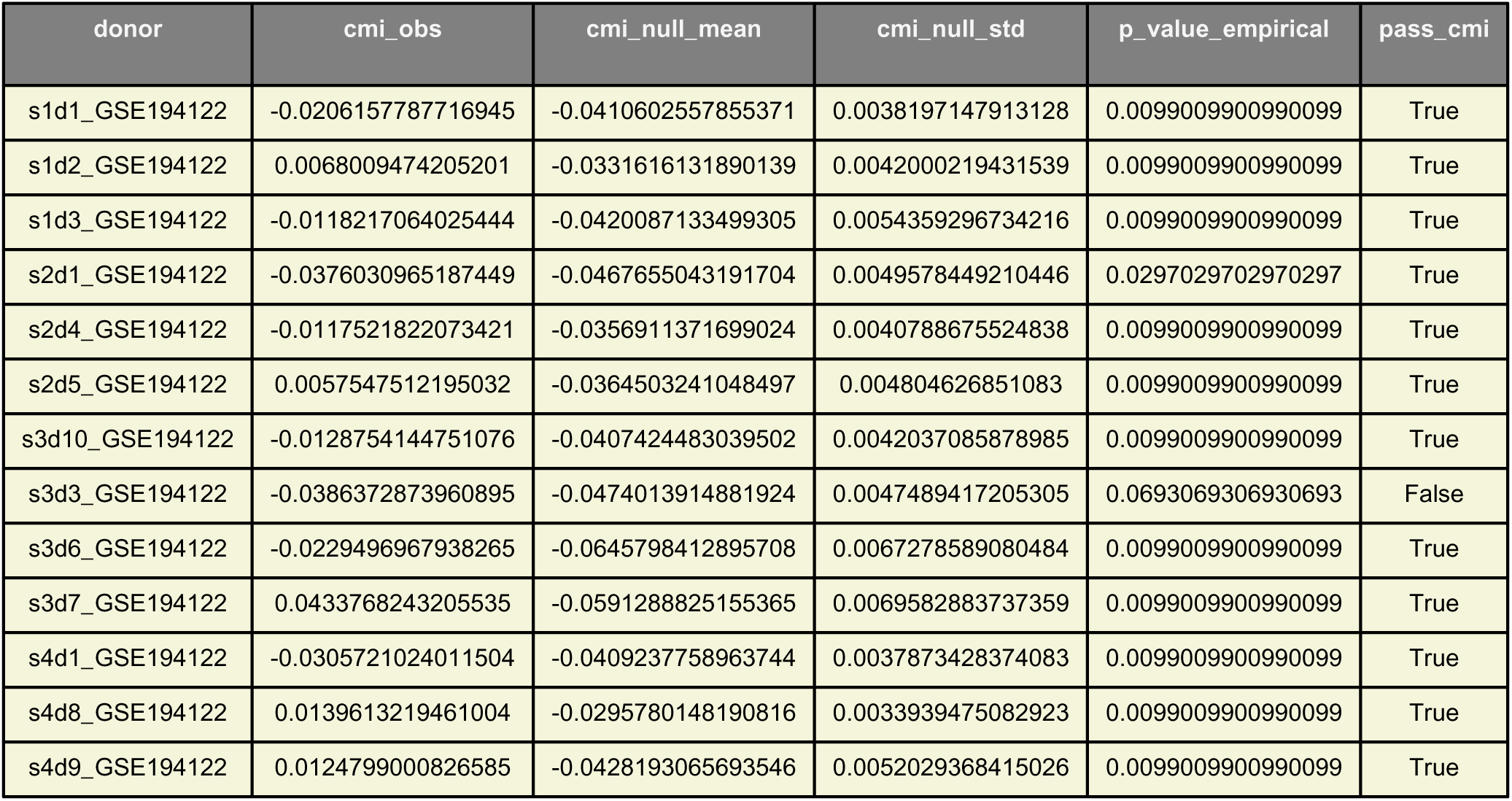
Conditional MI Results.

**Table S4:**
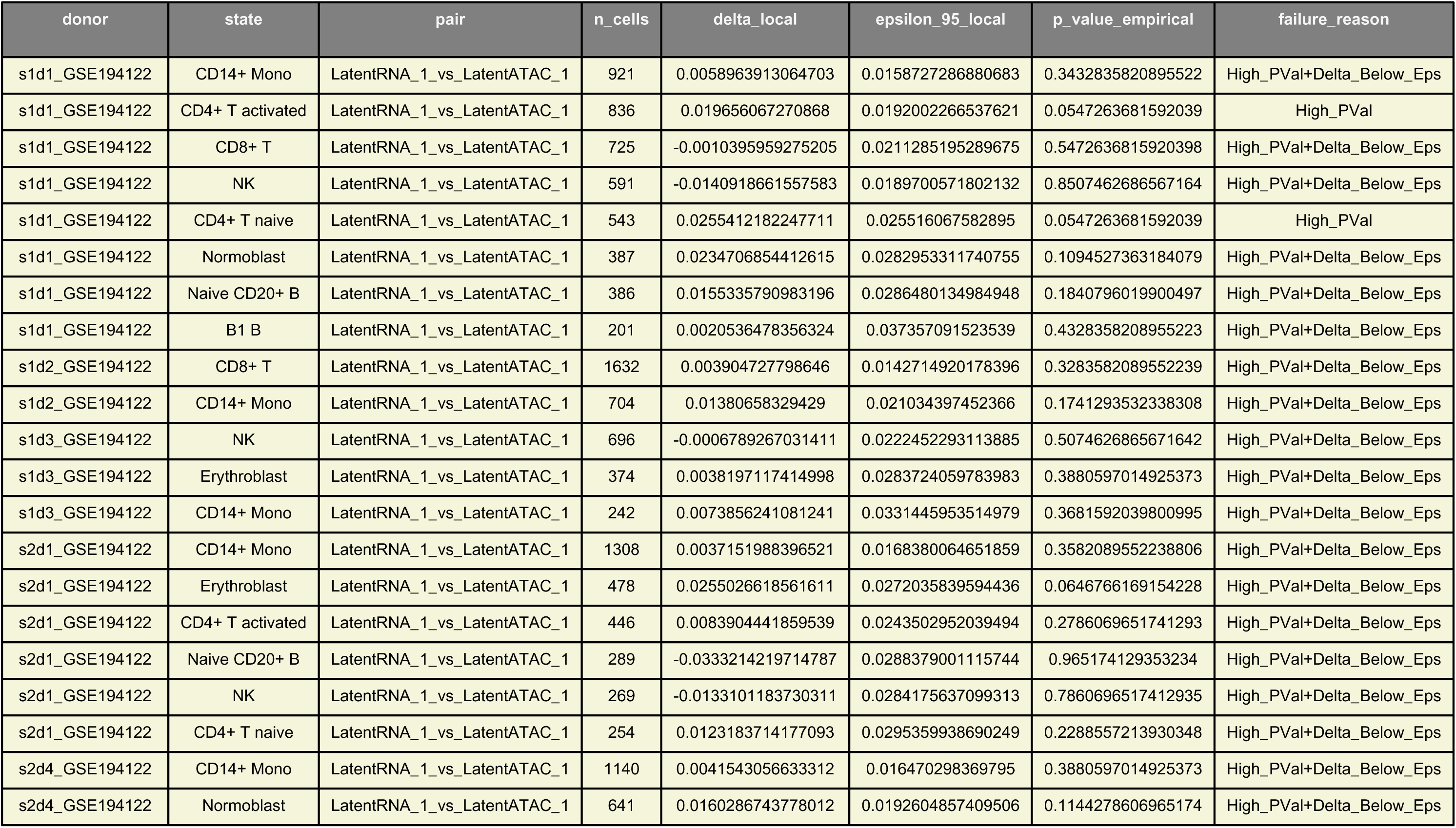
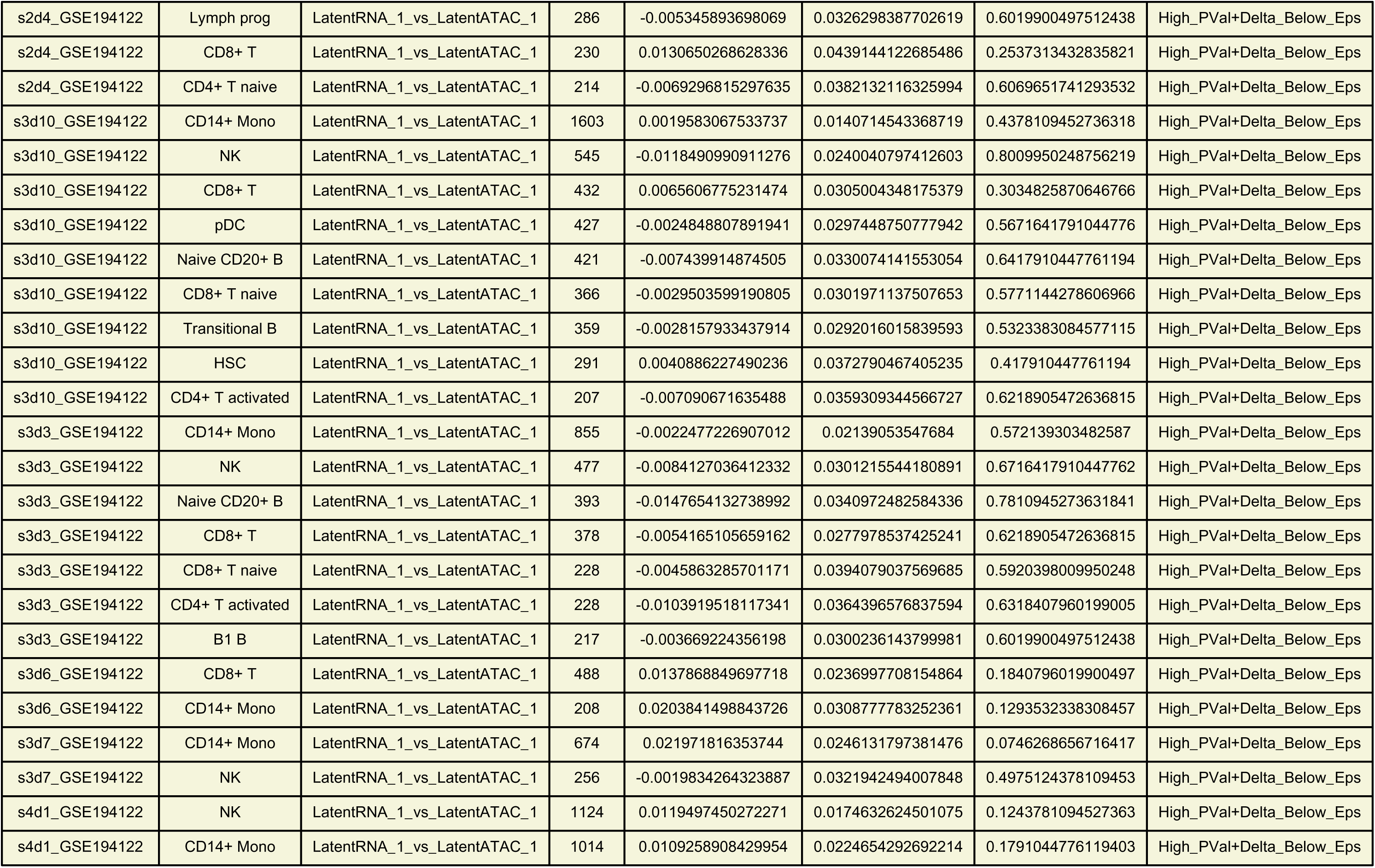
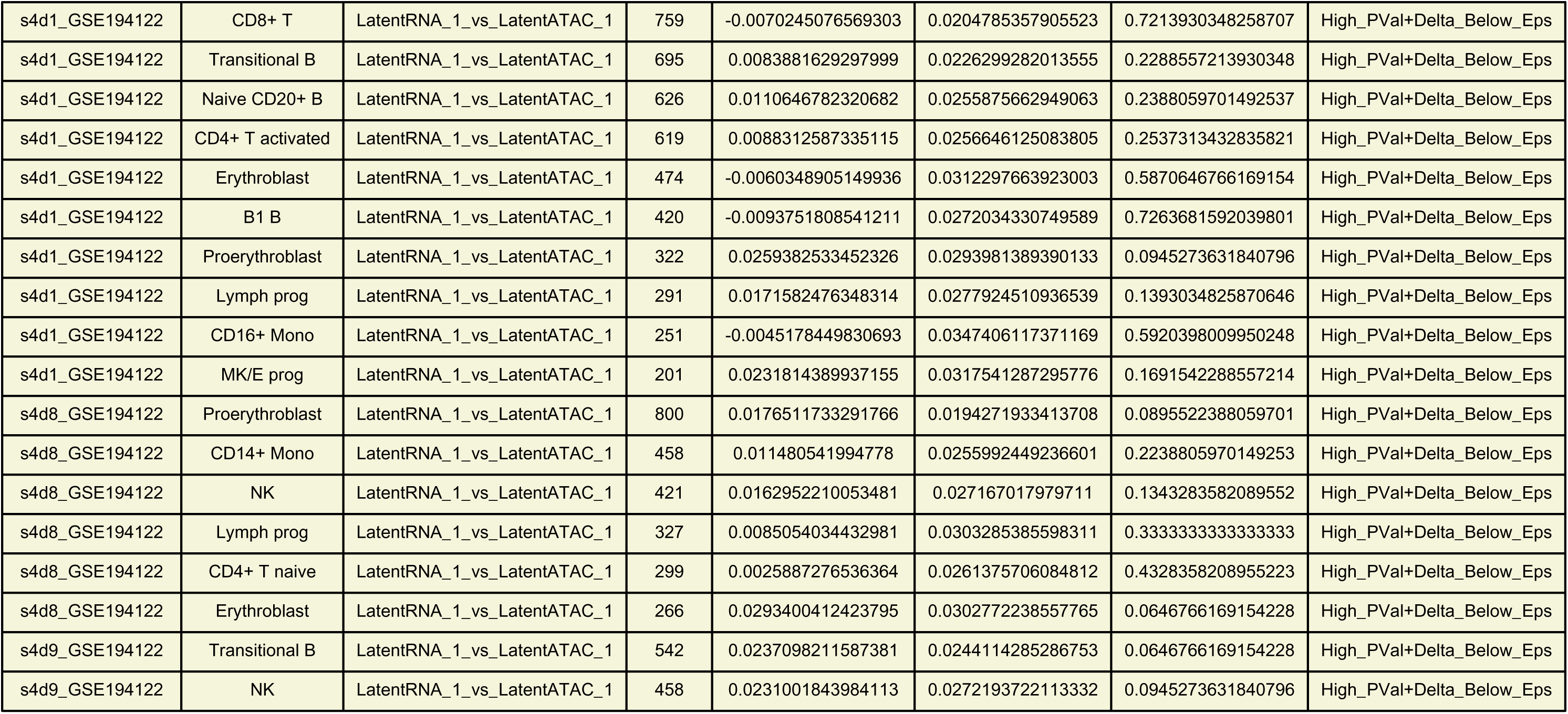
Counterexample Report.

## References

1. Stuart, T. & Satija, R. Integrative single-cell analysis. Nat. Rev. Genet. 20, 257–272 (2019).

2. Ma, S. et al. Chromatin potential identified by shared single-cell profiling of RNA and chromatin. Cell 183, 1103–1116 (2020).

3. Cao, J. et al. Joint profiling of chromatin accessibility and gene expression in thousands of single cells. Science 361, 1380–1385 (2018).

4. Buenrostro, J. D. et al. Single-cell chromatin accessibility reveals principles of regulatory variation. Nature 523, 486–490 (2015).

5. Granja, J. M. et al. ArchR is a scalable software package for integrative single-cell chromatin accessibility analysis. Nat. Genet. 53, 403–411 (2021).

6. Lareau, C. A. et al. Droplet-based combinatorial indexing for massive-scale single-cell chromatin accessibility. Nat. Biotechnol. 37, 916–924 (2019).

7. Stuart, T. et al. Comprehensive integration of single-cell data. Cell 177, 1888–1902 (2019).

15. Kraskov, A., Stögbauer, H. & Grassberger, P. Estimating mutual information. Phys. Rev. E 69, 066138 (2004).

16. Cover, T. M. & Thomas, J. A. Elements of Information Theory (Wiley, 2006).

17. Shannon, C. E. A mathematical theory of communication. Bell Syst. Tech. J. 27, 379–423 (1948).

18. Paninski, L. Estimation of entropy and mutual information. Neural Comput. 15, 1191–1253 (2003).

19. Simpson, E. H. The interpretation of interaction in contingency tables. J. R. Stat. Soc. B 13, 238–241 (1951).

20. Lopez, R., Regier, J., Cole, M. B., Jordan, M. I. & Yosef, N. Deep generative modeling for single-cell transcriptomics. Nat. Methods 15, 1053–1058 (2018).

21. Luecken, M. D. & Theis, F. J. Current best practices in single-cell RNA-seq analysis. Mol. Syst. Biol. 15, e8746 (2019).

22. Trapnell, C. et al. The dynamics and regulators of cell fate decisions are revealed by pseudotemporal ordering of single cells. Nat. Biotechnol. 32, 381–386 (2014).

23. Setty, M. et al. Characterization of cell fate probabilities in single-cell data with Palantir. Nat. Biotechnol. 37, 451–460 (2019).

24. Wolf, F. A., Angerer, P. & Theis, F. J. SCANPY: large-scale single-cell gene expression data analysis. Genome Biol. 19, 15 (2018).

25. van der Maaten, L. & Hinton, G. Visualizing data using t-SNE. J. Mach. Learn. Res. 9, 2579–2605 (2008).

26. McInnes, L., Healy, J. & Melville, J. UMAP: Uniform Manifold Approximation and Projection for dimension reduction. *arXiv* 1802.03426 (2018).

27. The Tabula Sapiens Consortium. The Tabula Sapiens: a multiple-organ, single-cell transcriptomic atlas of humans. Science 376, eabl4896 (2022).

