## Supplementary material for "Hierarchical bounds on RNA–chromatin statistical dependence across cellular states in paired single-cell multiome data": data supplementary: Manifest_PRX_Life_FIXED.pdf

PRX LIFE EXPORT MANIFEST (FINAL TEXT FIX)

Date: 2026-01-07 10:45:07

Target: PRX Life

Bundle Hash: 9730cd228d1d89a48444da777e41f1d77fddfa5912858bfa63d735b65bb1d07d

----------------------------------------

Data_Sources_and_Provenance.pdf |
c236792ede3491ab30817d3822ea4c23c9e98e6d7fb980dab3b6c239ff335c2c

Fig1_Global_MI_vs_Nulls.pdf |
888cb8cddbae890f981fc9177713f8c442aa76bc85a0c4c088b245778acf9e37

Fig2_Local_MI_State_Distributions.pdf |
dc66319d3b9164f981bb614f61a20a2dfeca4674862c5c7b624c2898cb2cf74f

Fig3_Compositional_Decomposition_rho.pdf |
3af45ac93fa2d25c055df62d62930f62806c0829f42a0b02e31c7e241e95d970

Fig4_Conditional_MI_Orthogonal_Test.pdf |
2414b085764723f5a34ec68a021da19019d0c2ef8406ac62110b6540a282dae1

Fig5_Hierarchical_Channel_Summary.pdf |
596b63b103c9b16ae33e822ef9ba841f4536bb25dbec44daba7401c420f581d9

SourceData_Fig1.pdf | 3181f4ad86c9a44bee5a62c3dd21e9871b30fb805f1e64a28a4b86ba8381e9ca

SourceData_Fig2.pdf | 60e9a411eae86d5d53b54c162c093f64b6464d0e70ff19c912591a49b0d3638e

SourceData_Fig3.pdf | 4a047c3bcd0583060c5a16eb133f692f1d0d3ce931a554087018f196f457bb4f

SourceData_Fig4.pdf | 6f6f36481c4031936f7228a092bf266ce2562cd08ce4cd495cde21fcb5649f3c

SourceData_Fig5.pdf | 6275cb04a2ea75aa067c9602dde19833ed6501494afc9f126eddebe4f2ccd7ef

SourceData_SuppFig4.pdf |
ded30456ca09ef8677b4c42098da1d3657a14efd03b8148c135aff4eb6e8cc2a

SuppFig1_Null_Model_Schematics.pdf |
1c4cb4caf81637199f5adea2eafb444d508ec9bac6c720e8ed41f95a643a7ca1

SuppFig2_Epsilon_Calibration_Noise_Floor.pdf |
342534c53224646e2bf87994136847471b52e5875c9c76b8f5e198d3f245d48a

SuppFig3_Permutation_Stability_Checks.pdf |
75f7ce1440957833497067749e2d37ca4174449fb015aae4b0a43112c7459e87

SuppFig4_Donor_Heterogeneity.pdf |
01a0f7a03bcc11040e72bdfcbbecf57beb87b8a41a1fa7fff8cfd3ae62738372

SuppFig5_Estimator_Bias_Sensitivity.pdf |
e14549cd2f95486e5d9936e9808c94547e284fa6658cb7c3b3e259c43fcffef6

SuppFig6_Parameter_Robustness.pdf |
f9a10db4353662e84554e08b3a76dbb3985292edddd01f2efd45d4e16254e02f

SuppFig7_Noise_Floor_and_Epsilon_Validation.pdf |
9cfceb88c82de5c48b5504a3ec5b9676c5278e0fbe0621d656e6fa80882ee506

Supplementary_Methods.pdf |
d436037fba3be331097397e94ef91c95c5dd4b5fbf8187d1a187821424bdf7cc

Supplementary_Notes_and_Counterexamples.pdf |
7789d9d76ed9fc19b55b09f4dbe9c48f4d04b8f6df8912ede7390d96014f4187

TableS1_Global_MI_Summary.pdf |
c46dd781a4b405cfa60009bccf3a8d68e0d59556814fd817f48614db74b6da65

TableS2_Local_MI_and_Hotspot_Classification.pdf |
139b90ee25bea6619f083a0c5579475d5b589e6e528a8c39b881328ffe044ab9

TableS3_Conditional_MI_Results.pdf |
c7259483e69a7a76a06a1a58fa7f4dc92b8a3fafb07a36b6663d7ecef7533418

TableS4_Counterexample_Donor_Report.pdf |
125528123e66fcf9bff7dc5deb4ae840f51034f0cdaa7e9ee563e922a742a4e1
