## Supplementary material for "Hierarchical bounds on RNA–chromatin statistical dependence across cellular states in paired single-cell multiome data": data supplementary: SourceData_Fig1.pdf

### Source Data Fig 1: Global MI

Derived from: RC3_R2_STRICT_BUNDLE (SHA256: 9730cd228d1d89a48444da777e41f1d77fddfa5912858bfa63d735b65bb1d07d)
Provenance: See Manifest_PRX_Life_FIXED.pdf

| donor | pair | mi_obs | mi_null_C | delta_C |
|---|---|---|---|---|
| s1d1_GSE194122 | LatentRNA_1_vs_LatentATAC_1 | 0.4770899410054898 | 0.0116073674014753 | 0.4654825736040145 |
| s1d1_GSE194122 | LatentRNA_1_vs_LatentATAC_2 | 0.3865310481324542 | 0.0227339310492883 | 0.3637971170831658 |
| s1d1_GSE194122 | LatentRNA_2_vs_LatentATAC_1 | 0.7954205148541718 | 0.0288993060208073 | 0.7665212088333644 |
| s1d2_GSE194122 | LatentRNA_1_vs_LatentATAC_1 | 0.6720277345130619 | 0.0092741359018675 | 0.6627535986111943 |
| s1d2_GSE194122 | LatentRNA_1_vs_LatentATAC_2 | 0.5313480151275289 | 0.0005518255746147 | 0.5307961895529142 |
| s1d2_GSE194122 | LatentRNA_2_vs_LatentATAC_1 | 0.1934848700419618 | 7.455543555977329e-05 | 0.193410314606402 |
| s1d3_GSE194122 | LatentRNA_1_vs_LatentATAC_1 | 0.4796373202880275 | 0.0144946189928765 | 0.4651427012951509 |
| s1d3_GSE194122 | LatentRNA_1_vs_LatentATAC_2 | 0.4488119940026465 | 0.0016880453835934 | 0.447123948619053 |
| s1d3_GSE194122 | LatentRNA_2_vs_LatentATAC_1 | 0.4506462022952568 | 0.0253766631272787 | 0.4252695391679781 |
| s2d1_GSE194122 | LatentRNA_1_vs_LatentATAC_1 | 0.7042765629868466 | 0.0113423872377012 | 0.6929341757491453 |
| s2d1_GSE194122 | LatentRNA_1_vs_LatentATAC_2 | 0.7696000731743933 | 0.0075176553252232 | 0.7620824178491701 |
| s2d1_GSE194122 | LatentRNA_2_vs_LatentATAC_1 | 0.5533519302838705 | 0.0109220261294864 | 0.5424299041543841 |
| s2d4_GSE194122 | LatentRNA_1_vs_LatentATAC_1 | 0.6609554473027739 | 0.0101923147626947 | 0.6507631325400792 |
| s2d4_GSE194122 | LatentRNA_1_vs_LatentATAC_2 | 0.3850660778738799 | 0.0189847420995093 | 0.3660813357743706 |
| s2d4_GSE194122 | LatentRNA_2_vs_LatentATAC_1 | 0.5108452402386341 | 0.0041398717568029 | 0.5067053684818312 |
| s2d5_GSE194122 | LatentRNA_1_vs_LatentATAC_1 | 0.7737890031661312 | 0.082189079023615 | 0.6915999241425161 |
| s2d5_GSE194122 | LatentRNA_1_vs_LatentATAC_2 | 0.3053209806000367 | 0.0001006403716765 | 0.3052203402283602 |
| s2d5_GSE194122 | LatentRNA_2_vs_LatentATAC_1 | 0.364032182932446 | 0.0391682419239702 | 0.3248639410084757 |
| s3d10_GSE194122 | LatentRNA_1_vs_LatentATAC_1 | 0.4412969666281281 | 0.0098326430769969 | 0.4314643235511311 |
| s3d10_GSE194122 | LatentRNA_1_vs_LatentATAC_2 | 0.3426697491466868 | 0.0005897069233766 | 0.3420800422233102 |

| donor | pair | mi_obs | mi_null_C | delta_C |
| --- | --- | --- | --- | --- |
| s3d10_GSE194122 | LatentRNA_2_vs_LatentATAC_1 | 0.5526936207345621 | 0.0111702715381907 | 0.5415233491963714 |
| s3d3_GSE194122 | LatentRNA_1_vs_LatentATAC_1 | 0.2446186945277322 | 0.0003968007600682 | 0.244221893767664 |
| s3d3_GSE194122 | LatentRNA_1_vs_LatentATAC_2 | 0.4192584925247242 | 0.0 | 0.4192584925247242 |
| s3d3_GSE194122 | LatentRNA_2_vs_LatentATAC_1 | 0.6596542648208761 | 0.0051309732358798 | 0.6545232915849962 |
| s3d6_GSE194122 | LatentRNA_1_vs_LatentATAC_1 | 0.7322491656751478 | 0.0529908107236222 | 0.6792583549515255 |
| s3d6_GSE194122 | LatentRNA_1_vs_LatentATAC_2 | 0.592476380950405 | 0.0129583723879662 | 0.5795180085624387 |
| s3d6_GSE194122 | LatentRNA_2_vs_LatentATAC_1 | 0.5950571402437301 | 0.0263888359393457 | 0.5686683043043844 |
| s3d7_GSE194122 | LatentRNA_1_vs_LatentATAC_1 | 0.7168729617145235 | 0.0497016887252323 | 0.6671712729892911 |
| s3d7_GSE194122 | LatentRNA_1_vs_LatentATAC_2 | 0.3092787819607991 | 0.0151957234003849 | 0.2940830585604141 |
| s3d7_GSE194122 | LatentRNA_2_vs_LatentATAC_1 | 0.4255454781346941 | 0.0475683970309012 | 0.3779770811037928 |
| s4d1_GSE194122 | LatentRNA_1_vs_LatentATAC_1 | 0.1764713819668024 | 0.000141484567579 | 0.1763298973992233 |
| s4d1_GSE194122 | LatentRNA_1_vs_LatentATAC_2 | 0.097045310734682 | 0.0005524974177833 | 0.0964928133168986 |
| s4d1_GSE194122 | LatentRNA_2_vs_LatentATAC_1 | 0.3763713823050718 | 0.0001792241163237 | 0.3761921581887481 |
| s4d8_GSE194122 | LatentRNA_1_vs_LatentATAC_1 | 0.6952741568805045 | 0.039971440969002 | 0.6553027159115025 |
| s4d8_GSE194122 | LatentRNA_1_vs_LatentATAC_2 | 0.7501918590646053 | 0.0400501315681838 | 0.7101417274964215 |
| s4d8_GSE194122 | LatentRNA_2_vs_LatentATAC_1 | 0.4809791651288062 | 0.035243353410442 | 0.4457358117183642 |
| s4d9_GSE194122 | LatentRNA_1_vs_LatentATAC_1 | 0.6865760266133343 | 0.0102437026093912 | 0.6763323240039429 |
| s4d9_GSE194122 | LatentRNA_1_vs_LatentATAC_2 | 0.3787786900681738 | 0.00225565565282 | 0.3765230344153537 |
| s4d9_GSE194122 | LatentRNA_2_vs_LatentATAC_1 | 0.3874667096553281 | 0.0039628785761272 | 0.3835038310792009 |
