## Supplementary material for "Hierarchical bounds on RNA–chromatin statistical dependence across cellular states in paired single-cell multiome data": data supplementary: SourceData_Fig2.pdf

### Source Data Fig 2: Local MI

Derived from: RC3_R2_STRICT_BUNDLE (SHA256: 9730cd228d1d89a48444da777e41f1d77fddfa5912858bfa63d735b65bb1d07d)
Provenance: See Manifest_PRX_Life_FIXED.pdf

| donor | state | pair | n_cells | mi_obs_raw | delta_local | epsilon_95_local | epsilon_99_local | p_value_empirical | pass_local_95 | pass_local_99 |
|---|---|---|---|---|---|---|---|---|---|---|
| s1d1_GSE194122 | CD14+ Mono | LatentRNA_1_vs_LatentATAC_1 | 921 | -0.0291213873080469 | 0.0058963913064703 | 0.0158727286880683 | 0.0232356177068574 | 0.3432835820895522 | False | False |
| s1d1_GSE194122 | CD4+ T activated | LatentRNA_1_vs_LatentATAC_1 | 836 | -0.0051192565634057 | 0.019656067270868 | 0.0192002266537621 | 0.027666450028539 | 0.0547263681592039 | False | False |
| s1d1_GSE194122 | Erythroblast | LatentRNA_1_vs_LatentATAC_1 | 797 | 0.0324123394836064 | 0.0571971925443421 | 0.0233996987746631 | 0.0344288145259887 | 0.0049751243781094 | True | True |
| s1d1_GSE194122 | CD8+ T | LatentRNA_1_vs_LatentATAC_1 | 725 | -0.0316461015668689 | -0.0010395959275205 | 0.0211285195289675 | 0.0255522307603548 | 0.5472636815920398 | False | False |
| s1d1_GSE194122 | NK | LatentRNA_1_vs_LatentATAC_1 | 591 | -0.0445038690438899 | -0.0140918661557583 | 0.0189700571802132 | 0.0284637454262492 | 0.8507462686567164 | False | False |
| s1d1_GSE194122 | CD4+ T naive | LatentRNA_1_vs_LatentATAC_1 | 543 | -0.0073473323330954 | 0.0255412182247711 | 0.025516067582895 | 0.0366343326125093 | 0.0547263681592039 | False | False |
| s1d1_GSE194122 | Normoblast | LatentRNA_1_vs_LatentATAC_1 | 387 | -0.0112426886881014 | 0.0234706854412615 | 0.0282953311740755 | 0.0399072975150445 | 0.1094527363184079 | False | False |
| s1d1_GSE194122 | Naive CD20+ B | LatentRNA_1_vs_LatentATAC_1 | 386 | -0.0245534250666299 | 0.0155335790983196 | 0.0286480134984948 | 0.0392159935554 | 0.1840796019900497 | False | False |
| s1d1_GSE194122 | B1 B | LatentRNA_1_vs_LatentATAC_1 | 201 | -0.0520384160666465 | 0.0020536478356324 | 0.037357091523539 | 0.0477837257219266 | 0.4328358208955223 | False | False |
| s1d2_GSE194122 | CD8+ T | LatentRNA_1_vs_LatentATAC_1 | 1632 | -0.0148293769735783 | 0.003904727798646 | 0.0142714920178396 | 0.0240487094356241 | 0.3283582089552239 | False | False |
| s1d2_GSE194122 | CD4+ T naive | LatentRNA_1_vs_LatentATAC_1 | 1289 | 0.0361154670084662 | 0.0584803721981296 | 0.0162650391145032 | 0.0271808520945239 | 0.0049751243781094 | True | True |
| s1d2_GSE194122 | NK | LatentRNA_1_vs_LatentATAC_1 | 982 | 0.0251342520795034 | 0.0504561385818323 | 0.0223664425295566 | 0.0307329094628588 | 0.0049751243781094 | True | True |
| s1d2_GSE194122 | CD4+ T activated | LatentRNA_1_vs_LatentATAC_1 | 709 | 0.0086966179016521 | 0.0356807419175797 | 0.0233754569444399 | 0.0337478953264586 | 0.0149253731343283 | True | False |
| s1d2_GSE194122 | CD14+ Mono | LatentRNA_1_vs_LatentATAC_1 | 704 | -0.0127896567431573 | 0.01380658329429 | 0.021034397452366 | 0.0327526899419872 | 0.1741293532338308 | False | False |
| s1d2_GSE194122 | ILC | LatentRNA_1_vs_LatentATAC_1 | 367 | 0.0123391535470744 | 0.0523786497373382 | 0.0313186781573407 | 0.042646093379662 | 0.0099502487562189 | True | True |
| s1d3_GSE194122 | CD8+ T | LatentRNA_1_vs_LatentATAC_1 | 1020 | 0.00309351484856 | 0.0276371331327552 | 0.020520051194339 | 0.0268802878750212 | 0.0099502487562189 | True | True |
| s1d3_GSE194122 | NK | LatentRNA_1_vs_LatentATAC_1 | 696 | -0.0294261882398263 | -0.0006789267031411 | 0.0222452293113885 | 0.0267299798230972 | 0.5074626865671642 | False | False |
| s1d3_GSE194122 | CD4+ T naive | LatentRNA_1_vs_LatentATAC_1 | 533 | 0.0400517409623262 | 0.0721476895746775 | 0.0300731162491661 | 0.0410450539732789 | 0.0049751243781094 | True | True |
| s1d3_GSE194122 | CD4+ T activated | LatentRNA_1_vs_LatentATAC_1 | 392 | 0.0215157024876297 | 0.0573897693943039 | 0.0296923707483993 | 0.0453078168442587 | 0.0049751243781094 | True | True |
| s1d3_GSE194122 | Erythroblast | LatentRNA_1_vs_LatentATAC_1 | 374 | -0.0359113648141455 | 0.0038197117414998 | 0.0283724059783983 | 0.0509633910350439 | 0.3880597014925373 | False | False |

| donor | state | pair | n_cells | mi_obs_raw | delta_local | epsilon_95_local | epsilon_99_local | p_value_empirical | pass_local_95 | pass_local_99 |
|---|---|---|---|---|---|---|---|---|---|---|
| s1d3_GSE194122 | CD14+ Mono | LatentRNA_1_vs_LatentATAC_1 | 242 | -0.0445483472586181 | 0.0073856241081241 | 0.0331445953514979 | 0.0430440718735279 | 0.3681592039800995 | False | False |
| s2d1_GSE194122 | CD14+ Mono | LatentRNA_1_vs_LatentATAC_1 | 1308 | -0.0217698492441957 | 0.0037151988396521 | 0.0168380064651859 | 0.023631222368302 | 0.3582089552238806 | False | False |
| s2d1_GSE194122 | Erythroblast | LatentRNA_1_vs_LatentATAC_1 | 478 | -0.0049634592208107 | 0.0255026618561611 | 0.0272035839594436 | 0.0390909593085538 | 0.0646766169154228 | False | False |
| s2d1_GSE194122 | CD4+ T activated | LatentRNA_1_vs_LatentATAC_1 | 446 | -0.0324990354813987 | 0.0083904441859539 | 0.0243502952039494 | 0.0350083441751199 | 0.2786069651741293 | False | False |
| s2d1_GSE194122 | Naive CD20+ B | LatentRNA_1_vs_LatentATAC_1 | 289 | -0.0783455690859753 | -0.0333214219714787 | 0.0288379001115744 | 0.038007349721517 | 0.965174129353234 | False | False |
| s2d1_GSE194122 | NK | LatentRNA_1_vs_LatentATAC_1 | 269 | -0.0655684267584417 | -0.0133101183730311 | 0.0284175637099313 | 0.0447089063185658 | 0.7860696517412935 | False | False |
| s2d1_GSE194122 | CD4+ T naive | LatentRNA_1_vs_LatentATAC_1 | 254 | -0.0437054053322167 | 0.0123183714177093 | 0.0295359938690249 | 0.0458889090766395 | 0.2288557213930348 | False | False |
| s2d1_GSE194122 | CD16+ Mono | LatentRNA_1_vs_LatentATAC_1 | 209 | -0.0115665867145668 | 0.0418869296034295 | 0.0379204071722893 | 0.054813321122568 | 0.044776119402985 | True | False |
| s2d4_GSE194122 | Erythroblast | LatentRNA_1_vs_LatentATAC_1 | 1485 | 0.0108185653812604 | 0.0301781341192011 | 0.0159870415287966 | 0.0267271980799288 | 0.0099502487562189 | True | True |
| s2d4_GSE194122 | CD14+ Mono | LatentRNA_1_vs_LatentATAC_1 | 1140 | -0.0230352826719464 | 0.0041543056633312 | 0.016470298369795 | 0.0240398408277137 | 0.3880597014925373 | False | False |
| s2d4_GSE194122 | Normoblast | LatentRNA_1_vs_LatentATAC_1 | 641 | -0.0144559796520109 | 0.0160286743778012 | 0.0192604857409506 | 0.0277639804987392 | 0.1144278606965174 | False | False |
| s2d4_GSE194122 | Naive CD20+ B | LatentRNA_1_vs_LatentATAC_1 | 590 | -0.005216829824354 | 0.0297345336271969 | 0.0215673054492205 | 0.0261346157644645 | 0.0099502487562189 | True | True |
| s2d4_GSE194122 | CD4+ T activated | LatentRNA_1_vs_LatentATAC_1 | 385 | -0.0020039754039444 | 0.0354246694972804 | 0.0274848156481466 | 0.0418210898451154 | 0.0298507462686567 | True | False |
| s2d4_GSE194122 | Proerythroblast | LatentRNA_1_vs_LatentATAC_1 | 316 | 0.0652604415736961 | 0.1076374631073629 | 0.0287909301824201 | 0.0397727438883871 | 0.0049751243781094 | True | True |
| s2d4_GSE194122 | Lymph prog | LatentRNA_1_vs_LatentATAC_1 | 286 | -0.0485380010014848 | -0.005345893698069 | 0.0326298387702619 | 0.0472162718579103 | 0.6019900497512438 | False | False |
| s2d4_GSE194122 | CD8+ T | LatentRNA_1_vs_LatentATAC_1 | 230 | -0.035101464559295 | 0.0130650268628336 | 0.0439144122685486 | 0.0562118840895689 | 0.2537313432835821 | False | False |
| s2d4_GSE194122 | CD4+ T naive | LatentRNA_1_vs_LatentATAC_1 | 214 | -0.0544753219319789 | -0.0069296815297635 | 0.0382132116325994 | 0.0537698134829975 | 0.6069651741293532 | False | False |
| s2d5_GSE194122 | CD14+ Mono | LatentRNA_1_vs_LatentATAC_1 | 1716 | 0.0254931042577073 | 0.045006704484421 | 0.0156217428220245 | 0.0183888651245434 | 0.0049751243781094 | True | True |
| s2d5_GSE194122 | NK | LatentRNA_1_vs_LatentATAC_1 | 848 | -0.0067773439996958 | 0.0240231160441952 | 0.0206524165614946 | 0.0319867684337676 | 0.0298507462686567 | True | False |
| s2d5_GSE194122 | CD8+ T | LatentRNA_1_vs_LatentATAC_1 | 752 | -0.0072778646135933 | 0.0265025670314771 | 0.0196641506639271 | 0.0338078236897606 | 0.0248756218905472 | True | False |
| s2d5_GSE194122 | CD4+ T activated | LatentRNA_1_vs_LatentATAC_1 | 493 | 0.0082323581021075 | 0.047586938186787 | 0.0288192542859541 | 0.0380776407854359 | 0.0049751243781094 | True | True |
| s2d5_GSE194122 | CD16+ Mono | LatentRNA_1_vs_LatentATAC_1 | 380 | 0.0402452167015621 | 0.0790496768598454 | 0.026864000332501 | 0.0462449564709814 | 0.0049751243781094 | True | True |
| s3d10_GSE194122 | CD14+ Mono | LatentRNA_1_vs_LatentATAC_1 | 1603 | -0.0220612683436645 | 0.0019583067533737 | 0.0140714543368719 | 0.017160934984803 | 0.4378109452736318 | False | False |
| s3d10_GSE194122 | NK | LatentRNA_1_vs_LatentATAC_1 | 545 | -0.048871545966822 | -0.0118490990911276 | 0.0240040797412603 | 0.0375654418855417 | 0.8009950248756219 | False | False |

| donor | state | pair | n_cells | mi_obs_raw | delta_local | epsilon_95_local | epsilon_99_local | p_value_empirical | pass_local_95 | pass_local_99 |
|---|---|---|---|---|---|---|---|---|---|---|
| s3d10_GSE194122 | G/M prog | LatentRNA_1_vs_LatentATAC_1 | 432 | 0.1893370983297284 | 0.2350204738139081 | 0.0259990955053132 | 0.0427636267954583 | 0.0049751243781094 | True | True |
| s3d10_GSE194122 | CD8+ T | LatentRNA_1_vs_LatentATAC_1 | 432 | -0.0348970346949943 | 0.0065606775231474 | 0.0305004348175379 | 0.0375653525024934 | 0.3034825870646766 | False | False |
| s3d10_GSE194122 | pDC | LatentRNA_1_vs_LatentATAC_1 | 427 | -0.0407567940256683 | -0.0024848807891941 | 0.0297448750777942 | 0.0382120554989439 | 0.5671641791044776 | False | False |
| s3d10_GSE194122 | Naive CD20+ B | LatentRNA_1_vs_LatentATAC_1 | 421 | -0.0458960958460767 | -0.007439914874505 | 0.0330074141553054 | 0.0374826607354763 | 0.6417910447761194 | False | False |
| s3d10_GSE194122 | Erythroblast | LatentRNA_1_vs_LatentATAC_1 | 391 | 0.0182677391777357 | 0.0543282245426582 | 0.0283259428008225 | 0.0376818384712029 | 0.0099502487562189 | True | True |
| s3d10_GSE194122 | CD8+ T naive | LatentRNA_1_vs_LatentATAC_1 | 366 | -0.0484982891887195 | -0.0029503599190805 | 0.0301971137507653 | 0.042751303000131 | 0.5771144278606966 | False | False |
| s3d10_GSE194122 | Transitional B | LatentRNA_1_vs_LatentATAC_1 | 359 | -0.0438170616066546 | -0.0028157933437914 | 0.0292016015839593 | 0.0414232303611271 | 0.5323383084577115 | False | False |
| s3d10_GSE194122 | HSC | LatentRNA_1_vs_LatentATAC_1 | 291 | -0.0396815965304906 | 0.0040886227490236 | 0.0372790467405235 | 0.057798771697256 | 0.417910447761194 | False | False |
| s3d10_GSE194122 | CD16+ Mono | LatentRNA_1_vs_LatentATAC_1 | 253 | 0.0273768966363636 | 0.0849664811376176 | 0.0262016429079498 | 0.0462365493963733 | 0.0049751243781094 | True | True |
| s3d10_GSE194122 | Lymph prog | LatentRNA_1_vs_LatentATAC_1 | 231 | 0.0303555219092226 | 0.0815920322402563 | 0.0325816794203854 | 0.0462734122664219 | 0.0049751243781094 | True | True |
| s3d10_GSE194122 | CD4+ T activated | LatentRNA_1_vs_LatentATAC_1 | 207 | -0.0621089673176404 | -0.007090671635488 | 0.0359309344566727 | 0.0474337783040397 | 0.6218905472636815 | False | False |
| s3d3_GSE194122 | CD14+ Mono | LatentRNA_1_vs_LatentATAC_1 | 855 | -0.0328856768898386 | -0.0022477226907012 | 0.02139053547684 | 0.0254239214756386 | 0.572139303482587 | False | False |
| s3d3_GSE194122 | NK | LatentRNA_1_vs_LatentATAC_1 | 477 | -0.0432401102951622 | -0.0084127036412332 | 0.0301215544180891 | 0.0388726305624658 | 0.6716417910447762 | False | False |
| s3d3_GSE194122 | Naive CD20+ B | LatentRNA_1_vs_LatentATAC_1 | 393 | -0.0519948929084659 | -0.0147654132738992 | 0.0340972482584336 | 0.0431562012809168 | 0.7810945273631841 | False | False |
| s3d3_GSE194122 | CD8+ T | LatentRNA_1_vs_LatentATAC_1 | 378 | -0.0450926823255812 | -0.0054165105659162 | 0.0277978537425241 | 0.0498360301451042 | 0.6218905472636815 | False | False |
| s3d3_GSE194122 | CD4+ T naive | LatentRNA_1_vs_LatentATAC_1 | 319 | -0.0057157932782168 | 0.0397096428986613 | 0.0305895056433747 | 0.0396050374671668 | 0.0149253731343283 | True | False |
| s3d3_GSE194122 | Erythroblast | LatentRNA_1_vs_LatentATAC_1 | 246 | 0.1108797067895297 | 0.1587765075600812 | 0.0331903823639296 | 0.0498041627152889 | 0.0049751243781094 | True | True |
| s3d3_GSE194122 | CD8+ T naive | LatentRNA_1_vs_LatentATAC_1 | 228 | -0.0555374424148418 | -0.0045863285701171 | 0.0394079037569685 | 0.0471271810099293 | 0.5920398009950248 | False | False |
| s3d3_GSE194122 | CD4+ T activated | LatentRNA_1_vs_LatentATAC_1 | 228 | -0.0625077232037529 | -0.0103919518117341 | 0.0364396576837594 | 0.0450504778355428 | 0.6318407960199005 | False | False |
| s3d3_GSE194122 | B1 B | LatentRNA_1_vs_LatentATAC_1 | 217 | -0.0560530249191675 | -0.003669224356198 | 0.0300236143799981 | 0.0475499631469299 | 0.6019900497512438 | False | False |
| s3d6_GSE194122 | CD8+ T | LatentRNA_1_vs_LatentATAC_1 | 488 | -0.022957146428741 | 0.0137868849697718 | 0.0236997708154864 | 0.0360565552914745 | 0.1840796019900497 | False | False |
| s3d6_GSE194122 | CD4+ T activated | LatentRNA_1_vs_LatentATAC_1 | 276 | 0.0577481187705961 | 0.1012549578012364 | 0.035497042071461 | 0.0449939363715526 | 0.0049751243781094 | True | True |
| s3d6_GSE194122 | CD14+ Mono | LatentRNA_1_vs_LatentATAC_1 | 208 | -0.0361194280473391 | 0.0203841498843726 | 0.0308777783252361 | 0.0452471489258253 | 0.1293532338308457 | False | False |
| s3d7_GSE194122 | CD14+ Mono | LatentRNA_1_vs_LatentATAC_1 | 674 | -0.0068651125479686 | 0.021971816353744 | 0.0246131797381476 | 0.0349431881163628 | 0.0746268656716417 | False | False |

| donor | state | pair | n_cells | mi_obs_raw | delta_local | epsilon_95_local | epsilon_99_local | p_value_empirical | pass_local_95 | pass_local_99 |
|---|---|---|---|---|---|---|---|---|---|---|
| s3d7_GSE194122 | NK | LatentRNA_1_vs_LatentATAC_1 | 256 | -0.0535607862731062 | -0.0019834264323887 | 0.0321942494007848 | 0.0482576574267862 | 0.4975124378109453 | False | False |
| s4d1_GSE194122 | NK | LatentRNA_1_vs_LatentATAC_1 | 1124 | -0.0189542615097515 | 0.0119497450272271 | 0.0174632624501075 | 0.0235747224404012 | 0.1243781094527363 | False | False |
| s4d1_GSE194122 | CD14+ Mono | LatentRNA_1_vs_LatentATAC_1 | 1014 | -0.0182357155889043 | 0.0109258908429954 | 0.0224654292692214 | 0.0287914866170699 | 0.1791044776119403 | False | False |
| s4d1_GSE194122 | CD8+ T | LatentRNA_1_vs_LatentATAC_1 | 759 | -0.042593256423931 | -0.0070245076569303 | 0.0204785357905523 | 0.0271432115189539 | 0.7213930348258707 | False | False |
| s4d1_GSE194122 | Transitional B | LatentRNA_1_vs_LatentATAC_1 | 695 | -0.0271523585687276 | 0.0083881629297999 | 0.0226299282013555 | 0.032707579627339 | 0.2288557213930348 | False | False |
| s4d1_GSE194122 | Naive CD20+ B | LatentRNA_1_vs_LatentATAC_1 | 626 | -0.0225441548493829 | 0.0110646782320682 | 0.0255875662949063 | 0.0316342067886206 | 0.2388059701492537 | False | False |
| s4d1_GSE194122 | CD4+ T activated | LatentRNA_1_vs_LatentATAC_1 | 619 | -0.0298078680945135 | 0.0088312587335115 | 0.0256646125083805 | 0.0311157795506284 | 0.2537313432835821 | False | False |
| s4d1_GSE194122 | Erythroblast | LatentRNA_1_vs_LatentATAC_1 | 474 | -0.0369726981360312 | -0.0060348905149936 | 0.0312297663923003 | 0.0392152951297757 | 0.5870646766169154 | False | False |
| s4d1_GSE194122 | B1 B | LatentRNA_1_vs_LatentATAC_1 | 420 | -0.0490840360903614 | -0.0093751808541211 | 0.0272034330749589 | 0.0356520373877563 | 0.7263681592039801 | False | False |
| s4d1_GSE194122 | CD4+ T naive | LatentRNA_1_vs_LatentATAC_1 | 401 | 0.011014864583144 | 0.0580686128555332 | 0.0243072112771641 | 0.0397888111220861 | 0.0049751243781094 | True | True |
| s4d1_GSE194122 | Proerythroblast | LatentRNA_1_vs_LatentATAC_1 | 322 | -0.0128004092022351 | 0.0259382533452326 | 0.0293981389390133 | 0.0452761864155311 | 0.0945273631840796 | False | False |
| s4d1_GSE194122 | Lymph prog | LatentRNA_1_vs_LatentATAC_1 | 291 | -0.043533601320111 | 0.0171582476348314 | 0.0277924510936539 | 0.0393676715345792 | 0.1393034825870646 | False | False |
| s4d1_GSE194122 | CD16+ Mono | LatentRNA_1_vs_LatentATAC_1 | 251 | -0.06780809732094 | -0.0045178449830693 | 0.0347406117371169 | 0.0488471837807897 | 0.5920398009950248 | False | False |
| s4d1_GSE194122 | MK/E prog | LatentRNA_1_vs_LatentATAC_1 | 201 | -0.0278342766456427 | 0.0231814389937155 | 0.0317541287295776 | 0.0431297133309417 | 0.1691542288557214 | False | False |
| s4d8_GSE194122 | CD8+ T | LatentRNA_1_vs_LatentATAC_1 | 4089 | 0.0371259832687496 | 0.0488085733873511 | 0.0100035685426326 | 0.0140880433253696 | 0.0049751243781094 | True | True |
| s4d8_GSE194122 | Naive CD20+ B | LatentRNA_1_vs_LatentATAC_1 | 883 | 0.0075171799136821 | 0.0378095269418305 | 0.0211907374378003 | 0.0319614372971934 | 0.0049751243781094 | True | True |
| s4d8_GSE194122 | Proerythroblast | LatentRNA_1_vs_LatentATAC_1 | 800 | -0.0210159838497148 | 0.0176511733291766 | 0.0194271933413708 | 0.022743356229838 | 0.0895522388059701 | False | False |
| s4d8_GSE194122 | CD4+ T activated | LatentRNA_1_vs_LatentATAC_1 | 684 | 0.0908078209379734 | 0.1179739134839466 | 0.0236879256473283 | 0.0336597435310137 | 0.0049751243781094 | True | True |
| s4d8_GSE194122 | Transitional B | LatentRNA_1_vs_LatentATAC_1 | 630 | -0.0004472142659857 | 0.0303445118082611 | 0.0258411536220365 | 0.0347368545228775 | 0.0398009950248756 | True | False |
| s4d8_GSE194122 | CD14+ Mono | LatentRNA_1_vs_LatentATAC_1 | 458 | -0.028806058494327 | 0.011480541994778 | 0.0255992449236601 | 0.0310812873742349 | 0.2238805970149253 | False | False |
| s4d8_GSE194122 | NK | LatentRNA_1_vs_LatentATAC_1 | 421 | -0.0187951909749308 | 0.0162952210053481 | 0.027167017979711 | 0.043410049616935 | 0.1343283582089552 | False | False |
| s4d8_GSE194122 | Lymph prog | LatentRNA_1_vs_LatentATAC_1 | 327 | -0.0314132555125805 | 0.0085054034432981 | 0.0303285385598311 | 0.0384089997892232 | 0.3333333333333333 | False | False |
| s4d8_GSE194122 | B1 B | LatentRNA_1_vs_LatentATAC_1 | 326 | 0.0364436625563566 | 0.0835477337757299 | 0.0288375559795814 | 0.0390120485135299 | 0.0049751243781094 | True | True |
| s4d8_GSE194122 | CD4+ T naive | LatentRNA_1_vs_LatentATAC_1 | 299 | -0.0386565556167735 | 0.0025887276536364 | 0.0261375706084812 | 0.0444378086688512 | 0.4328358208955223 | False | False |

| donor | state | pair | n_cells | mi_obs_raw | delta_local | epsilon_95_local | epsilon_99_local | p_value_empirical | pass_local_95 | pass_local_99 |
|---|---|---|---|---|---|---|---|---|---|---|
| s4d8_GSE194122 | Erythroblast | LatentRNA_1_vs_LatentATAC_1 | 266 | -0.0307881462079873 | 0.0293400412423795 | 0.0302772238557765 | 0.0417903622709493 | 0.0646766169154228 | False | False |
| s4d9_GSE194122 | Naive CD20+ B | LatentRNA_1_vs_LatentATAC_1 | 897 | 0.0072188076204788 | 0.0327682203366176 | 0.0196991000996507 | 0.0232806733301078 | 0.0049751243781094 | True | True |
| s4d9_GSE194122 | CD8+ T | LatentRNA_1_vs_LatentATAC_1 | 805 | 0.0311708228981721 | 0.0597109867507919 | 0.0191076338660811 | 0.0314582684265158 | 0.0049751243781094 | True | True |
| s4d9_GSE194122 | Transitional B | LatentRNA_1_vs_LatentATAC_1 | 542 | -0.0086197558379179 | 0.0237098211587381 | 0.0244114285286753 | 0.0411112461469359 | 0.0646766169154228 | False | False |
| s4d9_GSE194122 | NK | LatentRNA_1_vs_LatentATAC_1 | 458 | -0.0133441556093849 | 0.0231001843984113 | 0.0272193722113332 | 0.0393303318179946 | 0.0945273631840796 | False | False |
| s4d9_GSE194122 | Lymph prog | LatentRNA_1_vs_LatentATAC_1 | 289 | 0.056263693296044 | 0.0967373117858802 | 0.030458795563973 | 0.043041595784506 | 0.0049751243781094 | True | True |
| s4d9_GSE194122 | CD8+ T naive | LatentRNA_1_vs_LatentATAC_1 | 219 | 0.0044326060913952 | 0.0556399139324645 | 0.0410703130620365 | 0.0525524320271515 | 0.0149253731343283 | True | False |
