## Supplementary material for "Hierarchical bounds on RNA–chromatin statistical dependence across cellular states in paired single-cell multiome data": data supplementary: SourceData_Fig3.pdf

### Source Data Fig 3: Compositional Decomposition

Derived from: RC3_R2_STRICT_BUNDLE (SHA256: 9730cd228d1d89a48444da777e41f1d77fddfa5912858bfa63d735b65bb1d07d)
Provenance: See Manifest_PRX_Life_FIXED.pdf

| donor | cmi_obs | cmi_null_mean | cmi_null_std | p_value_empirical | pass_cmi |
|---|---|---|---|---|---|
| s1d1_GSE194122 | -0.0206157787716945 | -0.0410602557855371 | 0.0038197147913128 | 0.0099009900990099 | True |
| s1d2_GSE194122 | 0.0068009474205201 | -0.0331616131890139 | 0.0042000219431539 | 0.0099009900990099 | True |
| s1d3_GSE194122 | -0.0118217064025444 | -0.0420087133499305 | 0.0054359296734216 | 0.0099009900990099 | True |
| s2d1_GSE194122 | -0.0376030965187449 | -0.0467655043191704 | 0.0049578449210446 | 0.0297029702970297 | True |
| s2d4_GSE194122 | -0.0117521822073421 | -0.0356911371699024 | 0.0040788675524838 | 0.0099009900990099 | True |
| s2d5_GSE194122 | 0.0057547512195032 | -0.0364503241048497 | 0.004804626851083 | 0.0099009900990099 | True |
| s3d10_GSE194122 | -0.0128754144751076 | -0.0407424483039502 | 0.0042037085878985 | 0.0099009900990099 | True |
| s3d3_GSE194122 | -0.0386372873960895 | -0.0474013914881924 | 0.0047489417205305 | 0.0693069306930693 | False |
| s3d6_GSE194122 | -0.0229496967938265 | -0.0645798412895708 | 0.0067278589080484 | 0.0099009900990099 | True |
| s3d7_GSE194122 | 0.0433768243205535 | -0.0591288825155365 | 0.0069582883737359 | 0.0099009900990099 | True |
| s4d1_GSE194122 | -0.0305721024011504 | -0.0409237758963744 | 0.0037873428374083 | 0.0099009900990099 | True |
| s4d8_GSE194122 | 0.0139613219461004 | -0.0295780148190816 | 0.0033939475082923 | 0.0099009900990099 | True |
| s4d9_GSE194122 | 0.0124799000826585 | -0.0428193065693546 | 0.0052029368415026 | 0.0099009900990099 | True |
