## Supplementary material for "Hierarchical bounds on RNA–chromatin statistical dependence across cellular states in paired single-cell multiome data": data supplementary: TableS2_Local_MI_and_Hotspot_Classification.pdf

### Table S2: Local MI Results

| donor | state | epsilon_95_local |
|---|---|---|
| s1d1_GSE194122 | CD14+ Mono | 0.0158727286880683 |
| s1d1_GSE194122 | CD4+ T activated | 0.0192002266537621 |
| s1d1_GSE194122 | Erythroblast | 0.0233996987746631 |
| s1d1_GSE194122 | CD8+ T | 0.0211285195289675 |
| s1d1_GSE194122 | NK | 0.0189700571802132 |
| s1d1_GSE194122 | CD4+ T naive | 0.025516067582895 |
| s1d1_GSE194122 | Normoblast | 0.0282953311740755 |
| s1d1_GSE194122 | Naive CD20+ B | 0.0286480134984948 |
| s1d1_GSE194122 | B1 B | 0.037357091523539 |
| s1d2_GSE194122 | CD8+ T | 0.0142714920178396 |
| s1d2_GSE194122 | CD4+ T naive | 0.0162650391145032 |
| s1d2_GSE194122 | NK | 0.0223664425295566 |
| s1d2_GSE194122 | CD4+ T activated | 0.0233754569444399 |
| s1d2_GSE194122 | CD14+ Mono | 0.021034397452366 |
| s1d2_GSE194122 | ILC | 0.0313186781573407 |
| s1d3_GSE194122 | CD8+ T | 0.020520051194339 |
| s1d3_GSE194122 | NK | 0.0222452293113885 |
| s1d3_GSE194122 | CD4+ T naive | 0.0300731162491661 |
| s1d3_GSE194122 | CD4+ T activated | 0.0296923707483993 |
| s1d3_GSE194122 | Erythroblast | 0.0283724059783983 |
| s1d3_GSE194122 | CD14+ Mono | 0.0331445953514979 |

| | | |
|---|---|---|
| s2d1_GSE194122 | CD14+ Mono | 0.0168380064651859 |
| s2d1_GSE194122 | Erythroblast | 0.0272035839594436 |
| s2d1_GSE194122 | CD4+ T activated | 0.0243502952039494 |
| s2d1_GSE194122 | Naive CD20+ B | 0.0288379001115744 |
| s2d1_GSE194122 | NK | 0.0284175637099313 |
| s2d1_GSE194122 | CD4+ T naive | 0.0295359938690249 |
| s2d1_GSE194122 | CD16+ Mono | 0.0379204071722893 |
| s2d4_GSE194122 | Erythroblast | 0.0159870415287966 |
| s2d4_GSE194122 | CD14+ Mono | 0.016470298369795 |
