## Supplementary material for "Hierarchical bounds on RNA–chromatin statistical dependence across cellular states in paired single-cell multiome data": data supplementary: TableS4_Counterexample_Donor_Report.pdf

### Table S4: Counterexample Report

| donor | state | pair | n_cells | delta_local | epsilon_95_local | p_value_empirical | failure_reason |
|---|---|---|---|---|---|---|---|
| s1d1_GSE194122 | CD14+ Mono | LatentRNA_1_vs_LatentATAC_1 | 921 | 0.0058963913064703 | 0.0158727286880683 | 0.3432835820895522 | High_PVal+Delta_Below_Eps |
| s1d1_GSE194122 | CD4+ T activated | LatentRNA_1_vs_LatentATAC_1 | 836 | 0.019656067270868 | 0.0192002266537621 | 0.0547263681592039 | High_PVal |
| s1d1_GSE194122 | CD8+ T | LatentRNA_1_vs_LatentATAC_1 | 725 | -0.0010395959275205 | 0.0211285195289675 | 0.5472636815920398 | High_PVal+Delta_Below_Eps |
| s1d1_GSE194122 | NK | LatentRNA_1_vs_LatentATAC_1 | 591 | -0.0140918661557583 | 0.0189700571802132 | 0.8507462686567164 | High_PVal+Delta_Below_Eps |
| s1d1_GSE194122 | CD4+ T naive | LatentRNA_1_vs_LatentATAC_1 | 543 | 0.0255412182247711 | 0.025516067582895 | 0.0547263681592039 | High_PVal |
| s1d1_GSE194122 | Normoblast | LatentRNA_1_vs_LatentATAC_1 | 387 | 0.0234706854412615 | 0.0282953311740755 | 0.1094527363184079 | High_PVal+Delta_Below_Eps |
| s1d1_GSE194122 | Naive CD20+ B | LatentRNA_1_vs_LatentATAC_1 | 386 | 0.0155335790983196 | 0.0286480134984948 | 0.1840796019900497 | High_PVal+Delta_Below_Eps |
| s1d1_GSE194122 | B1 B | LatentRNA_1_vs_LatentATAC_1 | 201 | 0.0020536478356324 | 0.037357091523539 | 0.4328358208955223 | High_PVal+Delta_Below_Eps |
| s1d2_GSE194122 | CD8+ T | LatentRNA_1_vs_LatentATAC_1 | 1632 | 0.003904727798646 | 0.0142714920178396 | 0.3283582089552239 | High_PVal+Delta_Below_Eps |
| s1d2_GSE194122 | CD14+ Mono | LatentRNA_1_vs_LatentATAC_1 | 704 | 0.01380658329429 | 0.021034397452366 | 0.1741293532338308 | High_PVal+Delta_Below_Eps |
| s1d3_GSE194122 | NK | LatentRNA_1_vs_LatentATAC_1 | 696 | -0.0006789267031411 | 0.0222452293113885 | 0.5074626865671642 | High_PVal+Delta_Below_Eps |
| s1d3_GSE194122 | Erythroblast | LatentRNA_1_vs_LatentATAC_1 | 374 | 0.0038197117414998 | 0.0283724059783983 | 0.3880597014925373 | High_PVal+Delta_Below_Eps |
| s1d3_GSE194122 | CD14+ Mono | LatentRNA_1_vs_LatentATAC_1 | 242 | 0.0073856241081241 | 0.0331445953514979 | 0.3681592039800995 | High_PVal+Delta_Below_Eps |
| s2d1_GSE194122 | CD14+ Mono | LatentRNA_1_vs_LatentATAC_1 | 1308 | 0.0037151988396521 | 0.0168380064651859 | 0.3582089552238806 | High_PVal+Delta_Below_Eps |
| s2d1_GSE194122 | Erythroblast | LatentRNA_1_vs_LatentATAC_1 | 478 | 0.0255026618561611 | 0.0272035839594436 | 0.0646766169154228 | High_PVal+Delta_Below_Eps |
| s2d1_GSE194122 | CD4+ T activated | LatentRNA_1_vs_LatentATAC_1 | 446 | 0.0083904441859539 | 0.0243502952039494 | 0.2786069651741293 | High_PVal+Delta_Below_Eps |
| s2d1_GSE194122 | Naive CD20+ B | LatentRNA_1_vs_LatentATAC_1 | 289 | -0.0333214219714787 | 0.0288379001115744 | 0.965174129353234 | High_PVal+Delta_Below_Eps |
| s2d1_GSE194122 | NK | LatentRNA_1_vs_LatentATAC_1 | 269 | -0.0133101183730311 | 0.0284175637099313 | 0.7860696517412935 | High_PVal+Delta_Below_Eps |
| s2d1_GSE194122 | CD4+ T naive | LatentRNA_1_vs_LatentATAC_1 | 254 | 0.0123183714177093 | 0.0295359938690249 | 0.2288557213930348 | High_PVal+Delta_Below_Eps |
| s2d4_GSE194122 | CD14+ Mono | LatentRNA_1_vs_LatentATAC_1 | 1140 | 0.0041543056633312 | 0.016470298369795 | 0.3880597014925373 | High_PVal+Delta_Below_Eps |
| s2d4_GSE194122 | Normoblast | LatentRNA_1_vs_LatentATAC_1 | 641 | 0.0160286743778012 | 0.0192604857409506 | 0.1144278606965174 | High_PVal+Delta_Below_Eps |

| | | | | | | | |
|---|---|---|---|---|---|---|---|
| s2d4_GSE194122 | Lymph prog | LatentRNA_1_vs_LatentATAC_1 | 286 | -0.005345893698069 | 0.0326298387702619 | 0.6019900497512438 | High_PVal+Delta_Below_Eps |
| s2d4_GSE194122 | CD8+ T | LatentRNA_1_vs_LatentATAC_1 | 230 | 0.0130650268628336 | 0.0439144122685486 | 0.2537313432835821 | High_PVal+Delta_Below_Eps |
| s2d4_GSE194122 | CD4+ T naive | LatentRNA_1_vs_LatentATAC_1 | 214 | -0.0069296815297635 | 0.0382132116325994 | 0.6069651741293532 | High_PVal+Delta_Below_Eps |
| s3d10_GSE194122 | CD14+ Mono | LatentRNA_1_vs_LatentATAC_1 | 1603 | 0.0019583067533737 | 0.0140714543368719 | 0.4378109452736318 | High_PVal+Delta_Below_Eps |
| s3d10_GSE194122 | NK | LatentRNA_1_vs_LatentATAC_1 | 545 | -0.0118490990911276 | 0.0240040797412603 | 0.8009950248756219 | High_PVal+Delta_Below_Eps |
| s3d10_GSE194122 | CD8+ T | LatentRNA_1_vs_LatentATAC_1 | 432 | 0.0065606775231474 | 0.0305004348175379 | 0.3034825870646766 | High_PVal+Delta_Below_Eps |
| s3d10_GSE194122 | pDC | LatentRNA_1_vs_LatentATAC_1 | 427 | -0.0024848807891941 | 0.0297448750777942 | 0.5671641791044776 | High_PVal+Delta_Below_Eps |
| s3d10_GSE194122 | Naive CD20+ B | LatentRNA_1_vs_LatentATAC_1 | 421 | -0.007439914874505 | 0.0330074141553054 | 0.6417910447761194 | High_PVal+Delta_Below_Eps |
| s3d10_GSE194122 | CD8+ T naive | LatentRNA_1_vs_LatentATAC_1 | 366 | -0.0029503599190805 | 0.0301971137507653 | 0.5771144278606966 | High_PVal+Delta_Below_Eps |
| s3d10_GSE194122 | Transitional B | LatentRNA_1_vs_LatentATAC_1 | 359 | -0.0028157933437914 | 0.0292016015839593 | 0.5323383084577115 | High_PVal+Delta_Below_Eps |
| s3d10_GSE194122 | HSC | LatentRNA_1_vs_LatentATAC_1 | 291 | 0.0040886227490236 | 0.0372790467405235 | 0.417910447761194 | High_PVal+Delta_Below_Eps |
| s3d10_GSE194122 | CD4+ T activated | LatentRNA_1_vs_LatentATAC_1 | 207 | -0.007090671635488 | 0.0359309344566727 | 0.6218905472636815 | High_PVal+Delta_Below_Eps |
| s3d3_GSE194122 | CD14+ Mono | LatentRNA_1_vs_LatentATAC_1 | 855 | -0.0022477226907012 | 0.02139053547684 | 0.572139303482587 | High_PVal+Delta_Below_Eps |
| s3d3_GSE194122 | NK | LatentRNA_1_vs_LatentATAC_1 | 477 | -0.0084127036412332 | 0.0301215544180891 | 0.6716417910447762 | High_PVal+Delta_Below_Eps |
| s3d3_GSE194122 | Naive CD20+ B | LatentRNA_1_vs_LatentATAC_1 | 393 | -0.0147654132738992 | 0.0340972482584336 | 0.7810945273631841 | High_PVal+Delta_Below_Eps |
| s3d3_GSE194122 | CD8+ T | LatentRNA_1_vs_LatentATAC_1 | 378 | -0.0054165105659162 | 0.0277978537425241 | 0.6218905472636815 | High_PVal+Delta_Below_Eps |
| s3d3_GSE194122 | CD8+ T naive | LatentRNA_1_vs_LatentATAC_1 | 228 | -0.0045863285701171 | 0.0394079037569685 | 0.5920398009950248 | High_PVal+Delta_Below_Eps |
| s3d3_GSE194122 | CD4+ T activated | LatentRNA_1_vs_LatentATAC_1 | 228 | -0.0103919518117341 | 0.0364396576837594 | 0.6318407960199005 | High_PVal+Delta_Below_Eps |
| s3d3_GSE194122 | B1 B | LatentRNA_1_vs_LatentATAC_1 | 217 | -0.003669224356198 | 0.0300236143799981 | 0.6019900497512438 | High_PVal+Delta_Below_Eps |
| s3d6_GSE194122 | CD8+ T | LatentRNA_1_vs_LatentATAC_1 | 488 | 0.0137868849697718 | 0.0236997708154864 | 0.1840796019900497 | High_PVal+Delta_Below_Eps |
| s3d6_GSE194122 | CD14+ Mono | LatentRNA_1_vs_LatentATAC_1 | 208 | 0.0203841498843726 | 0.0308777783252361 | 0.1293532338308457 | High_PVal+Delta_Below_Eps |
| s3d7_GSE194122 | CD14+ Mono | LatentRNA_1_vs_LatentATAC_1 | 674 | 0.021971816353744 | 0.0246131797381476 | 0.0746268656716417 | High_PVal+Delta_Below_Eps |
| s3d7_GSE194122 | NK | LatentRNA_1_vs_LatentATAC_1 | 256 | -0.0019834264323887 | 0.0321942494007848 | 0.4975124378109453 | High_PVal+Delta_Below_Eps |
| s4d1_GSE194122 | NK | LatentRNA_1_vs_LatentATAC_1 | 1124 | 0.0119497450272271 | 0.0174632624501075 | 0.1243781094527363 | High_PVal+Delta_Below_Eps |
| s4d1_GSE194122 | CD14+ Mono | LatentRNA_1_vs_LatentATAC_1 | 1014 | 0.0109258908429954 | 0.0224654292692214 | 0.1791044776119403 | High_PVal+Delta_Below_Eps |

| | | | | | | | |
|---|---|---|---|---|---|---|---|
| s4d1_GSE194122 | CD8+ T | LatentRNA_1_vs_LatentATAC_1 | 759 | -0.0070245076569303 | 0.0204785357905523 | 0.7213930348258707 | High_PVal+Delta_Below_Eps |
| s4d1_GSE194122 | Transitional B | LatentRNA_1_vs_LatentATAC_1 | 695 | 0.0083881629297999 | 0.0226299282013555 | 0.2288557213930348 | High_PVal+Delta_Below_Eps |
| s4d1_GSE194122 | Naive CD20+ B | LatentRNA_1_vs_LatentATAC_1 | 626 | 0.0110646782320682 | 0.0255875662949063 | 0.2388059701492537 | High_PVal+Delta_Below_Eps |
| s4d1_GSE194122 | CD4+ T activated | LatentRNA_1_vs_LatentATAC_1 | 619 | 0.0088312587335115 | 0.0256646125083805 | 0.2537313432835821 | High_PVal+Delta_Below_Eps |
| s4d1_GSE194122 | Erythroblast | LatentRNA_1_vs_LatentATAC_1 | 474 | -0.0060348905149936 | 0.0312297663923003 | 0.5870646766169154 | High_PVal+Delta_Below_Eps |
| s4d1_GSE194122 | B1 B | LatentRNA_1_vs_LatentATAC_1 | 420 | -0.0093751808541211 | 0.0272034330749589 | 0.7263681592039801 | High_PVal+Delta_Below_Eps |
| s4d1_GSE194122 | Proerythroblast | LatentRNA_1_vs_LatentATAC_1 | 322 | 0.0259382533452326 | 0.0293981389390133 | 0.0945273631840796 | High_PVal+Delta_Below_Eps |
| s4d1_GSE194122 | Lymph prog | LatentRNA_1_vs_LatentATAC_1 | 291 | 0.0171582476348314 | 0.0277924510936539 | 0.1393034825870646 | High_PVal+Delta_Below_Eps |
| s4d1_GSE194122 | CD16+ Mono | LatentRNA_1_vs_LatentATAC_1 | 251 | -0.0045178449830693 | 0.0347406117371169 | 0.5920398009950248 | High_PVal+Delta_Below_Eps |
| s4d1_GSE194122 | MK/E prog | LatentRNA_1_vs_LatentATAC_1 | 201 | 0.0231814389937155 | 0.0317541287295776 | 0.1691542288557214 | High_PVal+Delta_Below_Eps |
| s4d8_GSE194122 | Proerythroblast | LatentRNA_1_vs_LatentATAC_1 | 800 | 0.0176511733291766 | 0.0194271933413708 | 0.0895522388059701 | High_PVal+Delta_Below_Eps |
| s4d8_GSE194122 | CD14+ Mono | LatentRNA_1_vs_LatentATAC_1 | 458 | 0.011480541994778 | 0.0255992449236601 | 0.2238805970149253 | High_PVal+Delta_Below_Eps |
| s4d8_GSE194122 | NK | LatentRNA_1_vs_LatentATAC_1 | 421 | 0.0162952210053481 | 0.027167017979711 | 0.1343283582089552 | High_PVal+Delta_Below_Eps |
| s4d8_GSE194122 | Lymph prog | LatentRNA_1_vs_LatentATAC_1 | 327 | 0.0085054034432981 | 0.0303285385598311 | 0.3333333333333333 | High_PVal+Delta_Below_Eps |
| s4d8_GSE194122 | CD4+ T naive | LatentRNA_1_vs_LatentATAC_1 | 299 | 0.0025887276536364 | 0.0261375706084812 | 0.4328358208955223 | High_PVal+Delta_Below_Eps |
| s4d8_GSE194122 | Erythroblast | LatentRNA_1_vs_LatentATAC_1 | 266 | 0.0293400412423795 | 0.0302772238557765 | 0.0646766169154228 | High_PVal+Delta_Below_Eps |
| s4d9_GSE194122 | Transitional B | LatentRNA_1_vs_LatentATAC_1 | 542 | 0.0237098211587381 | 0.0244114285286753 | 0.0646766169154228 | High_PVal+Delta_Below_Eps |
| s4d9_GSE194122 | NK | LatentRNA_1_vs_LatentATAC_1 | 458 | 0.0231001843984113 | 0.0272193722113332 | 0.0945273631840796 | High_PVal+Delta_Below_Eps |
