## Supplementary figures and images for "Hierarchical bounds on RNA–chromatin statistical dependence across cellular states in paired single-cell multiome data"

### Fig1_Global_MI_vs_Nulls.pdf

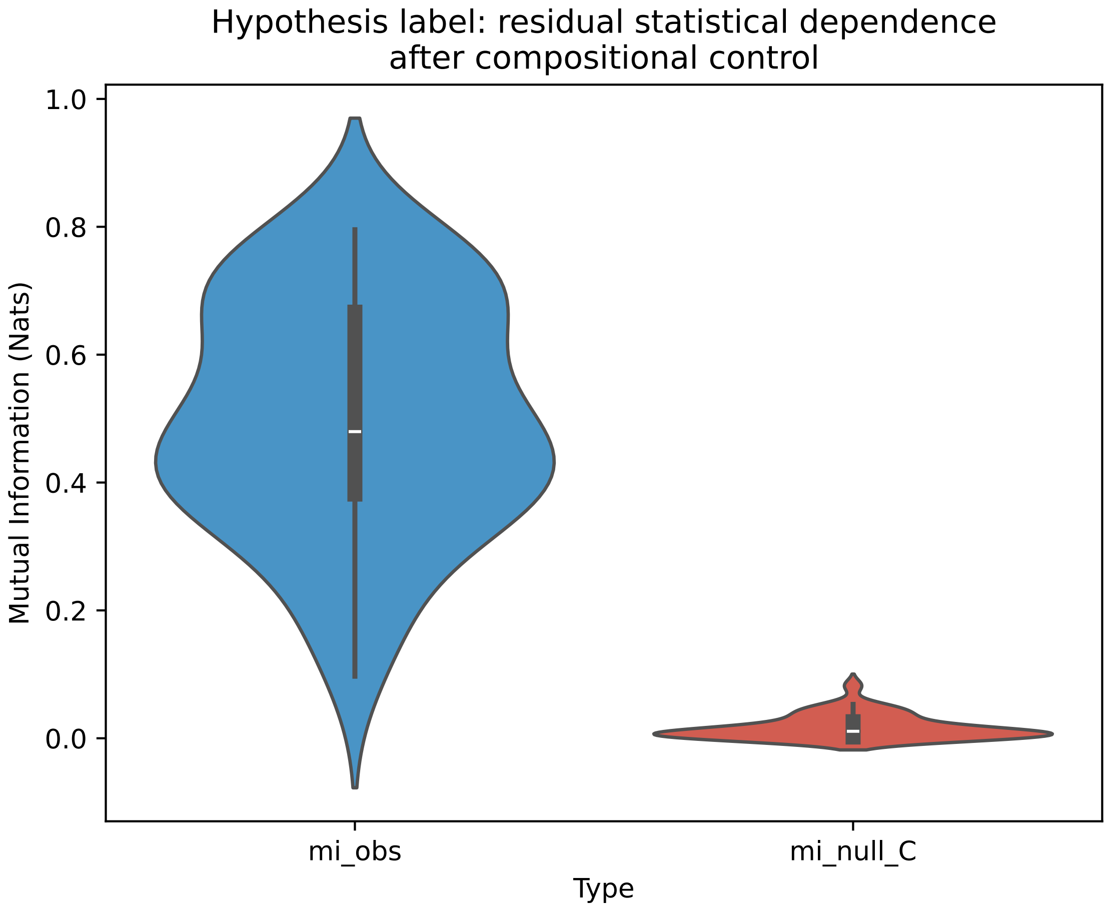

### Fig2_Local_MI_State_Distributions.pdf

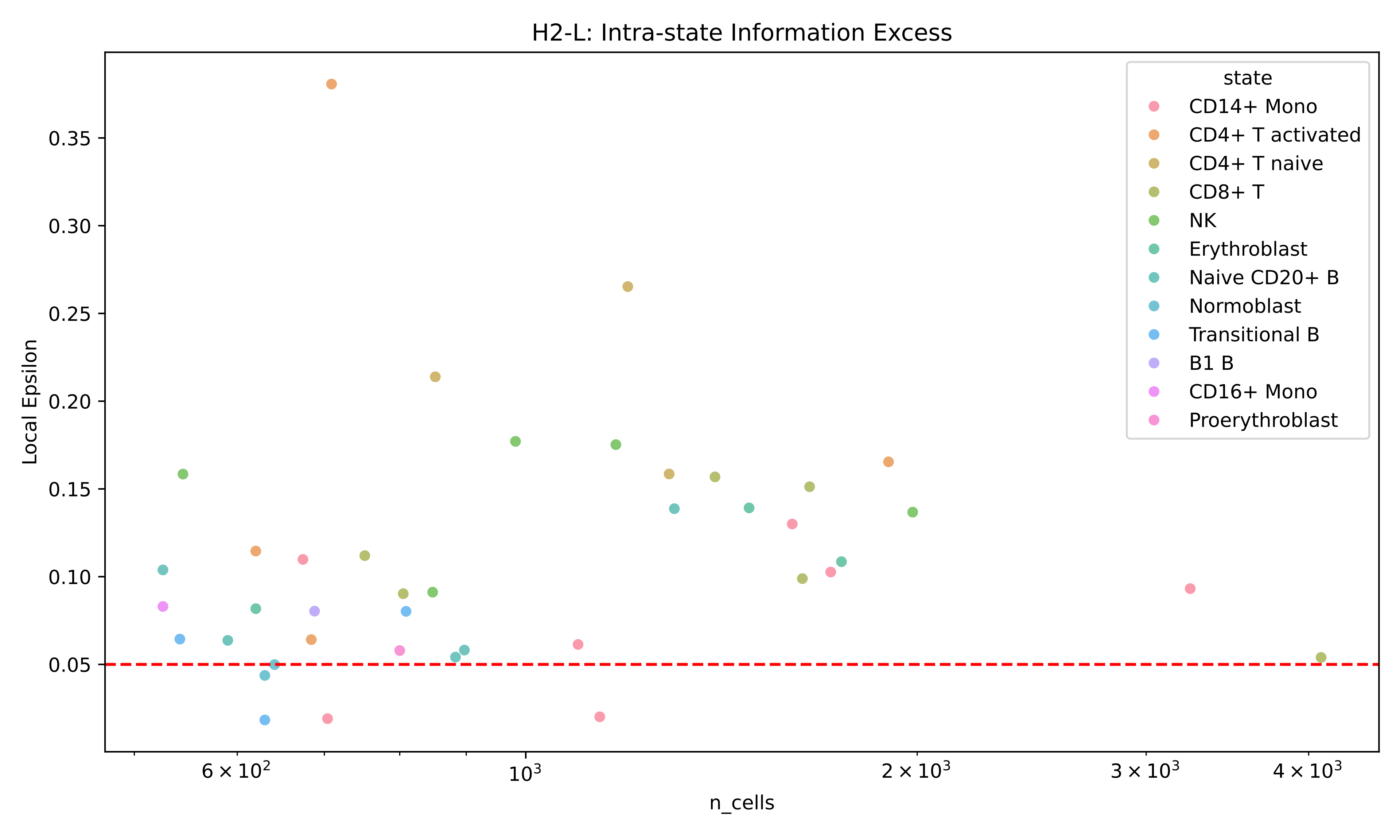

### Fig3_Compositional_Decomposition_rho.pdf

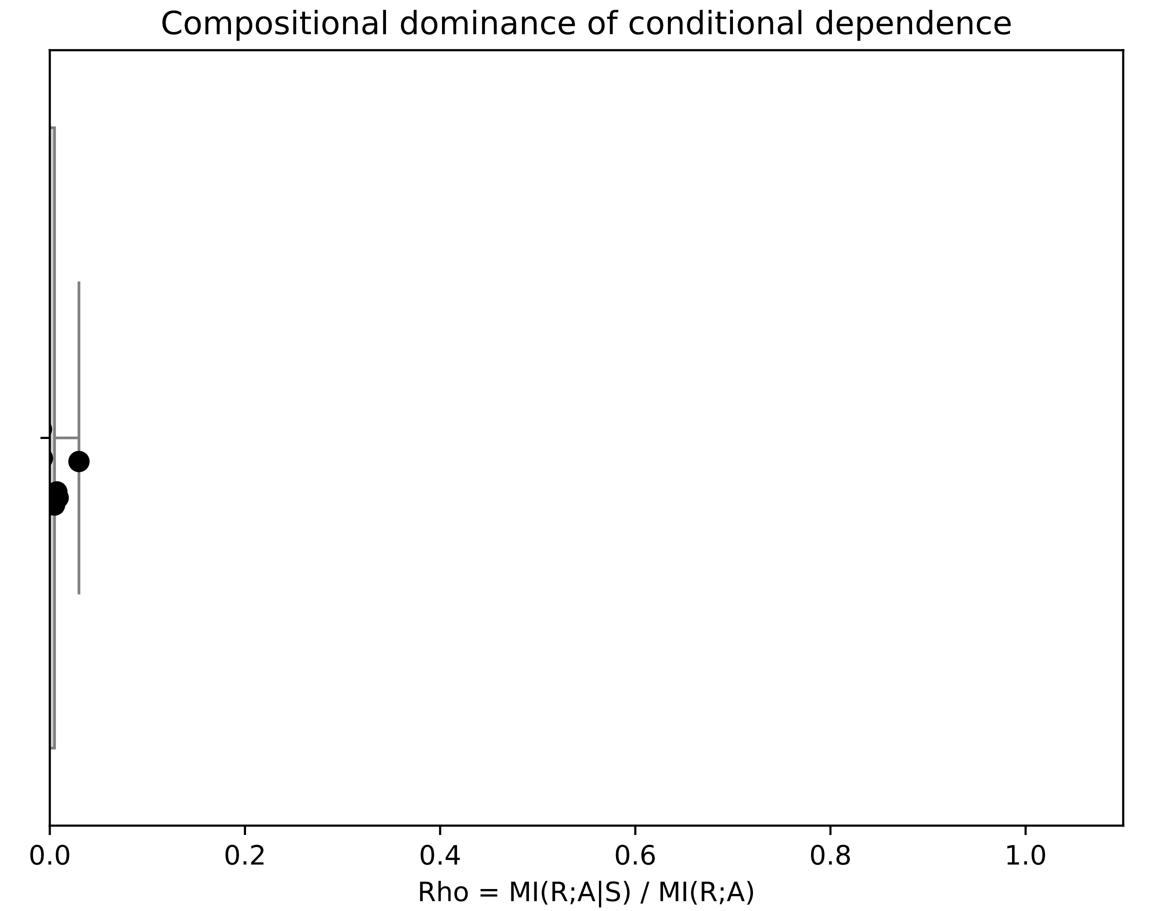

### Fig4_Conditional_MI_Orthogonal_Test.pdf

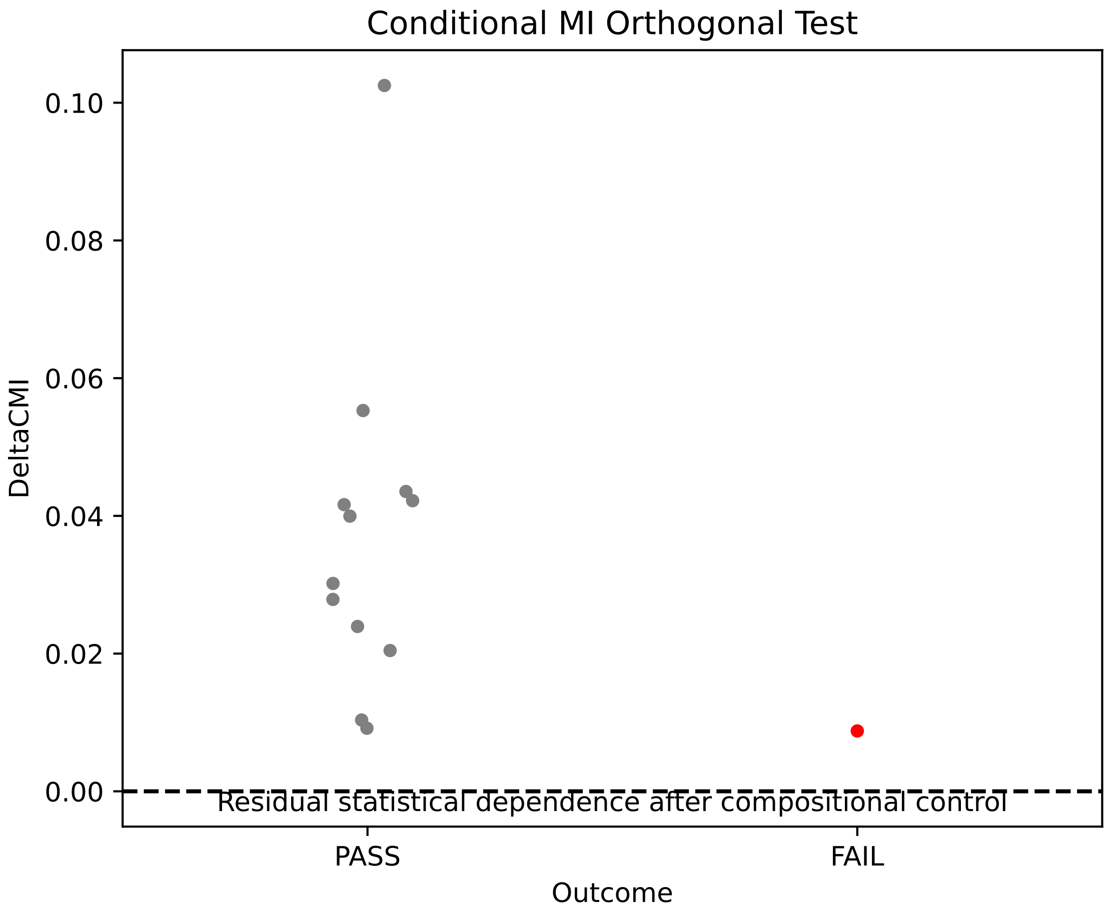

### Fig5_Hierarchical_Channel_Summary.pdf

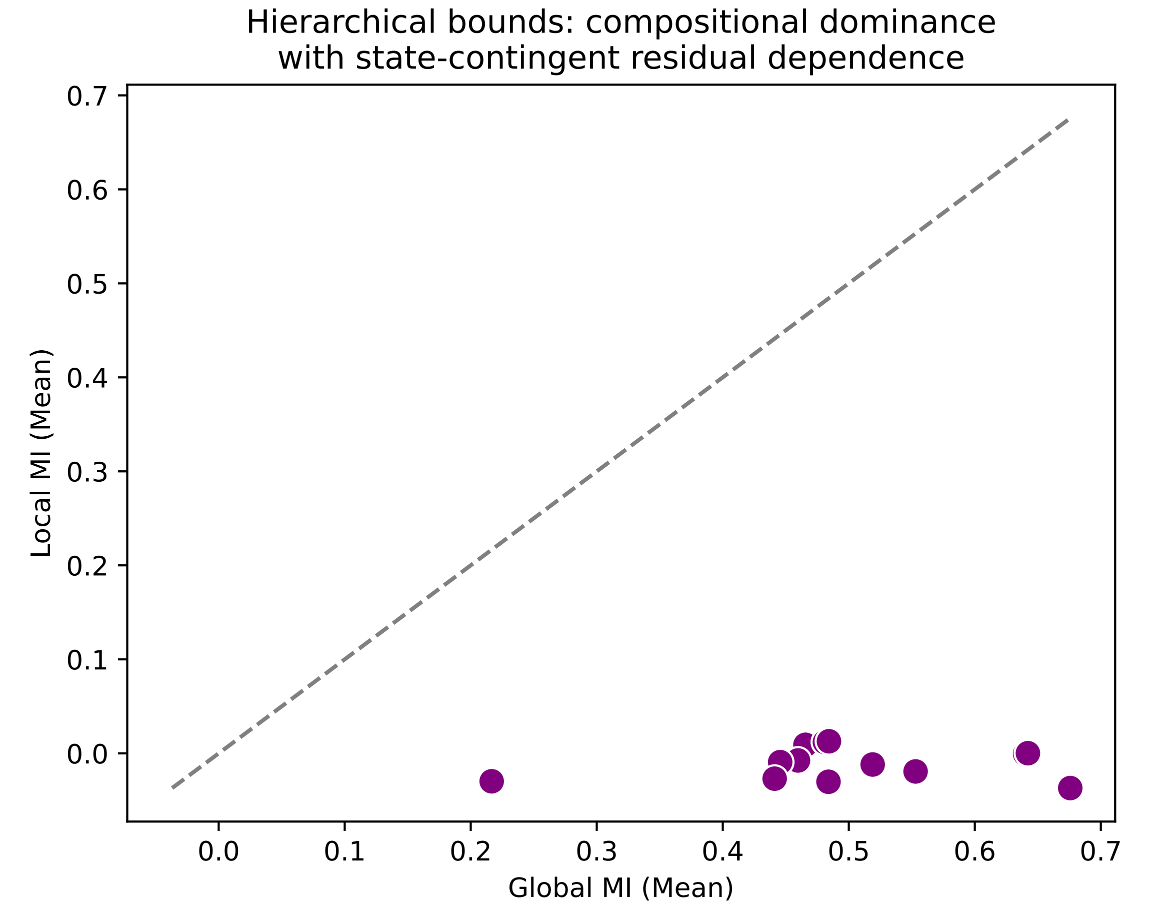

### SuppFig2_Epsilon_Calibration_Noise_Floor.pdf

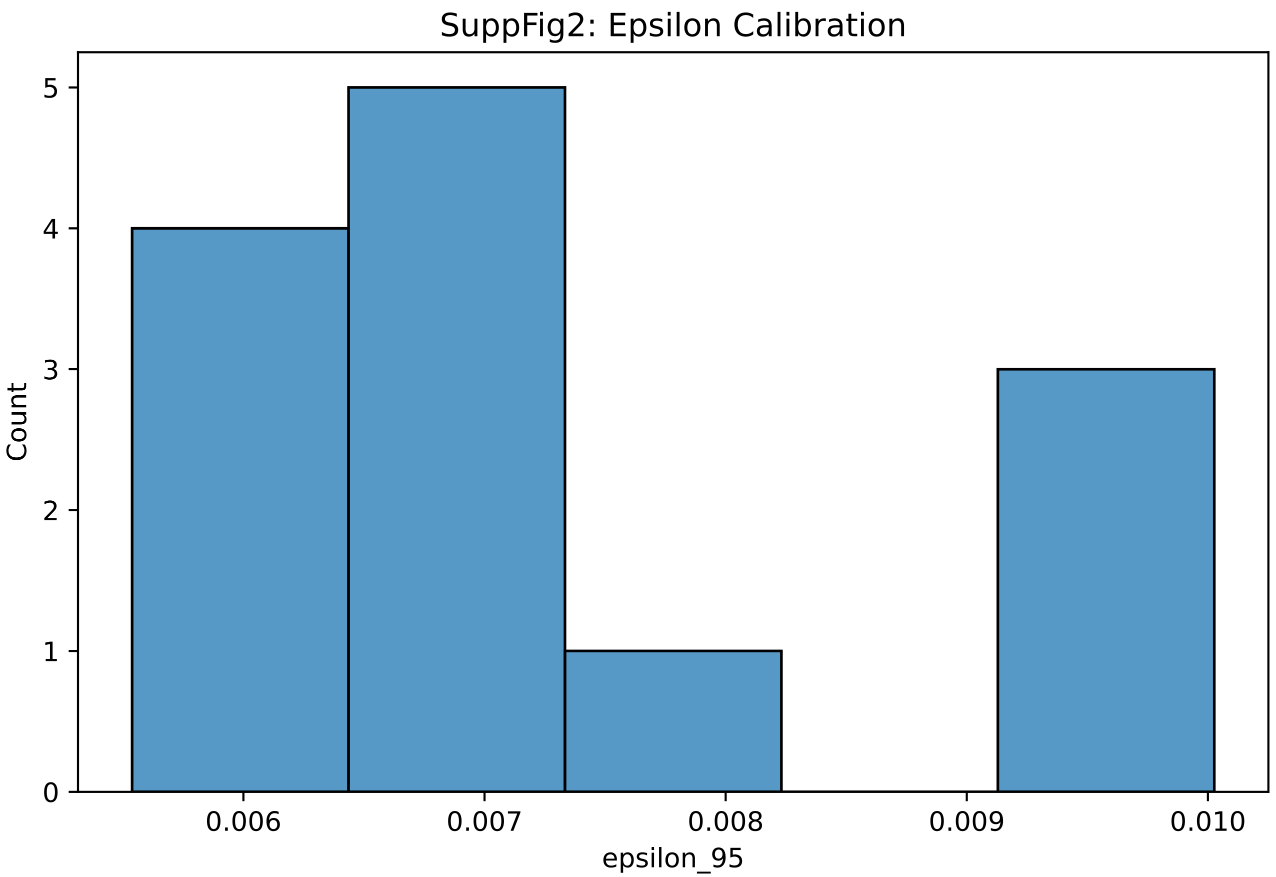

### SuppFig3_Permutation_Stability_Checks.pdf

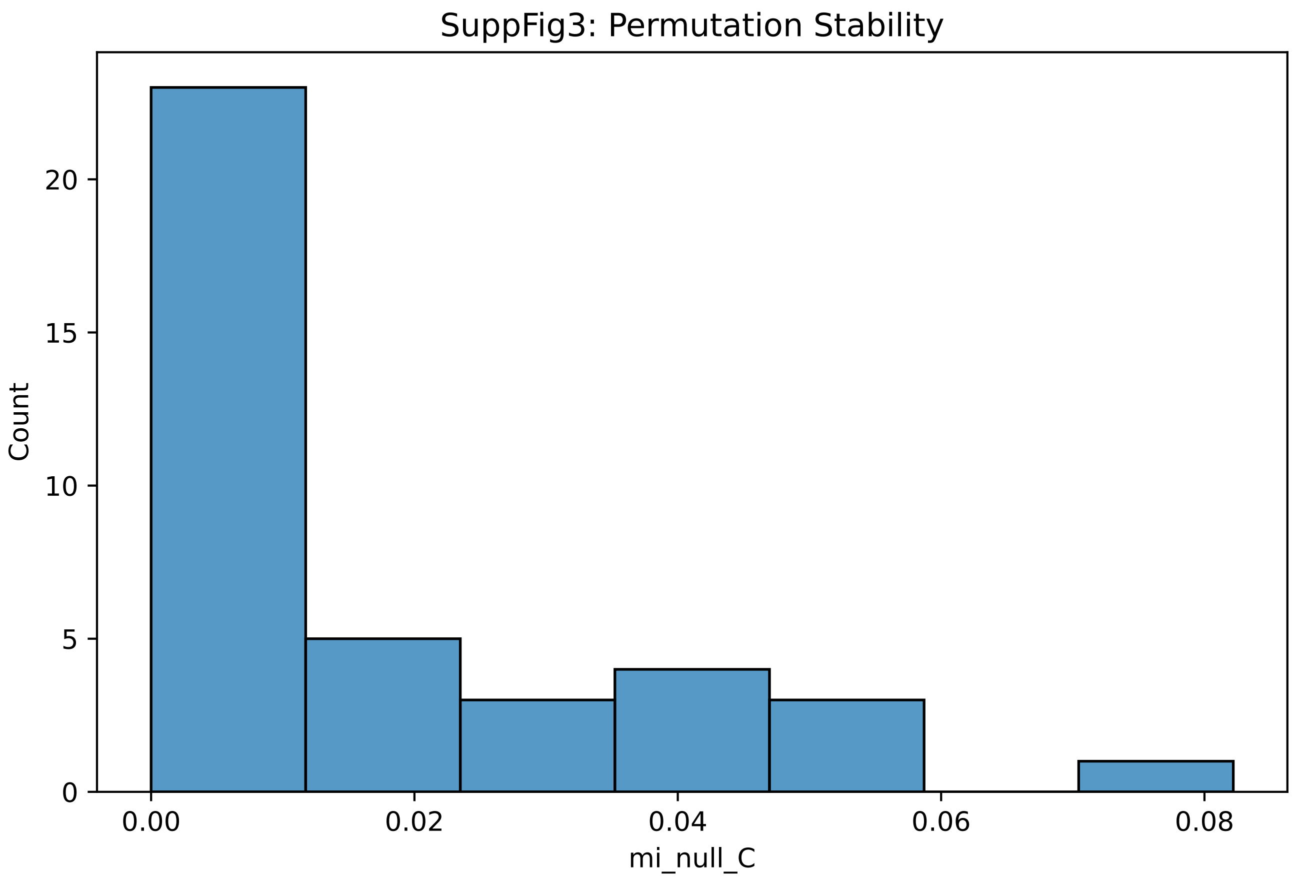

### SuppFig4_Donor_Heterogeneity.pdf

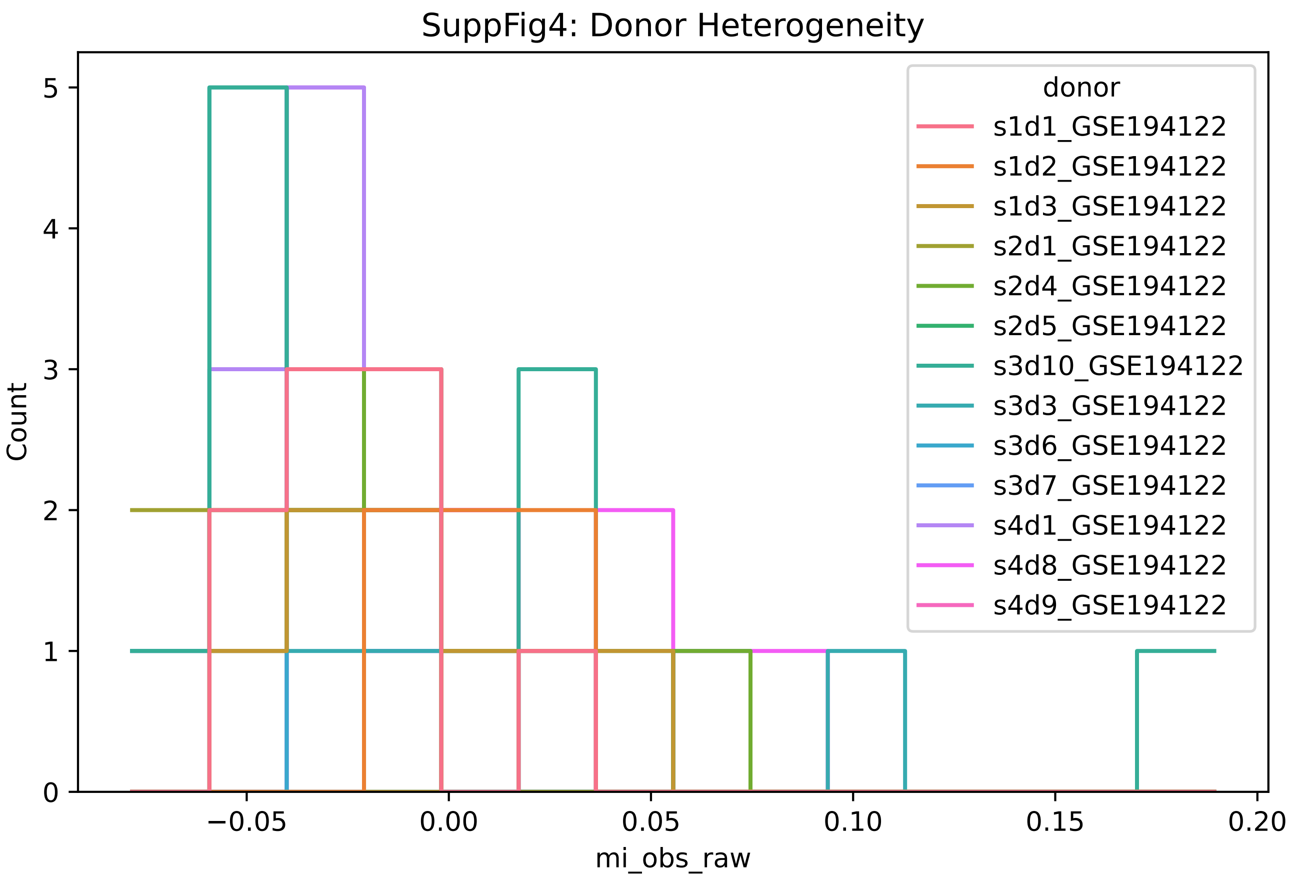

### SuppFig5_Estimator_Bias_Sensitivity.pdf

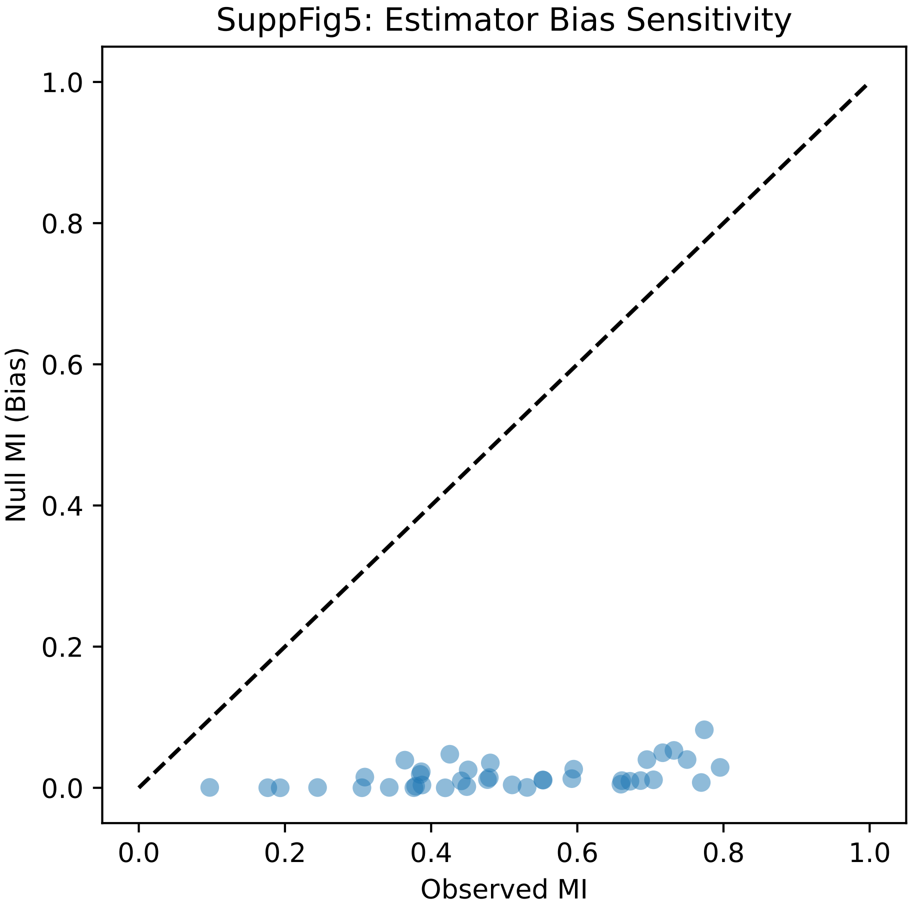

### SuppFig6_Parameter_Robustness.pdf

#### SuppFig6: Parameter Robustness

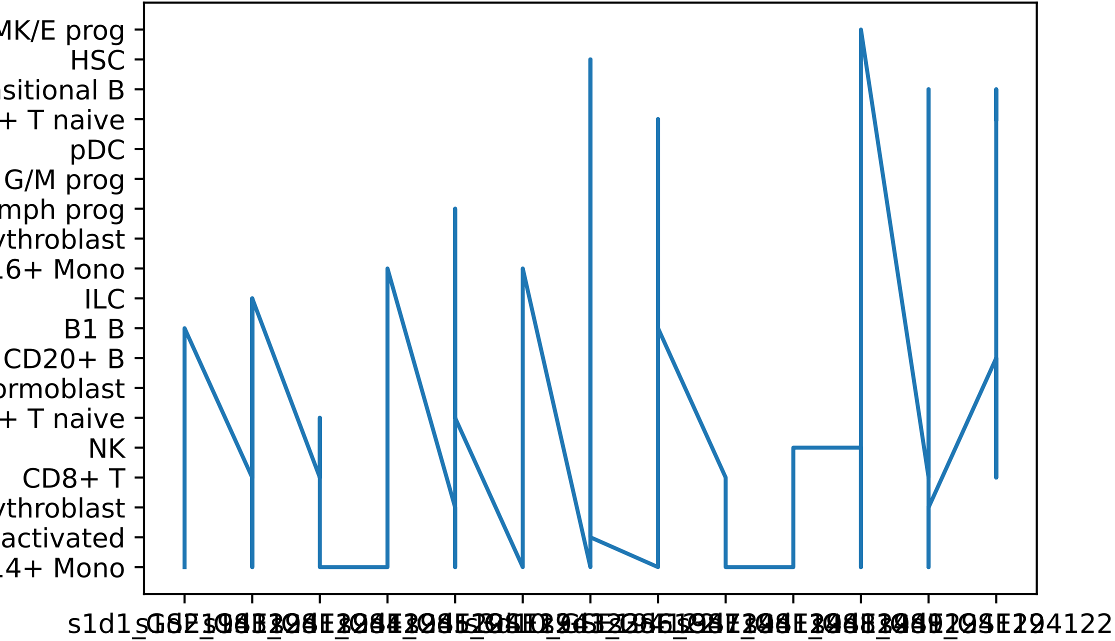

### SuppFig7_Noise_Floor_and_Epsilon_Validation.pdf

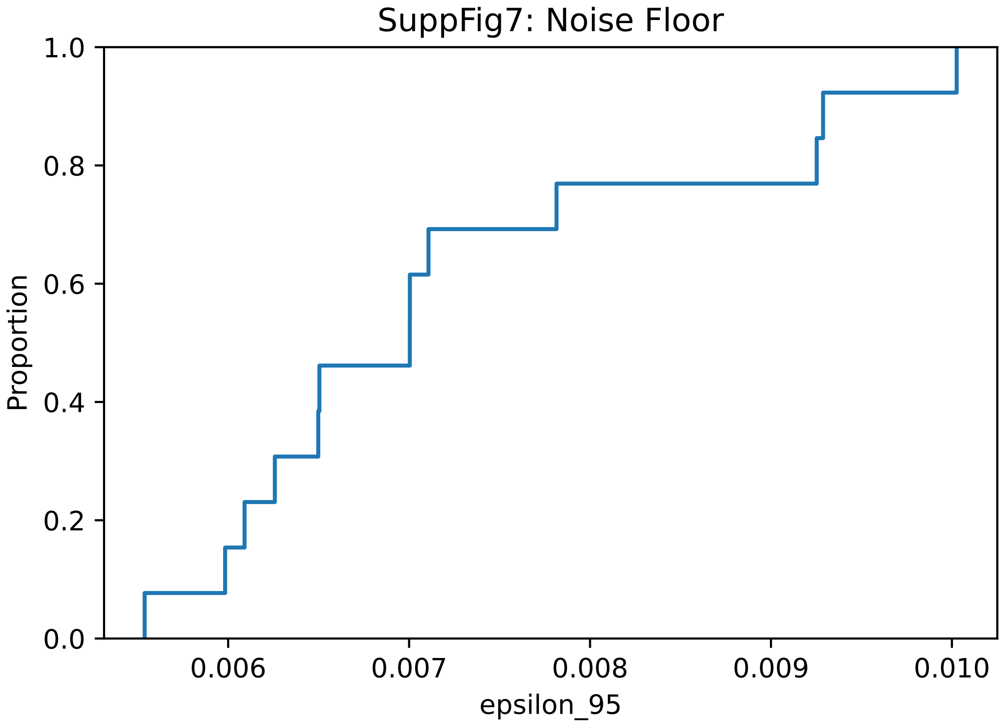
